# MT-LLE: Multi-task locally linear embedding for interpretable disease modeling from longitudinal omics data

**DOI:** 10.64898/2026.08.24.746083

**Authors:** Sundous Hussein, Iain R. Konigsberg, Katerina J. Kechris, Russell P. Bowler, Farnoush Banaei-Kashani

## Abstract

Constructing interpretable disease models from longitudinal omics data is a central challenge in precision medicine. The goal is a low-dimensional representation in which a patient’s position encodes their molecular state and clinical severity, and along which disease progression can be read directly. Existing dimensionality reduction methods (e.g., UMAP, Variational Autoencoders) fall short of this goal: they optimize a single generic objective and are blind to clinical labels and to the temporal ordering of measurements. Consequently, trajectory inference is typically applied after the fact to an embedding that was never optimized to reveal progression, decoupling the representation from disease dynamics.

Manifold learning offers a natural route to such representations, and we build on Locally Linear Embedding (LLE) to preserve the local geometry of the omics data (i.e., keeping molecularly similar patients close together in the low-dimensional space). Geometry alone, however, yields a space that is faithful to molecular similarity yet uninformative about clinical severity and progression. We therefore recast the problem as multi-task learning: MT-LLE jointly optimizes five objectives: geometric reconstruction, supervised organization by clinical stage, embedding and phenotype forecasting, and clustering. Because naively combining such heterogeneous objectives induces gradient conflicts that distort the molecular geometry, an embedded reinforcement learning agent dynamically schedules their weights during training, establishing global geometry before refining clinical boundaries.

Across two independent Chronic Obstructive Pulmonary Disease (COPD) cohorts (SPIROMICS and COPDGene), MT-LLE deliberately relaxes exact geometric reconstruction, by a modest margin, in exchange for substantial gains in clinical structure. On held-out patients, a linear model reads disease severity (GOLD stage, 0–4) from the MT-LLE embedding 35–40% more accurately than from standard dimensionality reduction (0.53 vs. 0.38 F1-Macro). The gap is starker for progression: forecasting a patient’s next-visit severity from their trajectory reaches 0.38 F1-Macro, while unsupervised baselines sit near zero (0.06–0.09 F1-Macro), a temporal signal those methods fail to capture. To test whether the reinforcement learning agent earns its place, we compared it against a fixed schedule that imposes the same ordering of objectives but cannot adapt during training; the learned agent outperforms it by 13–20% across clinical metrics, showing the gains come from adapting the weights to how training unfolds, not merely from ordering the objectives correctly, and at no cost to geometric fidelity. Beyond these quantitative gains, the manifold supports complementary analyses that surface structure invisible to standard staging: static phenotyping isolates subjects with active molecular pathology despite preserved lung function; trajectory inference maps two mechanistically distinct progression axes (inflammatory fibrosis and pan-immune activation); and kinematic analysis of each patient’s speed and acceleration identifies subjects whose molecular trajectories accelerate ahead of detectable spirometric decline.

## Introduction

Understanding heterogeneous diseases remains one of the foremost challenges in biomedical research. Complex pathologies, from neurodegeneration to cancer, exhibit extensive molecular and phenotypic heterogeneity that evolves over time, and longitudinal omics datasets now offer an unprecedented opportunity to model these dynamics. Translating high-dimensional molecular profiles into an interpretable disease model, however, remains an open problem. The goal is to move beyond static snapshots and construct a disease manifold: a continuous, low-dimensional map in which each patient’s position encodes their molecular state and clinical severity, nearby patients share similar molecular profiles, and disease progression can be read directly from movement across the map.

Current computational approaches fall short of this goal. Standard dimensionality reduction techniques (e.g., Uniform Manifold Approximation and Projection (UMAP) [1], Variational Autoencoders (VAEs) [2]), optimize a single, generic objective such as preserving local neighborhood structure, which leaves them blind to clinical phenotypes and to the temporal ordering of measurements. As a result, the distance between two patients in the learned space does not reflect their biological or clinical similarity. This limitation compounds when progression is the target: because the embedding is fixed before any temporal structure is considered, trajectory inference is applied post-hoc, as a separate step on a space that was never optimized to reveal progression. Representation learning and trajectory modeling are thereby decoupled, and the resulting manifolds are not shaped to expose coherent progression pathways or distinct molecular subtypes — a continuity that the discrete, rigid categories of traditional clinical staging also fail to capture.

To address this, we present **Multi-Task Locally Linear Embedding (MT-LLE)**, a framework for constructing interpretable disease models from longitudinal omics data. A useful disease model must be locally faithful to the omics geometry, globally organized by clinical outcome, and temporally coherent enough to support forecasting. Combining these objectives is a genuine optimization problem: naively training them together produces negative transfer, in which competing tasks degrade rather than reinforce the shared representation. MT-LLE resolves this with a reinforcement learning (RL) agent that dynamically balances the geometric, supervised, and temporal objectives during training, establishing global geometry before refining clinical boundaries rather than forcing all objectives to compete from the outset.

Our contributions are as follows:

- **A multi-task manifold framework for interpretable disease representations**. We integrate five objectives into a single latent space that is simultaneously faithful to the omics geometry, organized by clinical severity, and temporally coherent: geometric reconstruction, supervised contrastive organization, embedding forecasting, phenotype forecasting, and clustering. Unlike prior multi-task omics models that treat patients as cross-sectional snapshots, and unlike trajectory-inference methods that infer dynamics only after the embedding is fixed, MT-LLE learns a single representation jointly shaped by geometry, clinical structure, and temporal dynamics for repeated-measures longitudinal cohorts.
- **Dynamic task weighting via reinforcement learning**. We present, to our knowledge, the first RL agent formulated to balance heterogeneous, non-metricized objectives for manifold learning. Whereas gradient-based methods such as GradNorm [3] and PCGrad [4], and prior RL weighting schemes [5] assume homogeneous supervised tasks, our agent rewards improvement in the manifold’s overall structure and discovers a staged schedule (geometric foundation, structural wiring, then supervised fine-tuning) that static and gradient-norm approaches cannot replicate.
- **Kinematic phenotyping and mechanistic discovery**. We characterize each patient not by their static position but by their speed and acceleration through the learned manifold, surfacing dynamic phenotypes invisible to static profiling and discrete staging. Applied to two independent Chronic Obstructive Pulmonary Disease (COPD) cohorts, MT-LLE resolves biological structure that standard clinical staging misses: subjects with active molecular pathology despite preserved lung function, a pro-resolving lipid-mediator phenotype, two mechanistically distinct progression axes (inflammatory fibrosis vs. pan-immune activation), and subjects whose molecular trajectories accelerate ahead of detectable spirometric decline.

## Related work

### Representation learning and trajectory inference for omics data

The integration of heterogeneous omics layers is foundational for precision medicine. Early approaches focused on unsupervised latent variable models, such as Multi-Omics Factor Analysis (MOFA) [6] and iCluster [7], which disentangle shared sources of variation. Network-based methods, such as Similarity Network Fusion (SNF) [8] and Sparse Multiple Canonical Correlation Network (SmCCNet) [9], construct patient similarity graphs or feature correlation networks to capture global topologies. More recently, Graph Neural Networks (GNNs) like MOGONET [10] have demonstrated success in learning supervised embeddings for classification. However, these approaches typically suffer from two limitations. First, network aggregation often obscures local, subject-specific heterogeneity by constructing global topologies based on population-level statistics. Second, standard nonlinear dimensionality reduction techniques used to visualize these data (e.g., t-SNE, UMAP, and VAEs) are inherently single-objective. They optimize for a specific geometric property, such as local neighborhood preservation or global variance, often ignoring clinical labels or temporal continuity. This yields manifolds blind to clinical labels and temporal ordering, where the distance between subjects may not reflect biological or clinical similarity.

Recent work has sought to incorporate supervision into the embedding process. Label-guided extensions of UMAP [1] and parametric variants [11] enable some degree of phenotype-aware organization, while contrastive representation learning methods such as SimCLR [12] and Supervised Contrastive Learning [13] have been adapted for omics data to enforce class-level separation in latent space. However, these approaches remain fundamentally single-objective: they optimize for either geometric fidelity or supervised separability, but not both simultaneously, and critically, none are designed to encode temporal progression. MT-LLE extends this line of work by integrating supervised contrastive learning as one of five jointly optimized objectives, explicitly balancing it against geometric reconstruction and temporal forecasting to prevent the manifold collapse that occurs when supervision is applied in isolation.

To address the temporal aspect, manifold learning methods have been applied to align heterogeneous omics datasets, as seen in ManiNetCluster [14] for gene networks and MMDMA [15] for single-cell modalities. While these approaches excel at integrating data types, they are not designed to explicitly model the smooth, continuous progression inherent in longitudinal data. In the single-cell domain, RNA velocity methods [16, 17] infer directional progression from splicing kinetics, and trajectory inference tools such as Monocle [18] and Slingshot [19] have established pseudotime as a standard framework for ordering cells along progression axes. For bulk longitudinal cohorts, latent ODE frameworks [20, 21] learn continuous-time latent trajectories by parameterizing the derivative of the latent state, enabling flexible modeling of irregularly sampled time points. These methods share two limitations relative to our setting. First, trajectory inference is applied post-hoc to a pre-computed embedding that was not optimized with trajectory discovery in mind, fundamentally decoupling representation learning from the goal of trajectory discovery. Second, the single-cell approaches order cells within a static population along a pseudotemporal axis; they do not model the repeated longitudinal measurement of the same subjects over calendar time, which is the regime MT-LLE targets. A patient’s trajectory in our setting is an observed sequence of real visits, not an inferred ordering of a snapshot population. Our framework bridges this gap by learning a manifold that is not only structured by the geometry of the omics data but is explicitly regularized by temporal forecasting objectives, so that the embedding and its progression structure are shaped together rather than learned in sequence.

### Multi-task learning for omics data

Multi-task learning (MTL) has emerged as a powerful paradigm for learning from omics data by learning a shared representation across several related tasks. Frameworks like OmiEmbed [22] and scMoMtF [23] have demonstrated that jointly training for tasks such as tumor classification, survival prediction, and data reconstruction yields more informative embeddings and improves predictive performance compared to single-task models. Similarly, attention-based models like MOMA [24] integrate multiple omics layers to identify candidate drug targets by learning to prioritize the most informative data types. However, these methods suffer from two critical limitations. First, they typically rely on standard deep neural network architectures (e.g., autoencoders) and focus on a set of homogeneous supervised prediction or reconstruction tasks. Second, and most importantly, these frameworks are designed for static cross-sectional data, treating patient profiles as fixed snapshots. They do not account for the temporal dependencies inherent in longitudinal studies. While prior multi-task frameworks for omics data have focused on cross-sectional settings and homogeneous supervised objectives, **MT-LLE** is, to our knowledge, the first to simultaneously optimize geometric manifold fidelity, supervised phenotype organization, and temporal trajectory forecasting in a unified framework for longitudinal omics data.

### Reinforcement learning for dynamic task weighting

A key challenge in MTL is balancing the influence of each task during training. Gradient-based conflict resolution methods have been proposed to address this directly: GradNorm [3] dynamically adjusts loss weights to equalize gradient norms across tasks, preventing any single objective from dominating the shared representation; PCGrad [4] projects task gradients to remove conflicting components before each update; and MGDA [25] frames MTL as a multi-objective optimization problem seeking Pareto-optimal gradient directions. More recent formulations, including CAGrad [26] and Nash-MTL [27], have proposed game-theoretic solutions to the same problem. However, these methods assume that all tasks are homogeneous in scale and type, typically all supervised classification or all regression, such that gradient norm equalization carries a consistent semantic meaning across tasks.

MT-LLE confronts a fundamentally different regime. Its five objectives span discrete classification, continuous geometric reconstruction, probabilistic temporal forecasting, and distributional clustering, operating on vastly different scales where a unit of gradient norm does not carry comparable meaning across task types. Static or gradient-norm-based weighting schemes are therefore insufficient: a weight that appropriately balances geometric and classification losses early in training will actively harm the manifold once a stable topology has been established. Furthermore, while static weights assume the optimal balance between tasks remains constant throughout training, the necessary staged optimization, where global geometry must be established before fine-tuning local clinical boundaries, demands a weighting strategy that evolves over time.

RL offers a principled solution to this dynamic scheduling problem. The application of RL in bioinformatics has largely focused on optimizing computational workflows or feature selection. In the broader machine learning literature, RL frameworks [5] have been proposed to dynamically weight task losses, but these existing approaches are designed for uniform sets of supervised classification tasks where performance metrics are homogeneous. Our contribution is the novel application of an RL agent specifically formulated for biological manifold learning, where the reward is based on the overall improvement in the manifold’s structure rather than a simple predictive metric. Unlike gradient-conflict methods, the RL agent learns a policy over the full composite loss landscape, enabling the adaptive three-phase training schedule, geometric foundation, structural wiring, and supervised fine-tuning that static or gradient-norm approaches cannot replicate.

## Materials and methods

### Preliminaries

We formalize the modeling of heterogeneous diseases as a problem of dimensionality reduction and structure discovery from longitudinal omics data. Let a longitudinal dataset be represented as a tensor *X* ∈ ℝ^*N×T×F*^, where *N* is the number of subjects, *T* denotes the number of longitudinal time points, and *F* represents the high-dimensional feature space of one or more omics modalities (e.g., proteomics, metabolomics). In the single-omics setting, *F* corresponds directly to the feature dimensionality of that modality; in the multi-omics setting, *F* represents the fused feature space as described in subsection “Multi-omics extension: mid-Fusion architecture”. We assume that the biologically relevant signal lies on a lower-dimensional manifold *M* embedded within this high-dimensional space.

Our objective is to learn a parametric mapping function Φ *X* →: Ƶ, where Ƶ ∈ ℝ^*N×T×d*^ is the latent representation and *d* ≪ *F*. Unlike standard dimensionality reduction, we require the latent space Ƶ to satisfy three simultaneous conditions:

1. **Linear Separability:** The manifold must untangle non-linear interactions such that distinct phenotypes (e.g., disease severity stages) become linearly separable.
2. **Temporal Continuity:** The projection *z*_*t*_ must contain sufficient information to predict *z*_*t*+1_, forcing the manifold to respect the arrow of time and enabling forecasting.
3. **Topological Fidelity:** The mapping must preserve the intrinsic local neighborhood structure of the subjects to facilitate meaningful subtype discovery.

### The MT-LLE framework

We introduce **MT-LLE**, a framework for learning continuous disease manifolds from longitudinal omics data. As illustrated in Fig 1, the pipeline operates in two distinct stages to transform high-dimensional patient profiles into a latent space in which disease progression can be read from a patient’s movement across the manifold. In **Stage 1 (Neural Locally Linear Embedding Encoder)**, a deep Neural LLE Encoder maps the data into a low-dimensional manifold, establishing a fundamental geometric structure that respects the intrinsic neighborhood topology. In **Stage 2 (Dynamic Multi-Task Shaping)**, we shape this geometry to satisfy conflicting constraints; an RL agent dynamically balances a composite objective function, ranging from supervised classification to temporal forecasting, to shape the manifold for clinical utility.

**Fig 1.**
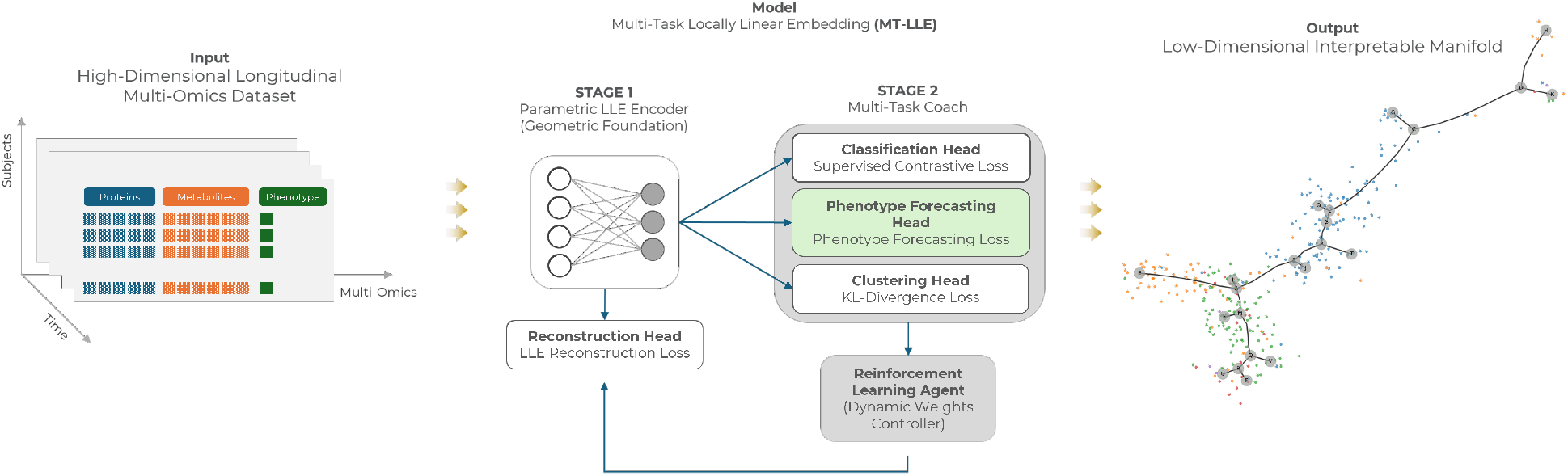
Overview of the multi-task locally linear embedding (MT-LLE) architecture. (A) A parametric locally linear embedding (LLE) encoder maps longitudinal omics data to a latent manifold. (B) The manifold is shaped by five competing objectives: LLE reconstruction, supervised contrastive learning, phenotype forecasting, temporal forecasting, and Kullback–Leibler (KL) divergence. (C) A reinforcement learning (RL) agent observes loss values as states and outputs dynamic weights as actions to balance the optimization.

#### Neural locally linear embedding

Standard LLE is fundamentally transductive, deriving embeddings directly via eigendecomposition without learning a reusable mapping function *f* (*x*). This precludes inductive inference on unseen patients. To enable clinical applicability, we propose neural LLE, a parametric reformulation that trains a deep encoder to approximate the geometric structure. While the supervised objectives (see the subsection “Shaping the manifold: multi-task objectives”) provide a coarse global structure based on class labels, the underlying omics data contain fine-grained information about a sample’s molecular state. The LLE loss acts as a critical regularizer, ensuring that samples with highly similar molecular profiles remain close on the manifold. This preserves local biological similarity that may be missed by discrete class assignments alone. This process unfolds in three steps:

1. **Geometric blueprint (pre-computation):** We first approximate the ideal local geometry by computing reconstruction weights *W*_*ij*_ in the high-dimensional space. For each patient *x*_*i*_, we solve for the weights that minimize || *x*_*i*_ − ∑ *W*_*ij*_*x*_*j*_ ||^2^, treating these as fixed ground-truth targets. To ensure the manifold captures cohort-level biology rather than just patient identity, we impose a constraint during neighbor selection: *N*(*i*) must consist of the *k*-nearest neighbors strictly excluding other timepoints from the same subject. This forces the encoder to define a patient’s state relative to the broader population.
2. **Neural encoder:** We define a deep neural encoder *E*_*ϕ*_ that maps inputs to embeddings: *z*_*i*_ = *E*_*ϕ*_(*x*_*i*_). Unlike standard LLE, this learns a differentiable mapping function *f* (*x*) capable of generalizing to new data.
3. **Reconstruction loss (ℒ**_*LLE*_): We train the encoder to produce embeddings that respect the fixed blueprint *W*. The loss enforces that the embedding *z*_*i*_ can be linearly reconstructed from its cross-sectional neighbors using the fixed weights:

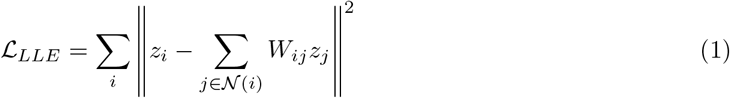

#### Shaping the manifold: multi-task objectives

While the Parametric LLE establishes the fundamental geometric structure, a purely geometric manifold may not be optimal for clinical tasks. We therefore “coach” this manifold with four additional objectives that act as forces within the latent space, bringing the full composite to five jointly optimized objectives: the geometric reconstruction loss of the previous section together with the four introduced here. This multi-task approach ensures that clinical labels (global structure), temporal dynamics (trajectory coherence), and cluster density (subtype definition) synergistically force the latent space to be navigable: clinical labels impose global structure, the two forecasting objectives enforce temporal coherence, and cluster density sharpens subtype definition. The total loss is defined as a composite of these forces, dynamically weighted by the RL agent:

1. **Supervised Contrastive Loss (**ℒ_SupCon_): To enforce global separability, we employ a supervised contrastive loss that acts as an organizing force. Leveraging pre-defined class labels (e.g., disease subtypes, cell types, or treatment outcomes) as anchors, this objective structures the manifold such that high-dimensional samples belonging to the same biological class are pulled together in the embedding space. In contrast, samples from different classes are pushed apart. This creates a global “map” where the main geography is defined by these known sample groups, ensuring the learned manifold is aligned with established biological knowledge. The loss is formally defined as:

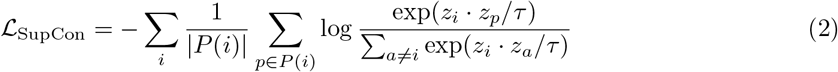

where *z*_*i*_ is the normalized latent embedding of an anchor sample *i. P* (*i*) denotes the set of indices for all positive samples (those sharing the same label as *i*) within the batch, and | *P* (*i*) | is its cardinality. The denominator sums over all distinct samples *a*, and *τ* is a scalar temperature parameter. Consistent with the neighbor selection constraint imposed in the LLE loss, positive pairs are drawn exclusively from cross-sectional subjects sharing the same class label, excluding other timepoints of the anchor subject.
2. **Forecasting Losses ( ℒ**_ZFore_, **ℒ**_YFore_): To transform the static map into a dynamic trajectory, we employ two Transformer-based forecasting losses. This forces the manifold to respect the continuity of disease progression. By requiring that a patient’s current position contains sufficient information to predict their future state, these objectives smooth the transitions between clinical islands and forbid discontinuous jumps, ensuring that patients move along continuous, predictable paths.
  - **Embedding forecasting (ℒ**_ZFore_): This head minimizes the error between the predicted future embedding 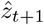 and the actual future embedding *z*_*t*+1_ given the sequence of past states. This strictly enforces smooth, forecastable transitions in the latent space:

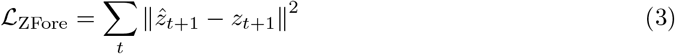
  - **Phenotype forecasting (ℒ**_YFore_): Simultaneously, a second head predicts the future clinical state *y*_*t*+1_. This ensures the trajectory vector aligns with the actual clinical course of the disease:

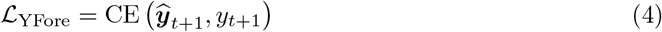
3. **Clustering Regularization (** _KL_): While supervised losses enforce known clinical boundaries, they may overlook subtle molecular sub-structures. To address this, we minimize the KL-divergence between the distribution of embeddings *q*(*z*) and a high-confidence target distribution *p*(*z*). This objective acts as a “densifier” for the latent space, encouraging decisiveness by tightening clusters and clearing the ambiguous space between subtypes. By promoting the formation of dense, well-separated groups, this loss pushes the model towards a data-driven organization, potentially revealing novel biological subgroups that exist within or across the pre-defined clinical classes:

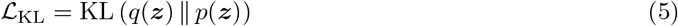

#### Dynamic task weighting via reinforcement learning

Optimizing the composite loss ℒ_total_ is non-trivial due to negative transfer, where competing objectives (e.g., LLE preserving local variance vs. SupCon collapsing clusters) degrade the final representation if weighted statically. To address this, we introduce a Task Weight Controller, an RL agent that acts as a dynamic scheduler. We formulate the weighting problem as a **Markov Decision Process (MDP)** where the agent learns a policy *π*_*ϕ*_ to balance task influence adaptively.

- **MDP Formulation**
  – **State space (***s*_*t*_): The state at step *t* is the vector of L1-normalized loss values from the previous step. This captures the relative difficulty of each task at the current moment:

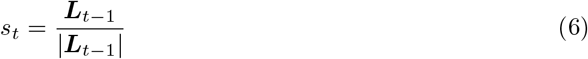

where,

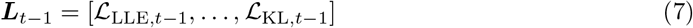
  – **Action space (***a*_*t*_): The action is the weight vector ***λ***_*t*_ applied to the losses for the current update. The weights are sampled from the policy distribution produced by the Controller’s final Softmax layer, ensuring *λ*_*i*_ = 1.
  – **Reward function (***R*_*t*_): To encourage actions that improve model fit, the reward is based on the reduction of the total loss. To stabilize the noisy signal, we use an advantage-based formulation relative to a baseline *B*_*t*−1_, calculated as the Exponential Moving Average (EMA) of past losses:

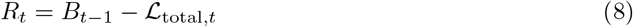

where,

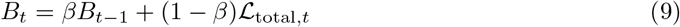

A positive reward is generated only if the agent’s chosen weights result in a loss lower than the recent trend, incentivizing the discovery of efficient optimization paths.
- **Policy optimization**: The agent’s parameters *ϕ* are updated using the REINFORCE policy gradient algorithm. The objective is to maximize the expected future reward by shifting the policy probability mass toward actions that yield positive rewards:

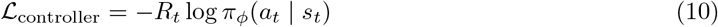

The computational graphs of the main encoder and the RL agent are decoupled. The action weights ***λ***_*t*_ are detached from the gradient tape before computing the main model’s loss. This ensures that the main model does not try to “game” the system by manipulating the weights directly, keeping the two optimization processes stable and independent.

A known limitation of the REINFORCE algorithm is high gradient variance, which can destabilize training. The design choices described above collectively mitigate this: the EMA baseline *B*_*t*_ reduces the variance of the advantage estimate by normalizing rewards against recent performance trends, while the

L1-normalized state representation and decoupled computational graph prevent the optimization from being destabilized by shifting loss magnitudes or adversarial weight manipulation.

#### Multi-omics extension: mid-Fusion architecture

When matched multi-omics data are available, MT-LLE can be extended to a mid-fusion architecture in which multiple independent MT-LLE encoders (one per omics modality) map each modality’s input into a shared *d*-dimensional latent space. The modality-specific manifolds are aligned via a cross-modal neighborhood alignment loss and fused using a bidirectional cross-attention mechanism followed by a learned gate, producing a unified fused representation *z*_fused_. The RL agent’s action and state spaces are extended to include per-modality LLE losses and modality fusion weights, enabling the agent to dynamically balance both task objectives and modality contributions during training. Full architectural details are provided in Section A in S1 Appendix.

### Experimental setup

#### Datasets and preprocessing

We developed and evaluated MT-LLE using omics data from two independent, multi-center longitudinal COPD cohorts.

- **Primary cohort (SPIROMICS):** We utilized data from the Subpopulations and Intermediate Outcome Measures in COPD Study [28], a prospective observational study designed to identify biomarkers for disease progression. The cohort includes current and former smokers with varying degrees of airflow obstruction as well as smoker controls. To ensure a robust longitudinal evaluation, we selected a subset of subjects who completed all four in-person follow-up visits.
- **Independent evaluation cohort (COPDGene):** To assess the generalizability of the learned manifold across independent data, we employed the Genetic Epidemiology of COPD study [29], a multi-center observational study that recruited over 10,000 current and former smokers (*>* 10 pack-years). For this analysis, we utilized proteomic and metabolomic profiles generated at the five-year follow-up visit (2013–2017).

For both cohorts, we curated three distinct experimental datasets to evaluate single-omics and multi-omics capabilities:

1. **Proteomics only:** SPIROMICS (*N* ≈ 162, *F* = 7000, 4 visits) and COPDGene (*N* ≈ 200, 3 visits).
2. **Metabolomics only:** SPIROMICS (*N* ≈ 300, 2 visits) and COPDGene (*N* ≈ 180, 3 visits).
3. **Multi-omics:** A consensus dataset restricted to subjects with complete matched data across both omics modalities (*N* ≈ 162, 2 visits). Multiple fusion strategies were evaluated on this dataset, as described in subsection “Multi-omics extension: mid-Fusion architecture”.

To prevent data leakage, all datasets were partitioned at the subject level (not the visit level). Subjects were split into training, validation, and testing sets, stratified by their baseline GOLD stage to maintain a consistent distribution of disease severity across splits. This ensures that the model is evaluated on its ability to generalize to entirely new patients, rather than just future timepoints of seen patients.

#### Baselines and ablation protocols

To rigorously validate the architectural necessity of the MT-LLE framework, we conducted a full factorial ablation study. Our objective was not merely to demonstrate that the full model yields the highest metrics, but to quantify the marginal contribution of each objective function to the latent geometry, and to examine how competing objectives interact to shape the final representation.

1. **Standard dimensionality reduction:** We benchmarked against UMAP (manifold learning baseline) and a VAE (generative baseline) to assess the value of the multi-task LLE framework over unsupervised dimensionality reduction.
2. **Single-objective baseline (LLE-only):** We trained the encoder using strictly the geometric reconstruction loss (ℒ _*LLE*_). This serves as the geometric baseline, expected to produce a geometrically faithful but clinically unstructured embedding.
3. **Factorial ablation (single vs. multi-task):** We trained intermediate variants with systematically varied objective subsets (e.g., LLE + Classification, LLE + Forecasting, LLE + Clustering, and all pairwise and higher-order combinations) to quantify the marginal contribution of each task and to characterize the gradient interactions between objectives.
4. **Task weighting strategy ablation:** To isolate the contribution of the RL agent specifically, we compared five weighting strategies applied to an identical encoder architecture and loss function set:
  a. **Static uniform**, equal weighting (*λ*_*i*_ = 1*/*5) throughout training.
  b. **GradNorm**, which dynamically adjusts loss weights to equalize gradient norms across tasks.
  c. **PCGrad**, which projects conflicting task gradients to remove interfering components before each update.
  d. **Cosine annealing**, a hand-crafted three-phase schedule that mirrors the structure of the RL agent’s discovered policy, prioritizing geometric objectives early, introducing temporal forecasting and clustering in the second phase, and applying supervised classification last, without any learned adaptation.

Every manifold generated by these variants was subjected to the identical analytical pipeline defined in the “Evaluation methodology” section, ensuring that the evaluation captures not only aggregate geometric scores but also the biological coherence of the resulting manifold.

#### Evaluation methodology

To rigorously validate the MT-LLE framework, we employ a multi-faceted evaluation strategy assessing **intrinsic geometric fidelity, clinical utility**, and **qualitative structural integrity**. To account for training variability arising from random weight initialization, data shuffling, and RL agent initialization, all experiments were repeated across 10 independent runs with distinct random seeds. All reported metrics represent the mean across these runs, with variability expressed as *±* one standard deviation.

#### Intrinsic geometric validation

We quantify the fidelity of the latent manifold using three complementary metrics, designed to assess structural integrity at the local, topological, and global scales.

- **Reconstruction error (***E*_geo_): To verify that the encoder preserves rather than fabricates latent structure, we measure how well the latent space preserves the local neighborhood geometry defined in the high-dimensional omics data. We compute the LLE reconstruction error on the held-out test set:

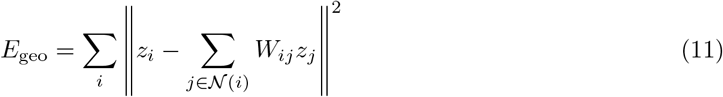

A low error confirms that if two patients are molecularly similar in the original space, they remain neighbors in the learned manifold.
- **Local neighborhood preservation:** We calculate the overlap between the *k*-nearest neighbors in the original space (*N*_orig_) and the latent manifold and the latent manifold (*N*_lat_):

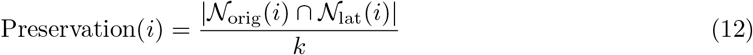

where *k* is the same neighborhood size used in the LLE geometric blueprint construction, treated as a model hyperparameter and selected via grid search (Supplementary Note 1). A high preservation score indicates that the manifold respects the intrinsic local topology of the cohort.
- **Pairwise distance preservation (Global Structure):** To ensure the global geography of the disease landscape is not distorted (e.g., preserving the relative distance between “Healthy” and “Severe” states), we evaluate the correlation between the pairwise distance matrices of the input *X* and embedding Ƶ. We compute the Spearman correlation (*ρ*_dist_):

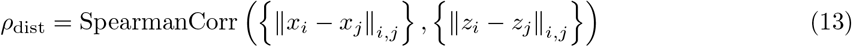

A high correlation confirms that the global hierarchy of patient similarity is maintained.

#### Extrinsic clinical validation

We assess the manifold’s ability to organize patients into clinically coherent states using two complementary approaches.

- **KNN-purity:** Our primary metric for clinical coherence. For every patient in the latent space, we compute the fraction of their 10 nearest neighbors that share the same ground-truth GOLD stage, where in this evaluation *k* is distinct from the *k* used in the LLE geometric blueprint construction. We selected *k* = 10 as a representative value and verified via sensitivity analysis that KNN-Purity remains stable as *k* increases (Supplementary Note 1), confirming that the reported values are not sensitive to this choice. Higher purity indicates successful disentanglement of distinct disease states in the embedding space.
- **Downstream task performance:** To evaluate the quality of the learned representation independently of the training objective, we extract frozen embeddings from the trained MT-LLE encoder and train separate downstream models on these embeddings, with no gradient flow back to the encoder. This design ensures that downstream performance reflects the intrinsic quality of the learned manifold rather than the capacity of jointly optimized task heads. We assess three tasks:
  1. **Classification:** A linear classifier is trained on the frozen training embeddings and evaluated on the held-out test embeddings. Performance is reported as F1-Macro score (the unweighted mean of per-class F1 scores, accounting for class imbalance across GOLD stages), verifying that distinct disease states are linearly separable in the learned manifold.
  2. **Clustering:** Silhouette Score computed directly on the frozen test embeddings, measuring the compactness and separation of disease subtypes without requiring any additional training.
  3. **Forecasting:** A Transformer-based sequence model is trained on the sequences of frozen training embeddings and evaluated on held-out test sequences. Performance is reported as the F1-Macro score for predicting the future GOLD stage at visit *t* + 1 given the embedding trajectory up to visit *t*. This verifies that the manifold encodes predictive temporal progression signals rather than static molecular snapshots. Note that this downstream forecaster is trained independently of the Transformer-based forecasting head used during MT-LLE training, ensuring a clean separation between representation learning and evaluation.

#### Qualitative validation

Beyond scalar metrics, we employ visual diagnostic tools to detect topological artifacts that aggregate statistics might mask.

- **Shepard diagrams (distortion check):** Scatter plots of pairwise input distances versus latent distances visualize global stress, revealing topological tearing (nearby inputs mapped far apart) or collapsing (distant inputs crushed together) artifacts.
- **Latent interpolation (continuity check):** We linearly interpolate between the centroids of the healthy (*z*_healthy_) and severe (*z*_severe_) subpopulations,

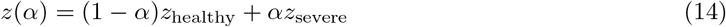

where *α* ∈ [0, 1], and decode the resulting path back to the omics feature space. A geometrically sound manifold yields gradual, coherent shifts in feature expression; a degenerate manifold produces abrupt discontinuities or unrealistic intermediate states.
- **Topological stability (robustness check):** We train the model on 10 independent random subsets, each comprising 50% of training subjects, align the resulting manifolds via Procrustes transformation, and quantify structural consistency using the average centroid shift across clinical subgroups.

### Manifold interpretation and disease mechanism discovery

Beyond quantitative benchmarking, we illustrate the analytical versatility of the learned manifold through three interpretive case studies spanning static phenotyping, trajectory inference, and longitudinal kinematic analysis, demonstrating the range of biological questions that the manifold enables researchers and clinicians to address.

#### Case study I: static phenotyping and functional annotation

To validate that dense manifold regions represent distinct molecular phenotypes rather than arbitrary geometric clusters, we employ a three-step protocol:

1. **Manifold clustering:** We apply K-Means clustering to the latent representation Ƶ, initializing the number of clusters at *k* = 5, motivated by the five GOLD severity stages used to characterize disease severity in both cohorts. The biological coherence of the resulting partition was assessed by examining the differential abundance profiles of each cluster; regions exhibiting highly similar molecular signatures were subsequently merged. This domain-informed initialization followed by molecular validation ensures that the final partition reflects genuine biological heterogeneity rather than arbitrary geometric boundaries.
2. **Differential abundance analysis:** For each identified region, we computed a region-level standardized abundance score (*Z*-score) for every molecular feature in the original high-dimensional feature space. To account for repeated longitudinal measurements and prevent participants with more visits from disproportionately influencing the result, visit-level abundances were first averaged within each participant and region, so that each participant contributed a single value per region. The *Z*-score for each feature was then calculated as the difference between the region-level mean (the mean of participant-level averages within the region) and the cohort-wide mean (the mean of participant-level averages across the full cohort), divided by the cohort-wide standard deviation. Positive *Z*-scores indicated higher mean abundance in the focal region relative to the overall cohort. Regions with fewer than ten participants, or with a maximum absolute *Z*-score exceeding a data-driven anomaly threshold (the mean plus two standard deviations of the maximum *Z*-scores across all regions), were flagged and excluded from biological interpretation, as small participant counts can inflate *Z*-scores independently of genuine biological signal. Among the remaining regions, the highest-ranking features by *Z*-score magnitude were retained for functional annotation. This analysis is descriptive rather than inferential: identified features are interpreted as region-associated molecular markers reflecting the manifold’s organization of the cohort, rather than as statistically validated or causal biological findings
3. **Functional annotations:** Given the targeted size of the identified marker sets, we performed manual thematic annotation rather than automated enrichment analysis. For proteomic markers, proteins were annotated by physiological role using the UniProt Knowledgebase, categorizing top drivers into functional themes (e.g., “tissue remodeling” vs. “innate immunity”). For metabolomic markers, metabolites were annotated using the Human Metabolome Database (HMDB) [30], mapping each metabolite to its associated biological pathway and physiological class.

#### Case study II: trajectory inference and bifurcation analysis

To characterize the continuous progression axes encoded in the manifold and identify points of molecular divergence, we employ Elastic Principal Graphs (ElPiGraph) [31] as a skeletonization tool. Critically, ElPiGraph is not used to learn relationships between patients; rather, it traces the progression axes that MT-LLE has already constructed by fitting a branching principal graph (a set of nodes and edges minimizing the Mean Squared Error of Projection (MSEP)) into the dense point cloud of the embedding space. This serves as a non-linear extension of Principal Component Analysis: instead of fitting a linear vector, it fits a branching curve that abstracts the manifold into a progression skeleton.

1. **Trajectory skeletonization:** We use ElPiGraph to fit a principal graph to the latent manifold, identifying the main progression trunk and significant branches. The root node was defined as the graph node closest to the centroid of subjects with GOLD stage 0 and no history of exacerbation, representing the most homeostatic molecular state in the cohort.
2. **Bifurcation analysis:** At branching points, defined as graph nodes with degree greater than 2, we identified the participant-level observations assigned to each diverging path and performed directional pairwise comparisons between branches, following the same participant-averaging procedure, standardized abundance score (*Z*-score) calculation, and minimum-participant and anomaly-threshold exclusion criteria described in “Case study I.” The principal difference is that each *Z*-score here reflects a direct comparison between two diverging branches, rather than a region against the full cohort: for each pairwise comparison, the *Z*-score for a given protein was calculated as the difference between its mean abundance in the named trajectory axis and the comparison branch, divided by the pooled standard deviation across both branches, with positive values indicating higher abundance in the named axis. Among comparisons passing the participant-count and anomaly-threshold criteria, the highest-ranking proteins by *Z*-score magnitude were assigned to functional themes using the annotation protocol described in “Case study I.” As in Case study I, these proteins are interpreted as candidate molecular descriptors of branch separation rather than as statistically validated or causal biological findings.

#### Case study III: kinematic phenotyping of longitudinal progression

Although the principal graph identifies potential progression pathways, it does not quantify the rate of individual movement along these axes. We therefore introduce a kinematic characterization of disease progression: rather than describing a patient by their static position on the manifold, we describe them by how they move through it, their speed and acceleration over calendar time, and cluster subjects on these dynamics. This shifts the unit of analysis from the molecular state to the molecular trajectory, surfacing phenotypes that static profiling cannot express.

- **Pseudotime kinematics (***v* **and** *a*): For all subjects with at least two longitudinal visits, we compute pseudotime speed and acceleration relative to the actual calendar time elapsed between visits. The pseudotime speed for subject *i* between visits *t* and *t* + 1 is defined as:

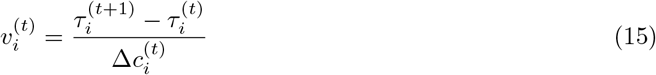

where 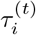 is the pseudotime coordinate of subject *i* at visit *t* and 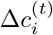 is the elapsed calendar time in years. Kinematic Acceleration is defined as:

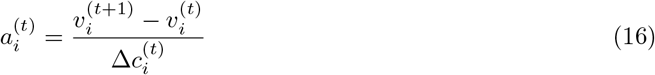

**Kinematic acceleration (***a*) is defined as the directional vector representing the rate of change of that speed over time.
- **Trajectory clustering:** The kinematic profiles (*v, a*) for each subject are aggregated into a feature matrix and partitioned using K-Means clustering. The number of clusters was set to *k* = 3, selected using the elbow criterion applied to the sum of squares within the cluster of *k* ∈ *{*2, …, 6*}*.
- **Molecular characterization:** To identify the biological drivers of each kinetic phenotype, we performed one-versus-rest differential abundance analysis on each trajectory cluster using the same participant-averaging procedure, standardized abundance score (*Z*-score) calculation, and minimum-participant and anomaly-threshold exclusion criteria described in “Case study I.” Since cluster membership was assigned once per subject rather than per visit, no participant could be assigned to both a focal cluster and the remainder, so no additional exclusion was required beyond this averaging. Among clusters passing the minimum-participant criterion, the highest-ranking proteins by *Z*-score magnitude were mapped to functional themes using the annotation protocol described in “Case study I.”

## Results

### MT-LLE optimizes the trade-off between geometric fidelity and clinical utility

We empirically validated the MT-LLE framework against baselines and ablated variants along three dimensions: intrinsic geometric quality (Table 1), local phenotype coherence, and downstream transferability (Table 2).

**Table 1.**
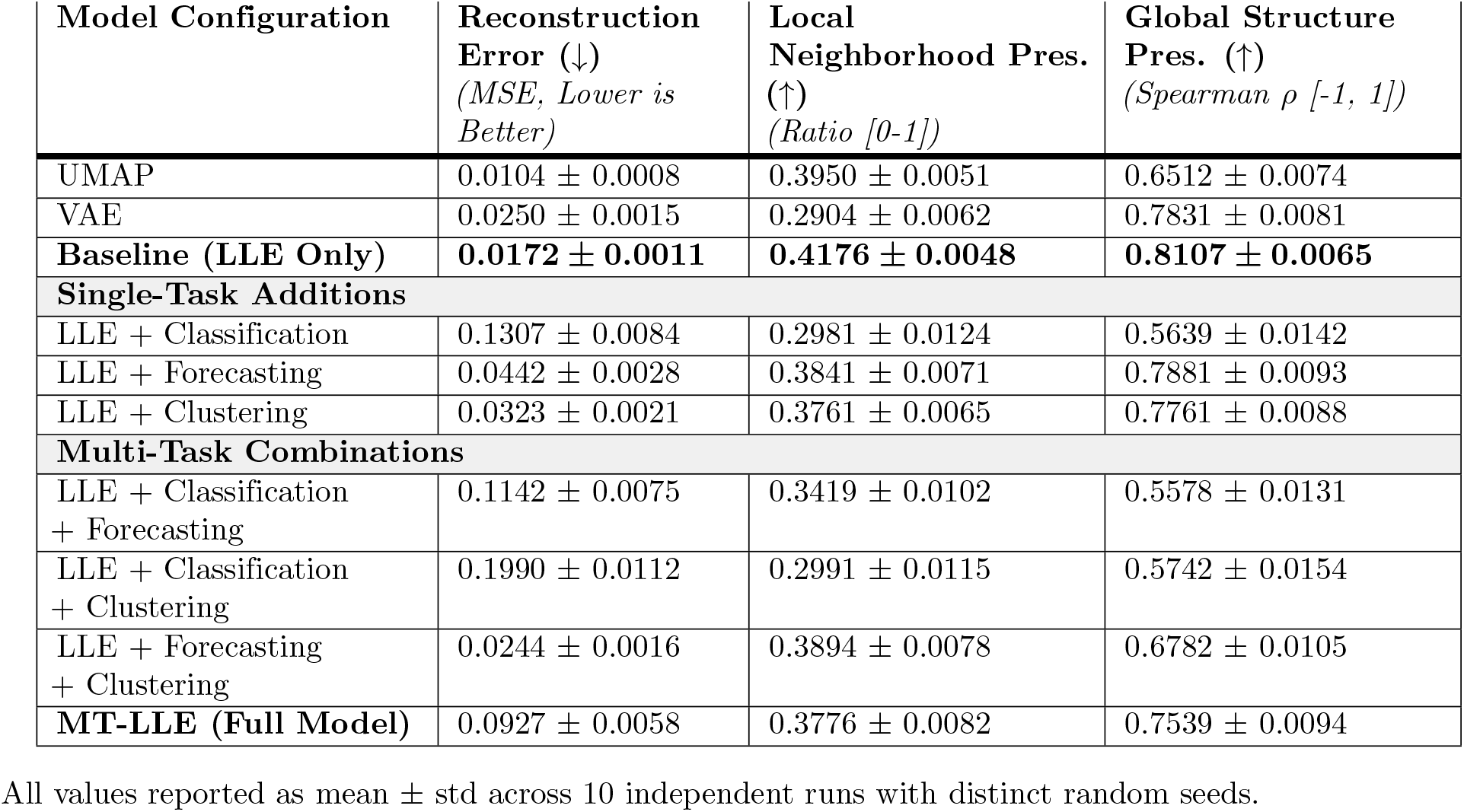
Intrinsic geometric quality. Performance of MT-LLE and baseline models on the SPIROMICS proteomics cohort.

| Model Configuration | Reconstruction Error (↓)<br>(MSE, Lower is Better) | Local Neighborhood Pres. (↑)<br>(Ratio [0-1]) | Global Structure Pres. (↑)<br>(Spearman $\rho$ [-1, 1]) |
| --- | --- | --- | --- |
| UMAP | 0.0104 ± 0.0008 | 0.3950 ± 0.0051 | 0.6512 ± 0.0074 |
| VAE | 0.0250 ± 0.0015 | 0.2904 ± 0.0062 | 0.7831 ± 0.0081 |
| **Baseline (LLE Only)** | **0.0172 ± 0.0011** | **0.4176 ± 0.0048** | **0.8107 ± 0.0065** |
| **Single-Task Additions** |  |  |  |
| LLE + Classification | 0.1307 ± 0.0084 | 0.2981 ± 0.0124 | 0.5639 ± 0.0142 |
| LLE + Forecasting | 0.0442 ± 0.0028 | 0.3841 ± 0.0071 | 0.7881 ± 0.0093 |
| LLE + Clustering | 0.0323 ± 0.0021 | 0.3761 ± 0.0065 | 0.7761 ± 0.0088 |
| **Multi-Task Combinations** |  |  |  |
| LLE + Classification + Forecasting | 0.1142 ± 0.0075 | 0.3419 ± 0.0102 | 0.5578 ± 0.0131 |
| LLE + Classification + Clustering | 0.1990 ± 0.0112 | 0.2991 ± 0.0115 | 0.5742 ± 0.0154 |
| LLE + Forecasting + Clustering | 0.0244 ± 0.0016 | 0.3894 ± 0.0078 | 0.6782 ± 0.0105 |
| **MT-LLE (Full Model)** | 0.0927 ± 0.0058 | 0.3776 ± 0.0082 | 0.7539 ± 0.0094 |

All values reported as mean ± std across 10 independent runs with distinct random seeds.

**Table 2.**
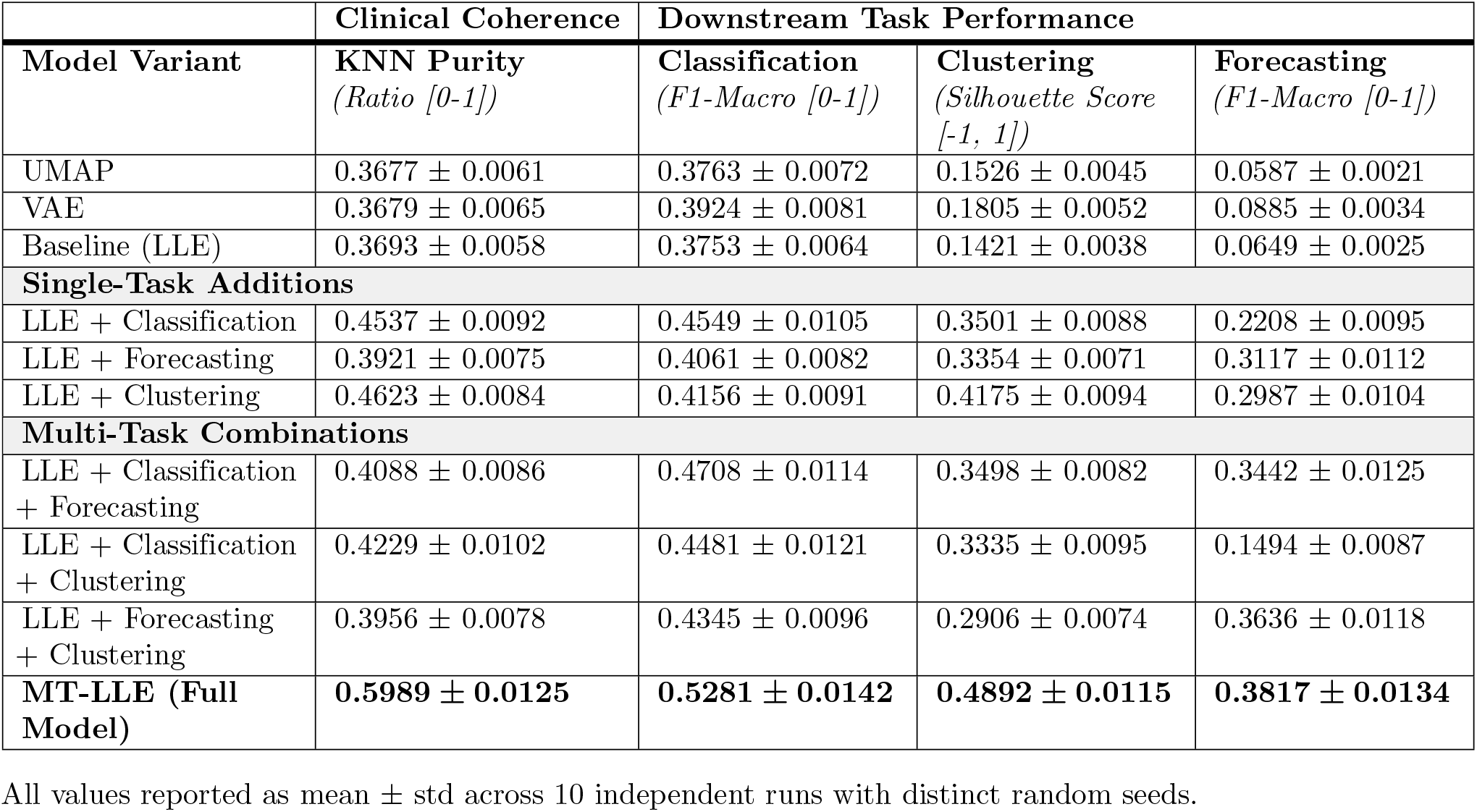
Manifold coherence and downstream task performance. Performance of MT-LLE and baseline models on the SPIROMICS proteomics cohort.

|  | Clinical Coherence | Downstream Task Performance |  |  |
| --- | --- | --- | --- | --- |
| Model Variant | KNN Purity<br>(Ratio [0-1]) | Classification<br>(F1-Macro [0-1]) | Clustering<br>(Silhouette Score [-1, 1]) | Forecasting<br>(F1-Macro [0-1]) |
| UMAP | 0.3677 ± 0.0061 | 0.3763 ± 0.0072 | 0.1526 ± 0.0045 | 0.0587 ± 0.0021 |
| VAE | 0.3679 ± 0.0065 | 0.3924 ± 0.0081 | 0.1805 ± 0.0052 | 0.0885 ± 0.0034 |
| Baseline (LLE) | 0.3693 ± 0.0058 | 0.3753 ± 0.0064 | 0.1421 ± 0.0038 | 0.0649 ± 0.0025 |
| **Single-Task Additions** |  |  |  |  |
| LLE + Classification | 0.4537 ± 0.0092 | 0.4549 ± 0.0105 | 0.3501 ± 0.0088 | 0.2208 ± 0.0095 |
| LLE + Forecasting | 0.3921 ± 0.0075 | 0.4061 ± 0.0082 | 0.3354 ± 0.0071 | 0.3117 ± 0.0112 |
| LLE + Clustering | 0.4623 ± 0.0084 | 0.4156 ± 0.0091 | 0.4175 ± 0.0094 | 0.2987 ± 0.0104 |
| **Multi-Task Combinations** |  |  |  |  |
| LLE + Classification + Forecasting | 0.4088 ± 0.0086 | 0.4708 ± 0.0114 | 0.3498 ± 0.0082 | 0.3442 ± 0.0125 |
| LLE + Classification + Clustering | 0.4229 ± 0.0102 | 0.4481 ± 0.0121 | 0.3335 ± 0.0095 | 0.1494 ± 0.0087 |
| LLE + Forecasting + Clustering | 0.3956 ± 0.0078 | 0.4345 ± 0.0096 | 0.2906 ± 0.0074 | 0.3636 ± 0.0118 |
| **MT-LLE (Full Model)** | **0.5989 ± 0.0125** | **0.5281 ± 0.0142** | **0.4892 ± 0.0115** | **0.3817 ± 0.0134** |

All values reported as mean ± std across 10 independent runs with distinct random seeds.

It is critical to note that our objective is not simply to surpass the unsupervised baselines on intrinsic geometric measures. As our downstream evaluations demonstrate, a manifold generated using purely unsupervised approaches lacks clinical actionability, while naive supervised forcing actively destroys the underlying biological geometry. Ultimately, geometric fidelity is insufficient if the resulting space does not align with clinical reality; **MT-LLE** is designed to explicitly optimize the trade-off between these two competing requirements.

To provide a coherent clinical and geometric narrative, the detailed quantitative, qualitative, and case study analyses in the following subsections focus primarily on the SPIROMICS proteomics dataset, selected for its high-resolution temporal sequencing (4 visits). The MT-LLE framework was rigorously benchmarked across multiple independent cohorts and molecular layers, including COPDGene and paired metabolomics; these generalizability and multi-omics fusion results are detailed in Section.

#### The failure of standard dimensionality reduction

Unsupervised baselines cannot resolve heterogeneous clinical states. Standard algorithms optimize for generic variance or local neighborhoods without regard for biological semantics, generating manifolds blind to clinical labels and temporal ordering that cannot disentangle complex, overlapping disease states. As shown in Table 1, the baseline LLE model achieves the lowest reconstruction error (0.0172 *±* 0.001), but this geometrically faithful embedding is clinically uninformative. Because it lacks any mechanism to encode temporal ordering or clinical labels, its performance across all downstream tasks is severely degraded (Classification F1-Macro: 0.3753 *±* 0.006; Forecasting F1-Macro: 0.0649 *±* 0.003; Silhouette Score: 0.1421 *±* 0.004; Table 2).

#### The cost of naive supervision and the gradient tug-of-war

Attempting to remedy this by applying supervised labels directly destroys the underlying biological geometry. Adding a supervised classification objective (LLE + Classification) causes a substantial increase in reconstruction error (0.1307 *±* 0.008), as the classification loss forces the continuous molecular space into discrete, rigid boundaries corresponding to noisy GOLD stage labels, fragmenting the underlying omics geometry.

This creates a fundamental optimization conflict when classification and clustering are combined (LLE + Classification + Clustering). The clustering objective attempts to group molecularly similar patients (for instance, a GOLD 2 and a GOLD 4 subject with comparable omics profiles) while the classification objective simultaneously separates them to satisfy clinical stage boundaries. This gradient conflict prevents either task from reaching its optimal configuration. Consequently, cluster separation degrades (Silhouette Score: 0.3335 *±* 0.010, below the clustering-only variant), classification performance decreases (F1-Macro: 0.4481 *±* 0.012, below classification alone), downstream forecasting degrades severely (F1-Macro: 0.1494 *±* 0.009), and reconstruction error rises sharply (0.1990 *±* 0.011).

#### Unsupervised clustering as the engine for temporal forecasting

Accurate trajectory forecasting requires grouping patients by their underlying molecular progression mechanism rather than by lagging clinical symptoms. When anchored purely by clinical labels (LLE + Classification), temporal forecasting remains weak (F1: 0.2208 *±* 0.010), because patients sharing a GOLD stage label may be on fundamentally different biological trajectories. Unsupervised clustering, by contrast, organizes the latent space according to intrinsic data density rather than predefined clinical categories. This density-based organization naturally produces molecularly coherent groupings that align with true disease progression pathways, substantially improving forecasting even without explicit temporal supervision (LLE + Clustering: Forecasting F1-Macro 0.2987 *±* 0.010, Silhouette Score 0.4175 *±* 0.009). As temporal forecasting is added alongside clustering, the two objectives synergize to reach a Forecasting F1 of 0.3636 *±* 0.012 (LLE + Forecasting + Clustering).

#### The synergy of forecasting and clustering

This synergy produces an interpretable trade-off in geometric metrics. When temporal forecasting and clustering are combined (LLE + Forecasting + Clustering), the forecasting objective pulls embeddings along a temporal axis, stretching spherical cluster formations into elongated trajectory structures. The Silhouette Score, which penalizes elongated and overlapping cluster shapes, consequently decreases to 0.2906 *±* 0.007. However, this geometric deformation is mechanistically appropriate: the goal is not to produce isolated, compact clusters but to construct a continuous, temporally ordered latent space in which molecular progression is forecastable.

#### Dynamic RL phasing solves the trade-off

To harmonize these conflicting objectives, the RL agent discovers a staged optimization policy rather than applying all objectives simultaneously. As shown in Fig 2, the agent first prioritizes geometric reconstruction to establish a stable topological foundation, then introduces clustering and temporal forecasting to organize the space into molecularly coherent hubs and sequence their transitions, and finally applies supervised classification only after this structural foundation is established. This phased approach prevents the gradient conflicts observed in the static ablations above, allowing supervised losses to refine existing biological boundaries rather than tear the underlying geometry.

**Fig 2.**
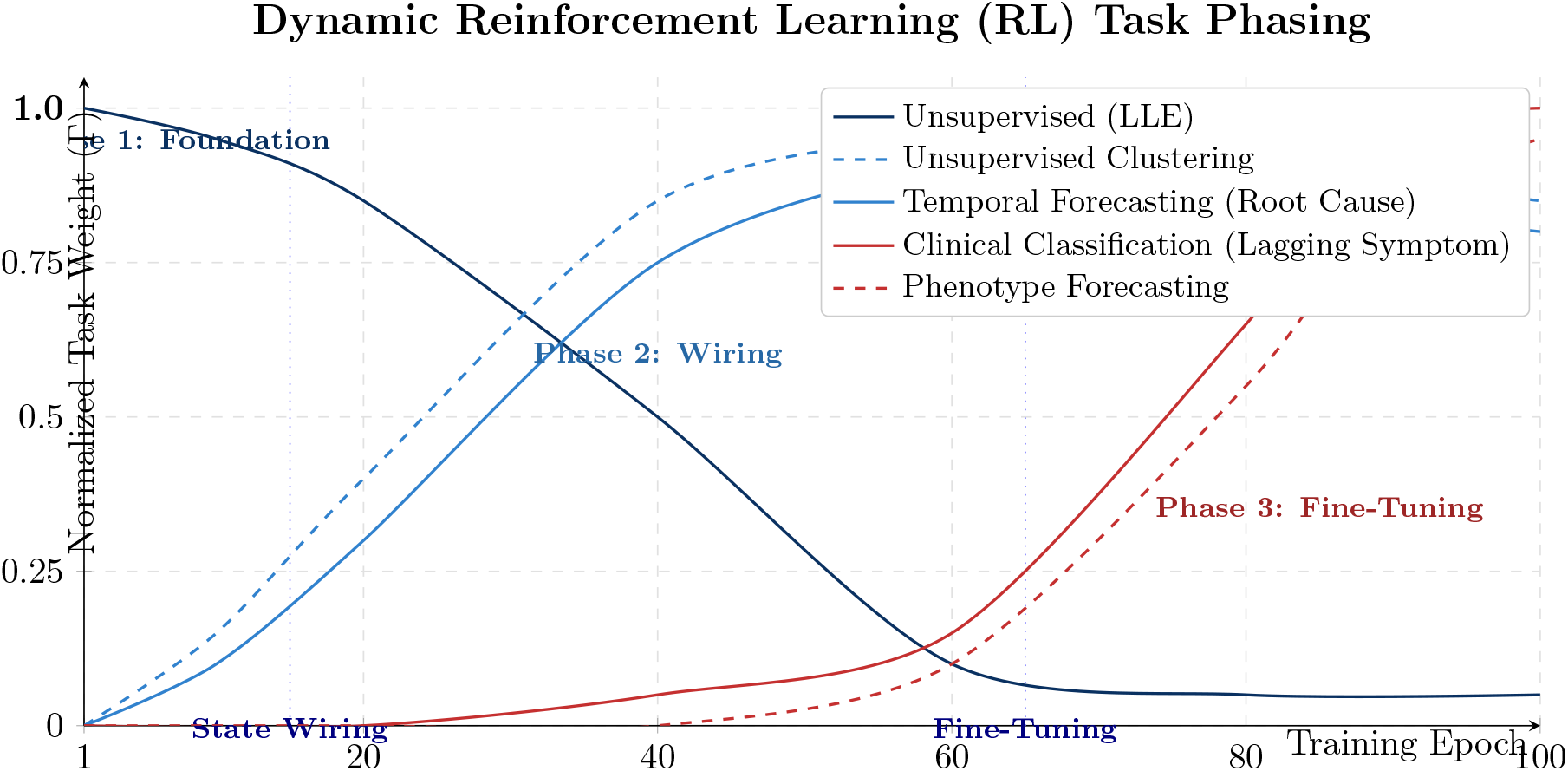
Dynamic Reinforcement Learning (RL) Task Phasing. The RL agent optimizes the trade-off between geometric fidelity and clinical utility by dynamically phasing task weights (Γ). **Phase 1 (Foundation):** The model prioritizes unsupervised LLE to build the baseline molecular topology. **Phase 2 (Wiring):** Clustering and Temporal Forecasting rise to organize the space into dense biological states and sequence their transitions. **Phase 3 (Fine-Tuning):** Rigid Clinical Classification and Phenotype Forecasting are applied only after the manifold is stable, fine-tuning clinical boundaries without tearing the established biological hubs.

Through this dynamic balancing, the full MT-LLE model recovers global structure preservation to 0.7539 *±* 0.009 while simultaneously achieving peak KNN purity (0.5989 *±* 0.013), downstream classification (0.5281 *±* 0.014 F1-Macro, a 35-40% relative improvement over the UMAP and VAE baselines in Table 2), clustering (0.4892 *±* 0.012), and forecasting (0.3817 *±* 0.013), demonstrating that the trade-off between geometric fidelity and clinical utility can be systematically optimized rather than left to chance.

#### Ablation of task weighting strategies

To isolate whether performance gains derive from dynamic task weighting in general or from the RL formulation specifically, we benchmarked all five weighting strategies on the SPIROMICS proteomics dataset across 10 independent seeds (Fig 3; exact values for all metrics are reported in Tables A and B in S1 Appendix).

**Fig 3.**
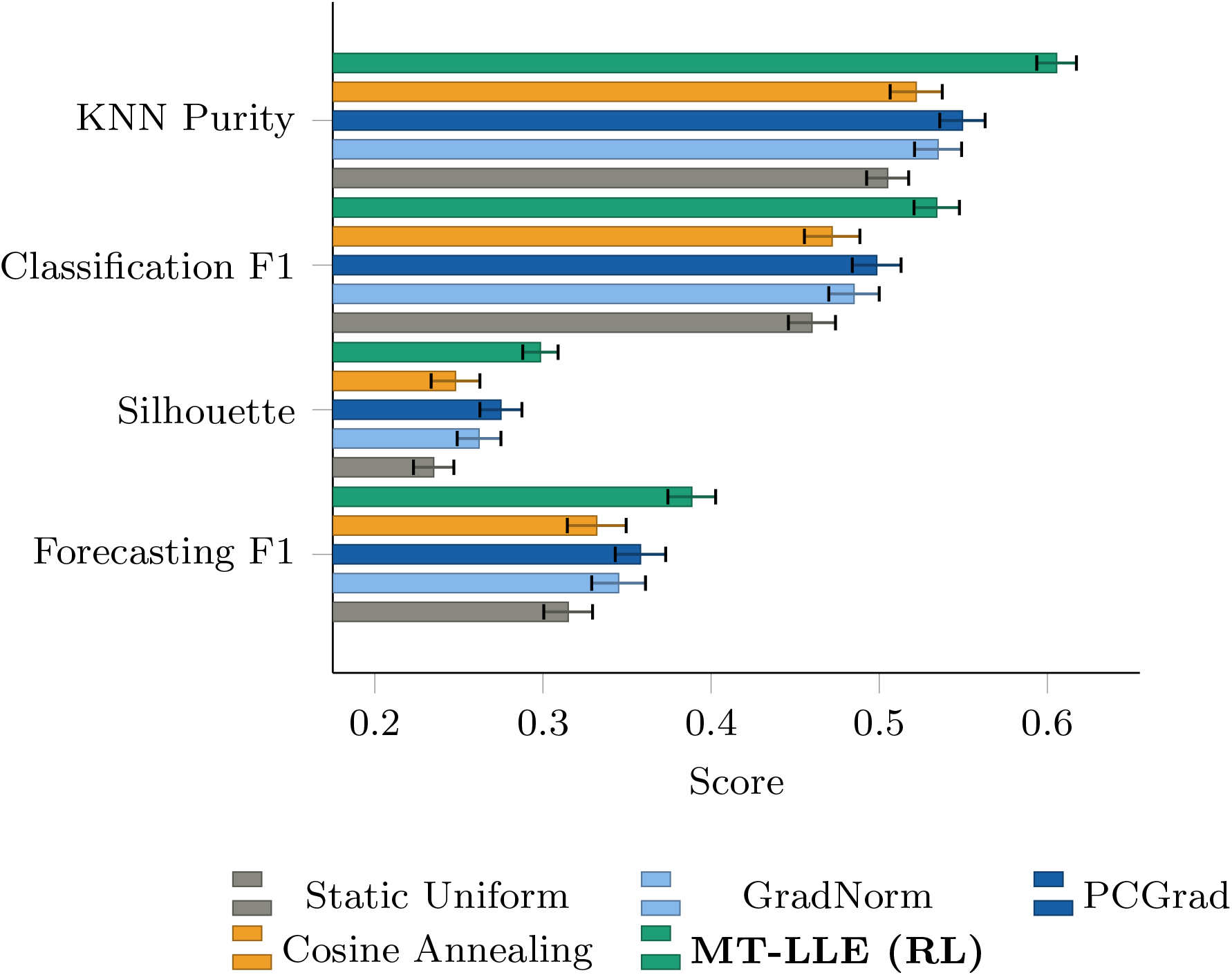
Downstream task performance across task-weighting strategies. MT-LLE with reinforcement learning (RL) achieves the highest value on all four metrics. Its improvement over cosine annealing indicates that the adaptive RL policy contributes beyond phase ordering alone. Values are reported as mean *±* standard deviation across 10 independent runs with distinct random seeds.

Static Uniform weighting already improves substantially over the unsupervised baselines in Table 2 (KNN Purity: 0.505 *±* 0.013 vs. 0.369), confirming that the multi-task objective set contributes independently of how task weights are assigned. Among the adaptive comparators, PCGrad consistently outperforms GradNorm across all downstream metrics despite lower computational overhead (0.082 vs. 0.101 seconds per epoch), suggesting that resolving gradient conflicts directly is more effective than equalizing gradient norms in this heterogeneous objective setting.

The critical comparison is between Cosine Annealing and the MT-LLE RL agent. Because the Cosine Annealing schedule replicates the three-phase structure of the RL agent’s discovered policy without any learned adaptation, it directly tests whether the phase structure alone, rather than the learned policy, accounts for the gains. MT-LLE substantially outperforms Cosine Annealing on all clinical metrics, with absolute improvements of 0.084 in KNN Purity, 0.062 in Classification F1, 0.057 in Forecasting F1, and 0.051 in Silhouette Score — relative improvements of 16%, 13%, 17%, and 20% respectively, all supported by non-overlapping confidence intervals. The RL agent’s adaptive, data-driven weight tuning within and across phases therefore provides measurable value that a hand-crafted schedule derived from domain knowledge cannot recover.

This performance advantage carries negligible computational cost: the RL agent adds only 0.003 seconds per epoch over Cosine Annealing, a 6% overhead. By contrast, GradNorm incurs 89% greater cost than the RL agent while underperforming it on every metric. As shown in Fig 4, among the five weighting strategies, MT-LLE simultaneously achieves the lowest reconstruction error and the highest local neighborhood and global structure preservation, confirming that improved clinical utility does not come at the expense of geometric fidelity. All differences relative to the next-best comparator (PCGrad) are supported by non-overlapping standard deviations.

**Fig 4.**
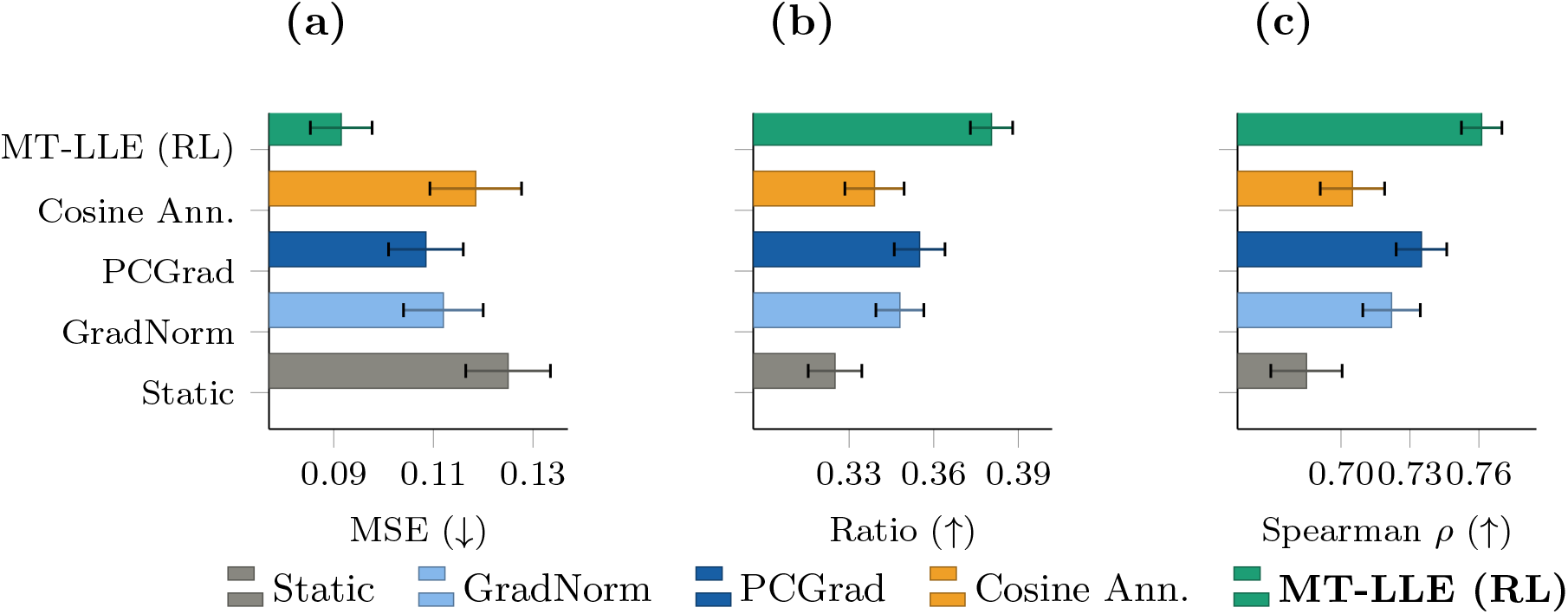
Intrinsic geometric quality across task-weighting strategies. (A) Reconstruction error, measured by mean squared error (MSE), for which lower values indicate better performance. (B) Local neighborhood preservation. (C) Global structure preservation. MT-LLE with reinforcement learning (RL) simultaneously achieves the lowest reconstruction error and the highest local and global structure preservation, indicating that the clinical utility gains shown in Fig 3 do not come at the expense of geometric fidelity. Values are reported as mean *±* standard deviation across 10 independent runs with distinct random seeds. Exact values are provided in Tables A and B in S1 Appendix.

### Qualitative geometric verification

Beyond aggregate metrics, we employed visual diagnostics to verify the structural integrity and reproducibility of the learned manifold.

#### Shepard diagrams and controlled tearing

Global structure analysis via Shepard diagrams (Fig 5) provides a visual diagnostic of how multi-task gradients reshape the biological manifold. The **Baseline (LLE Only)** variant serves as the geometric anchor, maintaining a tight diagonal formation and high correlation (*ρ* = 0.7071), confirming local distance preservation in the absence of clinical constraints.

**Fig 5.**
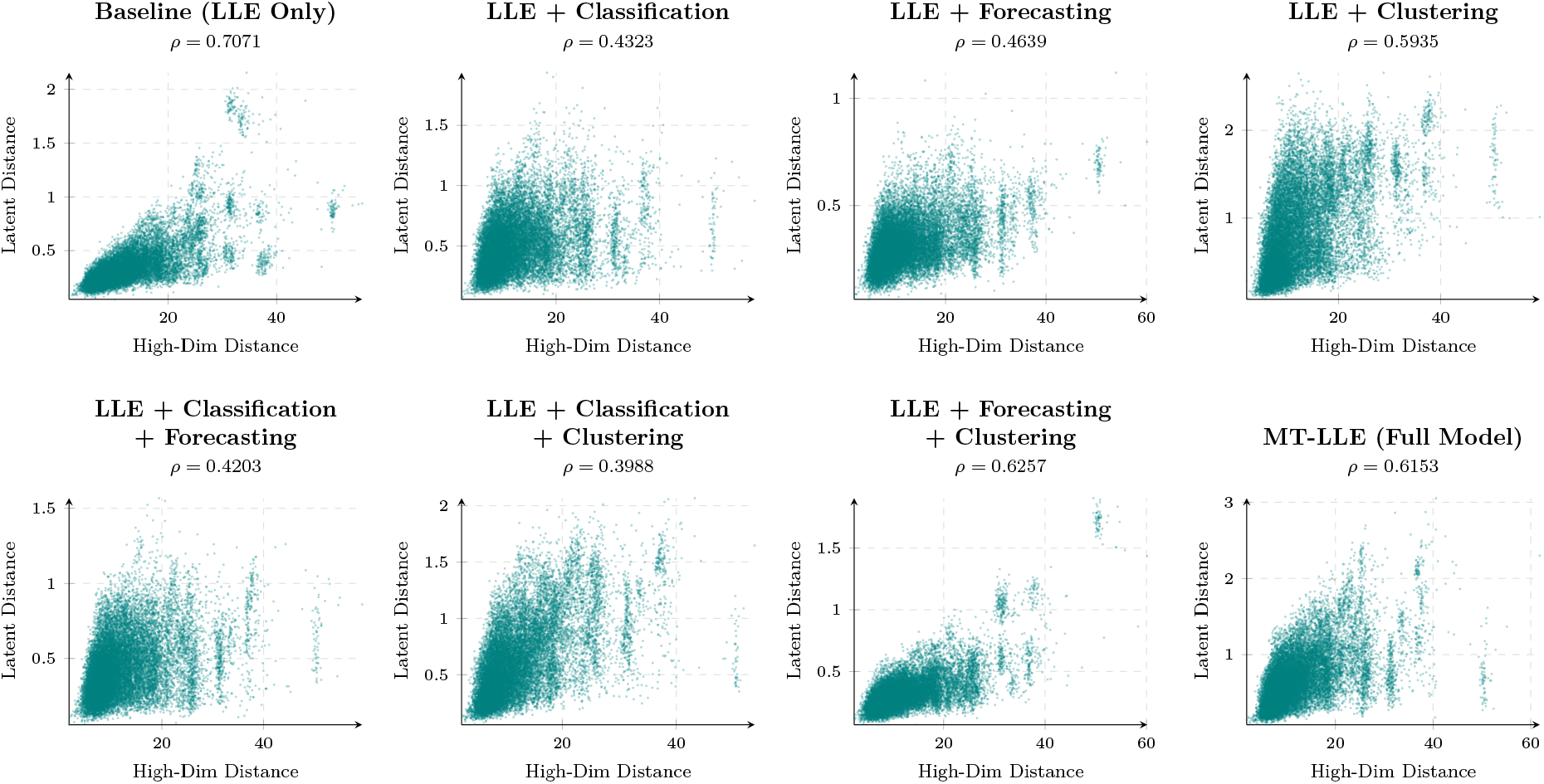
Shepard diagrams of global manifold structure. Shepard diagrams show the preservation of pairwise distances across model variants. The *x*-axis represents pairwise distances in the original high-dimensional proteomic space, and the *y*-axis represents the corresponding distances in the latent space. The baseline model in the upper-left panel primarily preserves local geometry, whereas MT-LLE with reinforcement learning in the lower-right panel exhibits *controlled tearing* : pillar-like vertical structures that reflect the separation of molecularly heterogeneous samples while retaining overall global structure, with a Spearman correlation of *ρ* = 0.6153.

In contrast, the **LLE + Classification** variant reveals the cost of discrete supervision: the correlation decreases to *ρ* = 0.4323, manifesting as broad vertical smearing in which the model forcibly fragments the local geometry to assign patients into rigid, predetermined GOLD stage boundaries. This demonstrates that naive supervision destroys the continuous biological progression encoded in the omics data.

Crucially, the **MT-LLE (Full Model)** recovers this structural integrity, restoring the correlation to *ρ* = 0.6153. The diagram exhibits controlled tearing: distinct, dense, pillar-like vertical columns branching off a stable main diagonal. This is the visual signature of the RL controller successfully resolving task conflict. The agent identifies groups of patients who appear molecularly similar in the raw data and separates them into distinct manifold regions. As described in the subsection “Case study I: Static phenotyping and functional annotation”, this mechanism allows the manifold to resolve molecular heterogeneity within clinically asymptomatic populations, separating molecularly active from stable subjects within the GOLD 0 group.

#### Latent interpolation and biological metastability

The MT-LLE interpolation path (Fig 6) reveals that while molecular transitions are mathematically continuous, they are biologically non-linear. The presence of stepped phase transitions, particularly between interpolation steps 16 and 20, provides visual evidence of biological metastability: patients appear to persist in stable molecular regimes until a cumulative proteomic tipping point triggers a shift toward more severe disease phenotypes.

**Fig 6.**
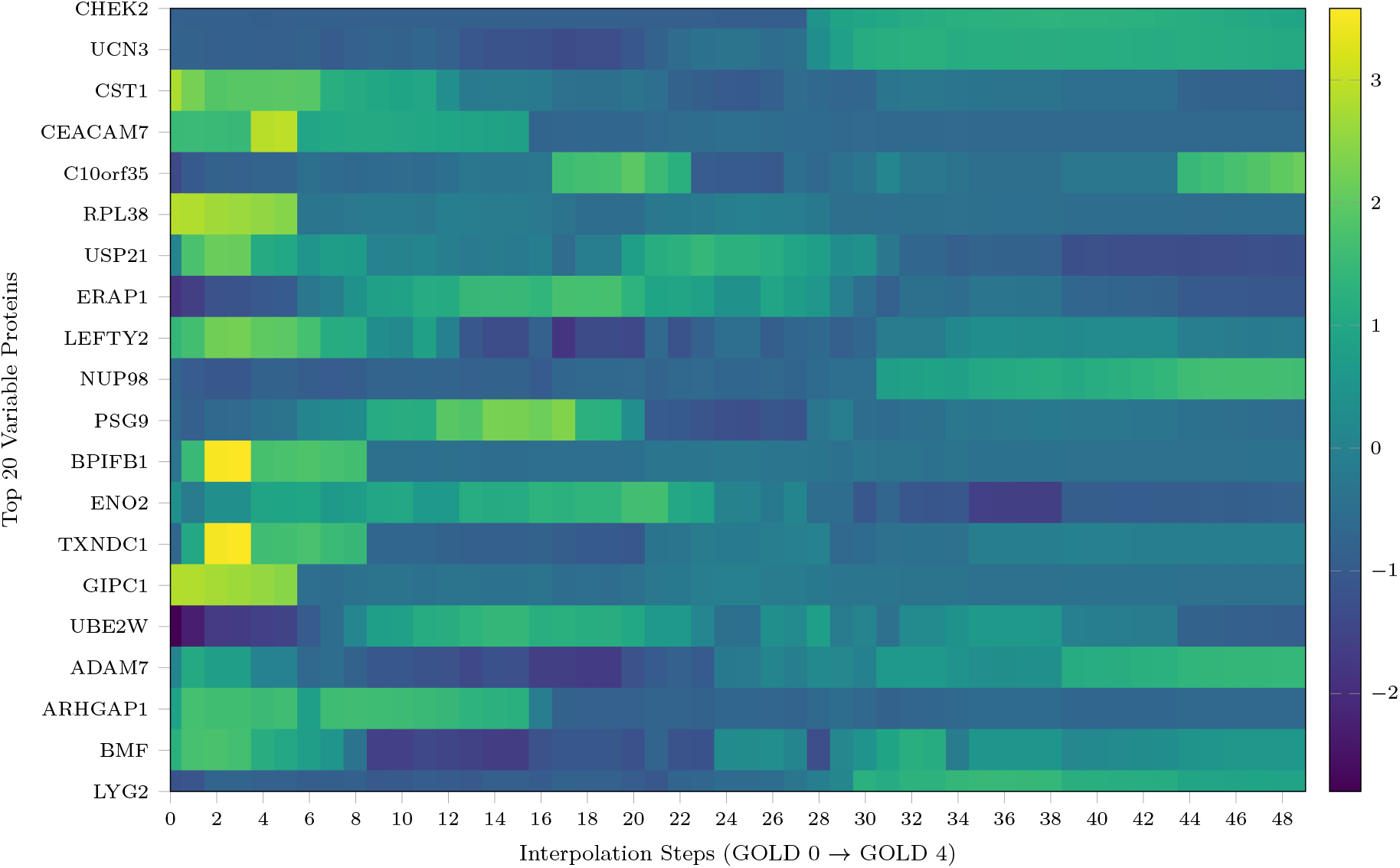
Latent interpolation from GOLD 0 to GOLD 4. Expression heatmap for the 20 most variable proteins across the interpolation trajectory. Stepped transitions highlight molecular tipping points, while the expression gradients of BPIFB1 and CHEK2 align with the progressive loss of airway defense and terminal cellular senescence, respectively.

The underlying protein signatures corroborate established COPD progression narratives:

- **Barrier collapse:** BPIFB1 (Bactericidal/Permeability-Increasing Fold-Containing Family B Member 1), a key airway defense protein, is highly expressed in the early homeostatic steps but undergoes a sharp decline as the trajectory advances, signaling a failure of innate lung immunity.
- **Senescence and exhaustion:** CHEK2, (Checkpoint Kinase 2), a driver of cellular senescence and DNA damage response, remains inactive through the early and middle steps and increases only as the trajectory approaches the GOLD 4 centroid, consistent with terminal cellular exhaustion.
- **Systemic spillover:** The emergence of non-pulmonary markers, including the placenta-associated immunomodulator PSG9 and Urocortin-3 (UCN3), reflects the systemic spillover effect characteristic of plasma proteomics in severe COPD, likely driven by organ stress, aberrant tissue remodelling, and hypoxia-induced vascular leakage.

The functional annotations and standard nomenclature for the 20 variable proteins analyzed in the latent interpolation path are detailed in in Table C in S1 Appendix.

#### Topological stability against local graph destruction

To evaluate structural robustness under data starvation, we trained the model on independent random subsets, each comprising 50% of the training subjects (subsample A vs. B). Because LLE relies fundamentally on local nearest-neighbor graphs, withholding 50% of subjects typically shatters local connectivity. Despite this, Procrustes alignment of the independently trained manifolds (Fig 7) demonstrates consistent geometric convergence across all 10 iterations. Quantitative tracking (Fig 8) yielded a mean centroid shift of 5.199 *×* 10^−2^, representing a structural drift of less than 2% relative to the manifold diameter. The invariance of global clinical centroids despite destruction of local neighborhood topology confirms that the macroscopic branching structures are intrinsic properties of the disease biology rather than stochastic sampling artifacts.

**Fig 7.**
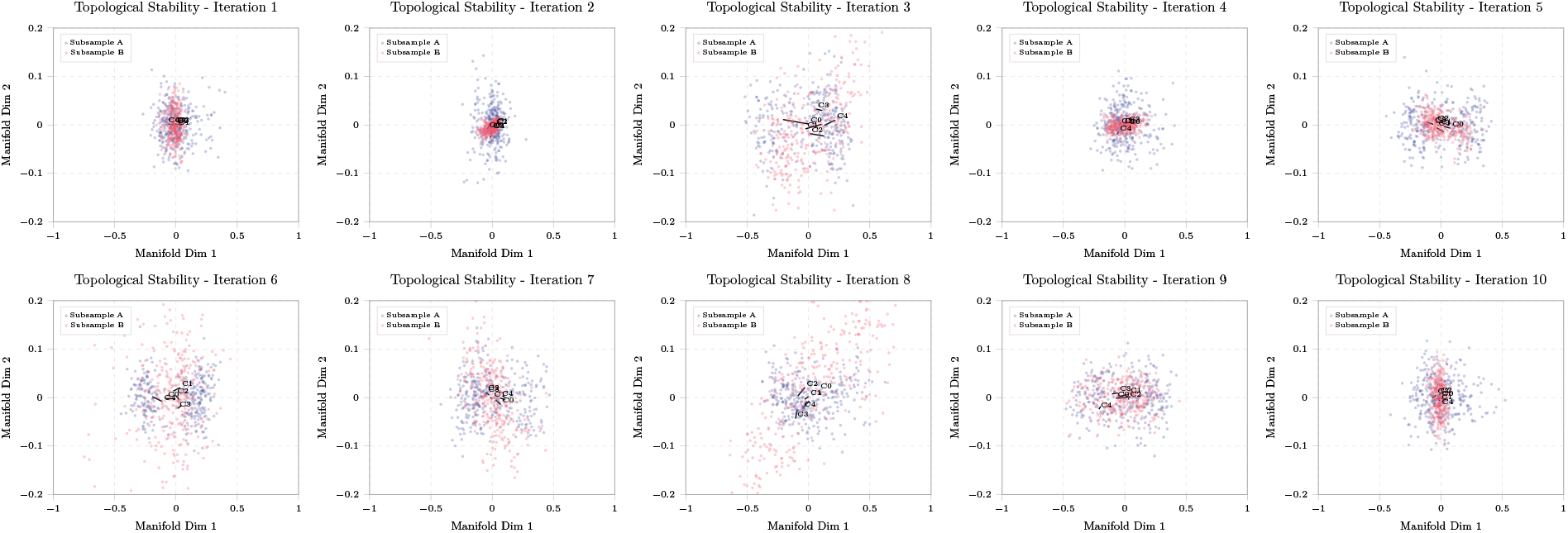
Topological stability under iterative subsampling. Independent manifolds generated from two cohort halves (subsample A and B) demonstrate consistent geometric structure across 10 random initializations. The repeated emergence of branching pillar-like structures supports the reproducibility of the learned manifold topology across cohort subsamples and indicates that these features are not driven by a single sample split or initialization.

**Fig 8.**
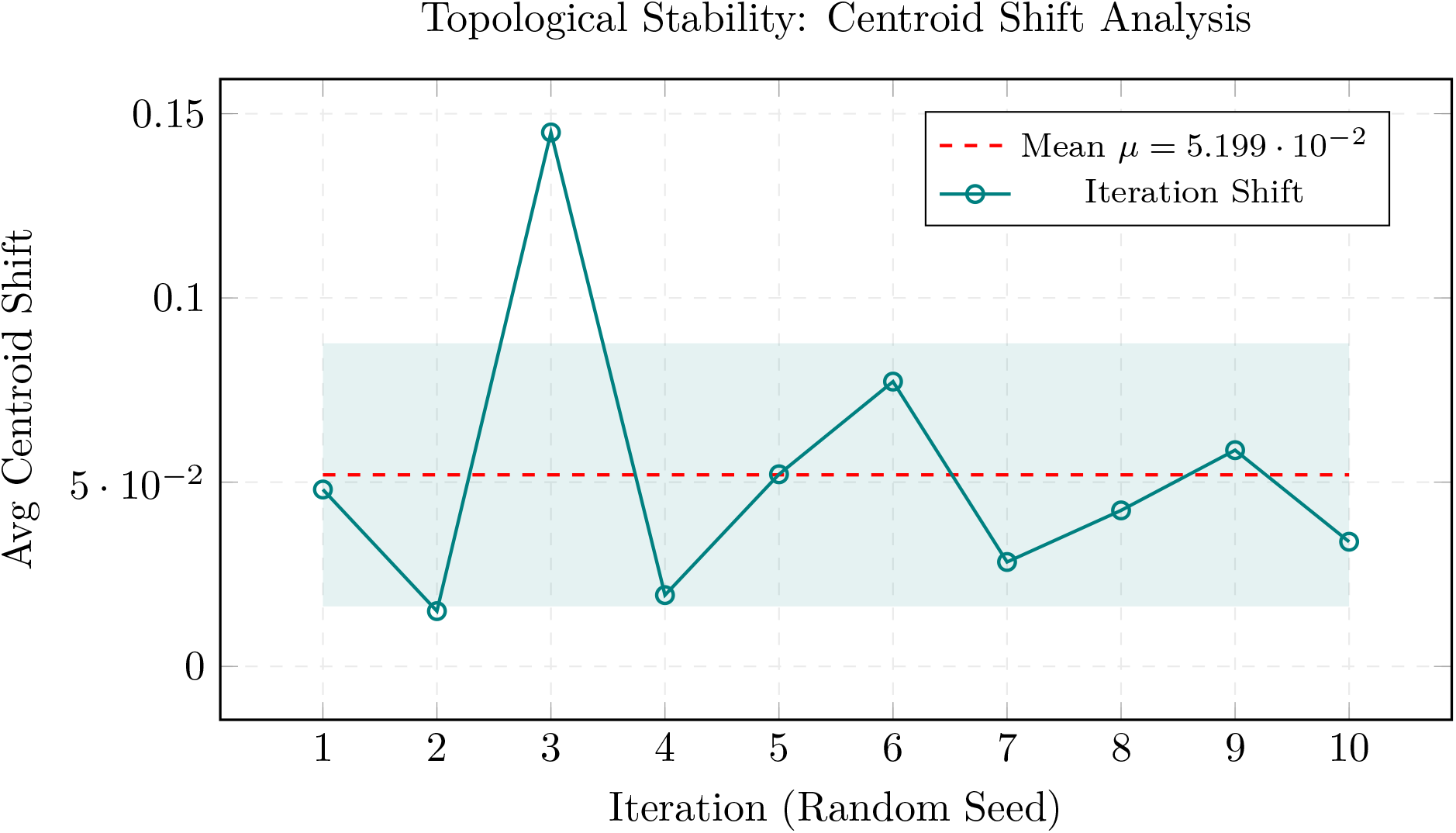
Quantitative analysis of centroid shift under subsampling. Mean centroid shift across 10 subsampling iterations (*µ* = 5.199 *×* 10^−2^), representing a structural drift of less than 2% relative to the manifold diameter. This low variance corroborates the stability of the reinforcement learning (RL) task-weighting policy across different training populations.

### Manifold interpretation and Disease Mechanism Discovery

#### Case study I: Static phenotyping and functional annotation

To demonstrate the necessity of the multi-task architecture, we compared the biological coherence of three distinct manifold configurations: (A) Supervised (LLE + classification), (B) Unsupervised (LLE + clustering), and (C) Hybrid MT-LLE (LLE + classification + clustering).

##### Supervised manifold (LLE + classification)

The classification manifold produced four usable regions containing 44, 24, 78, and 156 participants, respectively, and one excluded micro-cluster with *n* = 6 and a maximum *Z*-score of 4.9 (Fig A in S1 Appendix). Separation occurred primarily along a GOLD stage gradient, but at the cost of mechanistic resolution (Table D in S1 Appendix). Region 1 (*n* = 24, GOLD 0) showed a coherent complement signature, MASP1 (lectin complement initiation) alongside its B cell receptor CR2 (CD21), consistent with early systemic immune engagement in current smokers without obstruction.

Region 2 (*n* = 78), comprising participants with GOLD 0, appeared to represent a demographic grouping characterised by innate mucosal defence signals, with SPINK8 (serine protease inhibitor) and FZD8 (Wnt receptor) but no unifying mechanism. Region 0 (*n* = 44), which included participants across multiple GOLD stages, combined three distinct processes —immune exhaustion (TIGIT), bone remodelling (BGLAP), and mitochondrial metabolic stress (ETHE1, PHGDH)— within a single severity-defined cluster. Region 3 (*n* = 156; maximum *Z*-score = 0.24) represented the GOLD 2–3 centroid: its near-zero proteomic signal indicated that grouping patients by severity alone obscures within-stage heterogeneity.

##### Unsupervised manifold (LLE + clustering)

The clustering manifold produced three usable regions containing 61, 194, and 49 participants, respectively, and two excluded outlier clusters containing one and three participants, with maximum *Z*-scores greater than 9 (Fig B in S1 Appendix; exact proteomic profiles are detailed in Table E in S1 Appendix). The largest region (*n* = 194; 63% of the cohort; maximum *Z*-score = 0.20) combined participants from all GOLD stages into an undifferentiated mass, consistent with optimization for geometric compactness without clinical anchoring. The model also identified a pre-obstructive active-smoker group (Region 0; *n* = 61) and an older emphysematous former-smoker group (Region 2; *n* = 49). The latter was biologically distinctive: its top markers included grancalcin (GCA), a neutrophil calcium sensor; STMN4, which is involved in microtubule dynamics; and notably **ALOX15B**, or 15-lipoxygenase B, which contributes to the synthesis of pro-resolving lipid mediators. Elevated ALOX15B in the oldest, leanest, predominantly former-smoker group is consistent with a resolution-phase phenotype in which active inflammation has subsided while pro-resolving lipid biosynthesis persists as a chronic compensatory process.

##### Hybrid MT-LLE manifold (LLE + classification + clustering)

The Hybrid model produced four usable regions containing 72, 93, 119, and 23 participants, respectively, with only one excluded cluster: the same single-subject outlier appeared across all three models with an identical maximum *Z*-score of 17.4, supporting its interpretation as a data-level outlier independent of the training objective (Fig 9; Table F in S1 Appendix).

**Fig 9.**
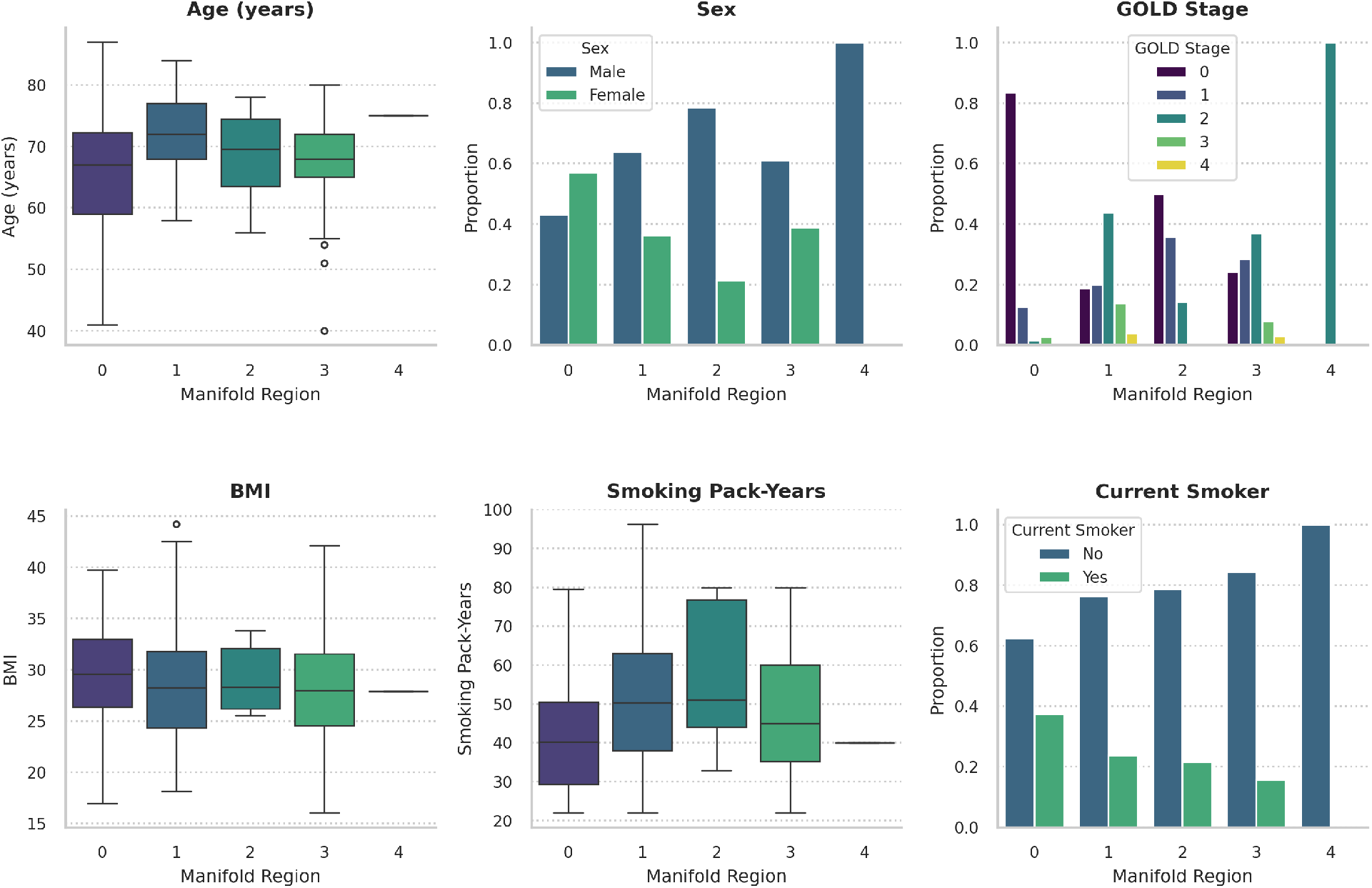
Clinical profiling of regions identified by the hybrid MT-LLE manifold. Clinical and demographic characteristics are summarized across manifold regions 0–4. (A) Age, (B) sex, (C) Global Initiative for Chronic Obstructive Lung Disease (GOLD) stage, (D) body mass index (BMI), (E) cumulative smoking exposure in pack-years, and (F) current smoking status. Boxplots show the median and interquartile range, with whiskers extending to 1.5 times the interquartile range; individual points denote observations beyond the whiskers. Bar plots show the within-region proportions for categorical variables. The profiles demonstrate distinct clinical identities across regions in demographic composition, disease severity, body composition, and smoking history. MT-LLE, multi-task locally linear embedding.

Region sizes were the most balanced among the three configurations, with a maximum cohort share of 39%, compared with 51% and 63% for the two single-objective models. Regions 0–2 preserved biologically coherent signals identified by both single-objective models (Table F in S1 Appendix). Region 0 (*n* = 72) recaptured the female pre-obstructive active-smoker phenotype. Region 1 (*n* = 93) distributed myeloid activation (TREML1), bone remodelling (BGLAP), and autophagy initiation (ATG7) across the severity gradient. Region 2 (*n* = 119) represented the GOLD 1–2 centroid, with uniformly low-magnitude proteomic signals (maximum *Z*-score = 0.165), confirming that this region captures the undifferentiated bulk of the cohort rather than a distinct molecular subtype.

##### Region 3: a Heavy-smoking stress-adaptation phenotype

Region 3 (*n* = 23) represents the distinctive finding of the hybrid model. In the LLE + classification manifold, these subjects were dispersed across GOLD-stage-defined regions with, only six extreme outliers co-located in one cluster(*n* = 6; *Z*-score = 4.9). In the LLE + clustering manifold, three of these participants formed a micro-cluster (*n* = 3; *Z*-score = 9.9). In neither single-objective model did the cluster reach the pre-specified minimum size threshold of *n* = 10 required for biological interpretation. In the hybrid model, the supervised loss constrained GOLD-stage dispersal while the clustering loss incorporated an additional 17 subjects, producing a stable region with a maximum *Z*-score of 1.20 that satisfied all predefined quality criteria.

Clinically, Region 3 was approximately 80% male and composed predominantly of former smokers (approximately 84%), with a minority of current smokers. Cumulative smoking burden was among the highest in the cohort, with a median exposure of approximately 50 pack-years.The GOLD distribution was skewed toward preserved spirometry, with approximately 50% of participants classified as GOLD 0 and 27% as GOLD 1, alongside representation across GOLD 2–4 (Fig 9). This pattern—heavy former smokers with predominantly preserved lung function—is consistent with a subgroup that has undergone durable physiological compensation rather than spirometric progression, despite extreme cumulative exposure.

Molecularly (Table F in S1 Appendix), the signature spanned four axes: a *glucocorticoid stress-response axis*, represented by KLF9, a cortisol-inducible Krüppel-like transcription factor, and HSD17B7, a bifunctional enzyme involved in androgen interconversion and cholesterol biosynthesis; an *epigenetic reprogramming axis*, represented by ACSS2, which supports histone acetylation through acetate-to-acetyl-CoA conversion under metabolic stress; an *endosomal and proteostatic adaptation axis*, represented by NEURL4, an E3 ubiquitin ligase, and KIAA0319L/AAVR, an endosomal trafficking receptor; and a *barrier repair axis*, represented by MARVELD2, a component of tricellular tight junctions, and DOCK9, a Cdc42 guanine nucleotide exchange factor. This chronic adaptive state was not recovered by either single-objective model, supporting the value of the multi-task architecture for resolving phenotypic structure that was not apparent under supervised or unsupervised dimensionality reduction alone.

#### Case study II: Trajectory inference and bifurcation analysis

Rather than a monolithic linear decline, the modeled trajectory (**Fig 10**) originated from a homeostatic root at node L, where GOLD 0 visits were concentrated, and diverged into two primary molecular progression axes. These branches were consistent with distinct progression patterns characterized by structural lung remodeling or sustained immune activation. Detailed branch-specific differential protein signatures are provided in S1 Appendix.

**Fig 10.**
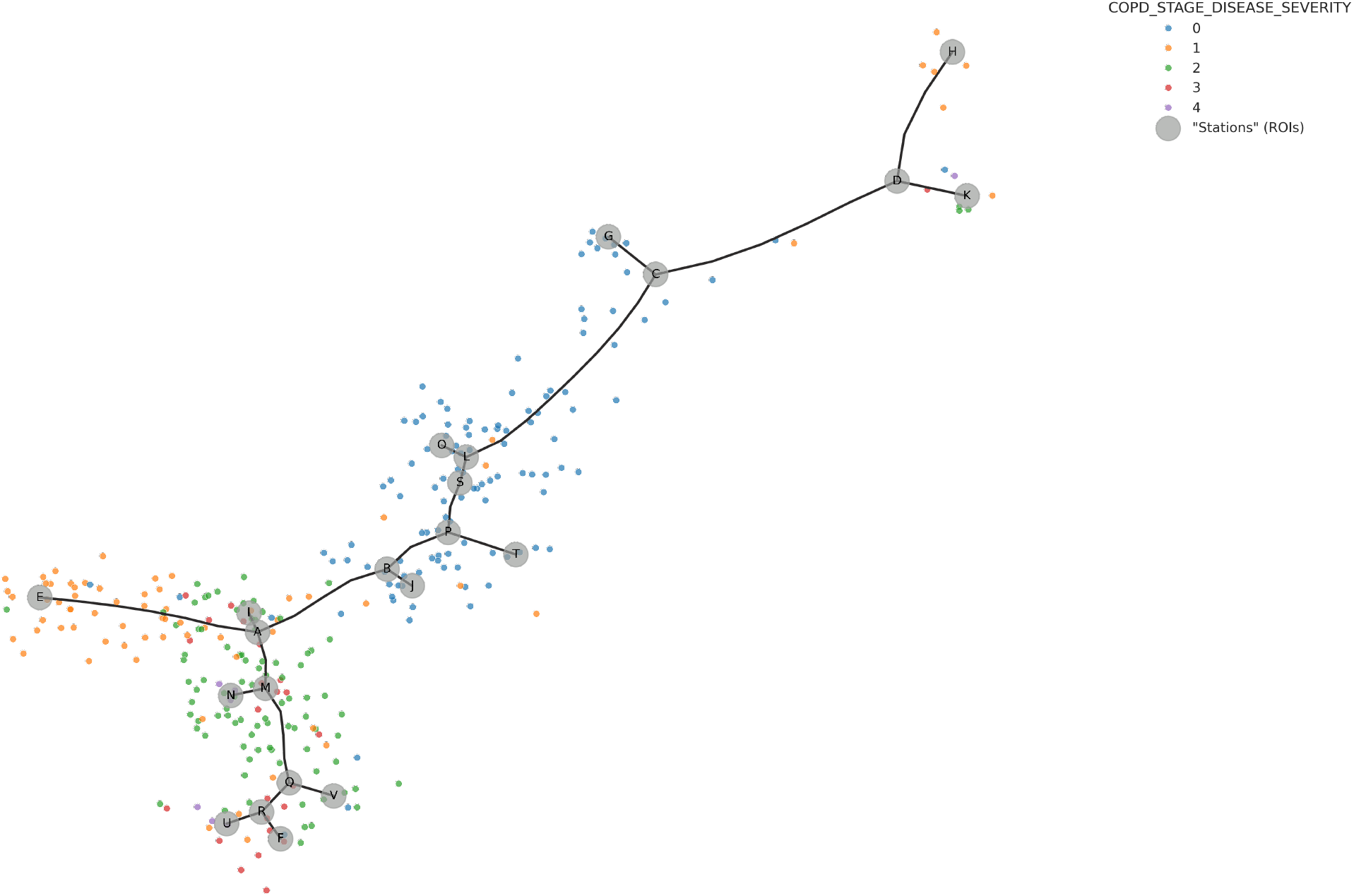
The MT-LLE trajectory skeleton. An elastic principal graph fitted to the multi-task locally linear embedding (MT-LLE) manifold reveals a branched disease-progression topology. Visits are colored by Global Initiative for Chronic Obstructive Lung Disease (GOLD) stage, ranging from 0 to 4. Labeled gray circles denote key manifold nodes. The trajectory originates at node L, where GOLD 0 visits are concentrated, and first diverges at node A.The two principal progression axes arising from node A are characterized in the main text; branch-specific differential protein signatures are provided in S1 Appendix.

##### Inflammatory fibrosis axis (path L → A → M → Q)

This trajectory was associated with severe structural progression and terminated in manifold regions enriched for GOLD 3 and GOLD 4 visits. The molecular signature differentiating this branch from the pan-immune activation axis was dominated by myeloperoxidase **(MPO)** and azurocidin **(AZU1)**, neutrophil granule proteins associated with degranulation and antimicrobial defense. This pattern suggested that neutrophil-driven oxidative and inflammatory activity may characterize this progression subtype. Relative to the tissue-maintenance branch (A → B), the differentiating profile shifted toward a fibrotic signature characterized by the profibrotic chemokine **CCL18**, which is associated with fibroblast activation and collagen deposition, and the basement-membrane organizer **FRAS1**. Together, these findings were consistent with a transition from neutrophilic inflammation toward progressive structural remodeling. Recurrence of EIF4A1 across multiple branch-wise comparisons further supported elevated translational activity as sustained feature of this axis (S1 Appendix), providing a plausible molecular route from early airway injury to advanced fibrotic disease.

##### Pan-immune activation axis (path L → A → I → E)

This axis diverged toward a state of broad immune activation and was associated with mild-to-moderate disease, particularly GOLD 1 and GOLD 2. The differentiating signature included **SLIT2**, a regulator of leukocyte chemotaxis that modulates monocyte and dendritic-cell migration through ROBO receptors, and the pro-inflammatory alarmin **S100A12**, which is released by activated granulocytes and amplifies inflammatory signaling. This branch also exhibited a strong adaptive immune signature. Upregulation of the major histocompatibility complex class II components **HLA-DQA1** and **HLA-DQB1** was consistent with active antigen presentation to helper T cells, while the immune-checkpoint modulator **CD276** (B7-H3) suggested concurrent regulation of T-cell responses.

Together, this pattern supported a progression subtype characterized by active antigen presentation and adaptive immune engagement rather than predominant structural remodeling, with potentially distinct therapeutic implications from the inflammatory fibrosis axis.

#### Case study III: Kinematic phenotyping of longitudinal progression

K-means clustering (*k* = 3; Fig. 11 and Fig. 12) identified three kinematic phenotypes. One cluster (Cluster 1, *n* = 4) fell below the minimum-participant threshold and was excluded from molecular interpretation. The two interpretable clusters are described below; detailed protein signatures and supplementary pseudotime trajectory figures are provided in section B of S1 Appendix.

**Fig 11.**
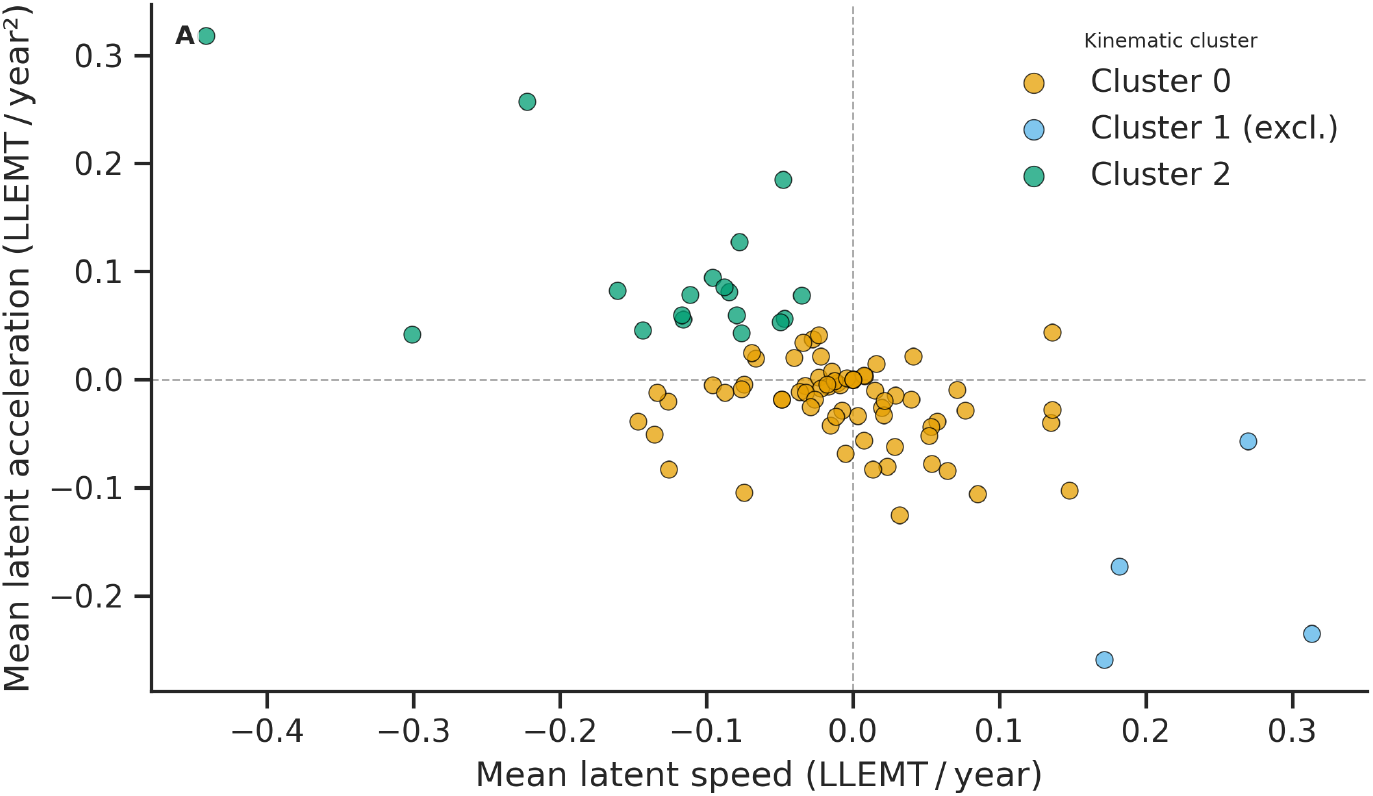
Kinematic phase space of disease progression. Subjects plotted by mean pseudo speed (*x*-axis; pseudotime change per calendar year, LLEMT yr^−1^) and mean pseudo acceleration (*y*-axis; change in speed per year, LLEMT yr^−2^). Cluster 0 (*n* = 70) occupies a broad region near the origin, consistent with limited net manifold displacement. Cluster 2 (*n* = 18) occupies the upper-left quadrant (negative mean speed, positive mean acceleration), indicating sustained movement toward lower-pseudotime manifold states at an increasing rate. Cluster 1 (*n* = 4) is shown for completeness but was excluded from molecular interpretation (minimum-participant criterion, *n <* 10).

**Fig 12.**
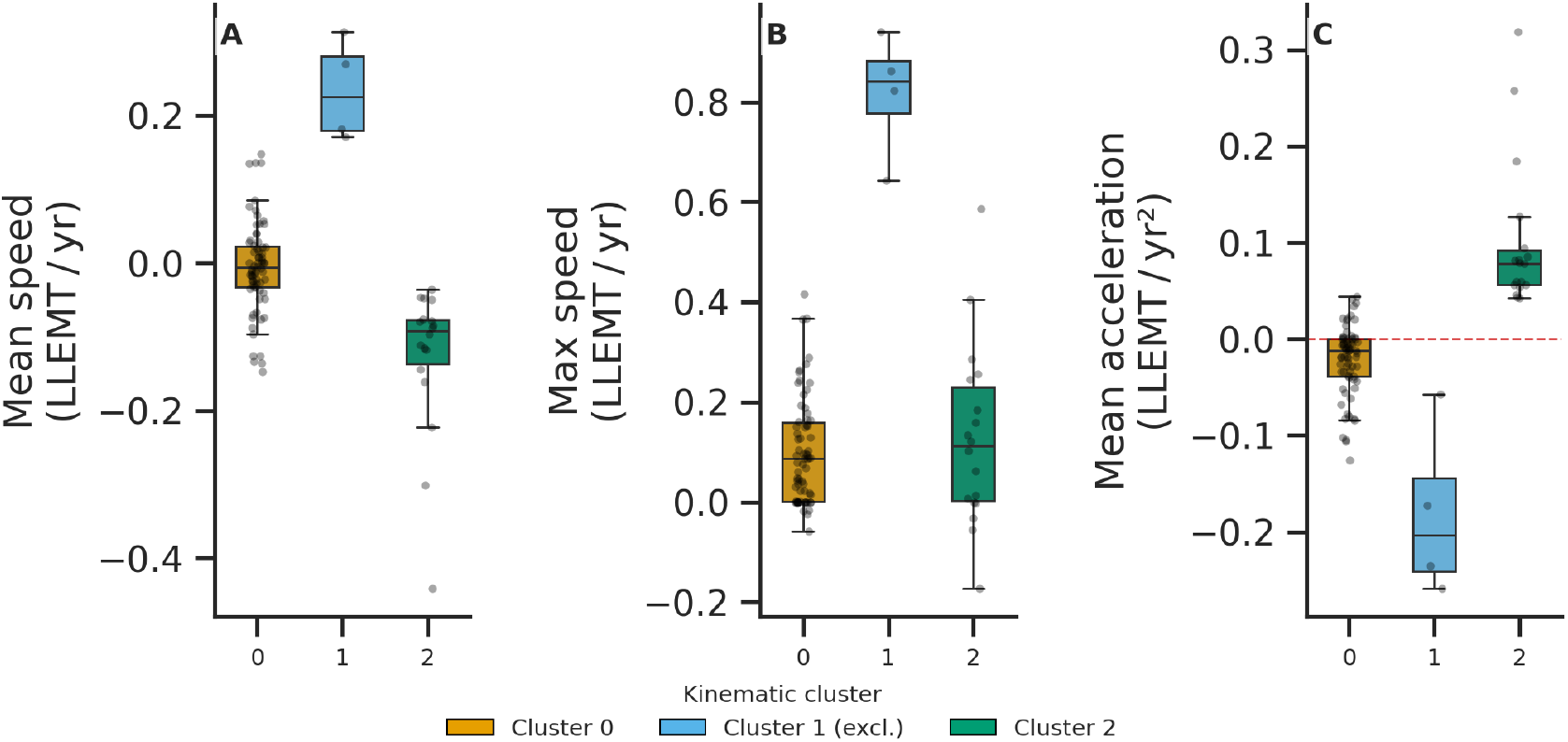
Kinematic metric distributions by cluster. Box plots of mean pseudo speed (A), maximum latent speed (B), and mean latent acceleration (C) stratified by kinematic cluster (Cluster 0, *n* = 70; Cluster 1, *n* = 4, excluded; Cluster 2, *n* = 18). Cluster 2 is the only cluster exhibiting both negative mean speed and positive mean acceleration simultaneously, distinguishing it kinematically from the near-homeostatic Cluster 0. Cluster 1 is shown for positional reference only; it was excluded from molecular interpretation due to insufficient participant count.

##### Cluster 0: Homeostatic majority (*n* = 70)

Subjects in Cluster 0 occupied a broad region near the origin of the kinematic phase space, with a mean latent speed of approximately − 0.02 LLEMT/yr and mean acceleration near zero (Fig. 11). Individual pseudotime trajectories were heterogeneous: 60% of subjects showed net pseudotime decline across follow-up, 29% showed net forward progression, and 11% showed a transient dip followed by recovery (S1 Appendix, Fig. D). The cluster mean declined modestly from approximately 1.20 to 1.02 over the observation period, consistent with a large, heterogeneous population with limited net directional molecular movement.

- **Clinical correlates:** GOLD stage trajectories were predominantly stable, with approximately 80% of visit intervals showing no GOLD stage change (S1 Appendix, Fig. F). Among subjects who were current smokers at baseline, 36.8% quit during follow-up (S1 Appendix, Fig. G).
- **Molecular signature:** Z-scores were uniformly low-magnitude (maximum 0.115), consistent with a large heterogeneous cluster whose mean profile does not strongly deviate from the cohort centroid. The highest-ranking proteins included **EGFL6** (epidermal growth factor-like protein 6; cell proliferation and adhesion), **TREML1** (triggering receptor expressed on myeloid cells-like 1; platelet and innate immune modulation), **LYG2** (lysozyme G-like protein 2; innate mucosal antibacterial defence), and **ATG7** (autophagy-related protein 7; autophagosome biogenesis, previously implicated in COPD epithelial cell death). Several proteins recurred from Case Study I: LYG2 and ATG7 from the Hybrid manifold Region 1, and ENO2 and MRPL2 from Region 2, suggesting that the near-homeostatic kinematic phenotype shares molecular character with the undifferentiated centroid regions identified by static phenotyping.
- **Interpretation:** Cluster 0 represents the large majority of the cohort in a state of limited net molecular change over the observed follow-up period. The low-magnitude, heterogeneous protein signature and modest pseudotime decline are consistent with this interpretation.

**Cluster 2: Retrograde-accelerating phenotype (***n* = 18 Cluster 2 represented a potential early-warning phenotype that may not be apparent from static molecular measurements alone.

Subjects in Cluster 2 occupied the upper-left quadrant of the kinematic phase space, with a mean pseudo speed of approximately − 0.12 LLEMT/yr and mean acceleration of approximately 0.08 LLEMT/yr^2^ (Fig. 11). This combination of negative mean speed and positive acceleration is kinematically distinctive: subjects were moving toward lower pseudotime values — toward molecular states characteristic of the healthier, root-adjacent region of the manifold — at an increasing rate. Pseudotime trajectory classification confirmed that 17 of 18 subjects (94%) showed sustained reversal, with the cluster mean declining sharply from approximately 1.37 at baseline to 0.93 at the final visit (S1 Appendix, Fig. E). This was the sharpest and most consistent directional pseudotime change observed in the cohort.

- **Clinical correlates:** GOLD stage trajectories were indistinguishable from Cluster 0, with approximately 80% of visit intervals showing no change (S1 Appendix, Fig. F). The molecular reversal captured by the manifold therefore preceded or was independent of any detectable change in GOLD classification. Strikingly, 66.7% of baseline smokers in Cluster 2 quit during follow-up, compared with 36.8% in Cluster 0 (S1 Appendix, Fig. G). Although the absolute number of baseline smokers in Cluster 2 was small (*n* ≈ 6), the approximately twofold higher cessation rate is consistent with smoking cessation as a contributor to the retrograde molecular trajectory. This is further supported by the biological character of the cluster’s molecular signature, described below.
- **Molecular signature:** Cluster 2 showed substantially higher Z-scores than Cluster 0 (range 0.38–0.63), consistent with a small, kinematically distinctive group whose molecular profile deviates markedly from the cohort mean. The highest-ranking proteins included **ADAMTS5** (ADAM metallopeptidase with thrombospondin motifs 5; extracellular matrix remodelling and cartilage homeostasis), **BDH2** (3-hydroxybutyrate dehydrogenase type 2, also annotated as DHRS6; ketone body and fatty acid metabolism), **ALOX15B** (arachidonate 15-lipoxygenase B; anti-inflammatory lipid mediator biosynthesis via the 15-LOX pathway, associated with resolution of inflammation), **TIGIT** (T-cell immunoreceptor with Ig and ITIM domains; immune checkpoint regulation, suppresses T-cell and NK-cell activity), and **UBE2W** (ubiquitin-conjugating enzyme E2 W; N-terminal ubiquitination and protein quality control). Collectively, this signature spans extracellular matrix remodelling, anti-inflammatory lipid signalling, immune regulation, and proteostasis — processes consistent with post-cessation inflammatory resolution rather than active disease progression.
- **Interpretation:** The combination of sustained retrograde molecular movement, twofold higher smoking cessation rate, and a protein signature enriched for anti-inflammatory and matrix-remodelling markers is consistent with Cluster 2 representing a subset of subjects undergoing molecular-level inflammatory resolution following smoking cessation. Crucially, this reversal is not accompanied by GOLD stage improvement, suggesting that the manifold captures an early or subclinical phase of biological recovery that conventional staging does not yet reflect. The small cluster size (*n* = 18) and the modest absolute number of quitters limit the strength of this interpretation, and prospective studies with larger cessation cohorts would be required to confirm it. Nonetheless, the capacity of the kinematic analysis to surface this phenotype — which is invisible to static profiling and to GOLD classification — illustrates the added resolution that incorporating longitudinal molecular dynamics can provide.

### Generalizability and multi-omics integration

#### Cross-modality and cross-cohort validation

While our primary analysis focused on the SPIROMICS proteomics dataset due to its high-resolution temporal sequencing, we performed identical quantitative evaluations across three additional cohort-modality datasets: SPIROMICS metabolomics, COPDGene proteomics, and COPDGene metabolomics. Detailed cohort- and modality-specific results are provided in Sections C, D, and E in S1 Appendix. Consistent with our primary findings, MT-LLE successfully optimized the trade-off between geometric fidelity and clinical utility across all datasets. Notably, while overall metric baselines shifted across modalities, for instance, metabolomics exhibited higher baseline reconstruction errors than proteomics due to the inherently higher biological noise and volatility of metabolite fluctuations, the relative ranking of model configurations remained invariant across all datasets. Across all molecular layers, the RL-phased MT-LLE consistently outperformed unsupervised baselines in downstream transferability, with MT-LLE KNN purity exceeding the best unsupervised baseline by 0.13-0.21 across datasets, confirming that the RL-phased multi-task optimization strategy generalizes consistently across independent cohorts and omics modalities.

#### Temporal-depth trade-off in multi-omics fusion

To investigate whether integrating multiple biological layers improved manifold resolution, we evaluated MT-LLE using paired proteomic and metabolomic data from SPIROMICS and COPDGene. We compared early fusion through feature concatenation with mid-fusion architecture described in subsection “Multi-omics extension: mid-Fusion architecture”, in which modality-specific encoders are aligned through cross-modal learning.

The mid-fusion approach mitigated differences in signal quality between the two modalities and outperformed both early fusion and the isolated metabolomics baseline. However, its overall performance did not exceed that of the single-omics proteomics model. In SPIROMICS, the best-performing mid-fusion configuration achieved a forecasting *F*_1_ score of 0.3480 *±* 0.013, compared with 0.3817 *±* 0.013 for the single-omics proteomics model trained using four visits. When the proteomics model was restricted to the same two-visit depth available for the paired dataset, its forecasting *F*_1_ score decreased to 0.3250 *±* 0.013. This substantially narrowed the performance gap and indicated that longitudinal depth contributed to the difference between the single- and multi-omics models.

This behavior arose from a fundamental constraint of the paired dataset: aligning the two modalities required truncating the longitudinal data from four clinical visits down to two. In the context of trajectory-aware manifold learning, this reduction in temporal depth substantially limited the model’s capacity to map the continuous, multi-stage molecular transitions of COPD progression, including the potential tipping points identified in the latent interpolation analysis (Fig 6). Consequently, although multi-omics integration provided a broader cross-sectional feature space, our results indicate that preserving longitudinal depth was more informative than expanding feature breadth for capturing the dynamics of disease progression in this setting.

## Conclusion

We presented MT-LLE, a multi-task manifold learning framework that learns clinically interpretable disease representations from longitudinal omics data, in which a patient’s position encodes their molecular state and severity and their movement encodes progression, by jointly optimizing geometric, supervised, temporal, and clustering objectives under the control of a reinforcement learning task scheduler. Across two independent COPD cohorts, MT-LLE resolved molecular heterogeneity not captured by standard clinical staging. The framework distinguished subjects with active molecular signatures despite preserved spirometry from stable subjects within the same clinical stage, and mapped molecularly distinct progression axes associated with neutrophil-driven inflammatory fibrosis and pan-immune activation. Kinematic trajectory analysis further stratified participants according to the dynamics of their molecular trajectories rather than their static states, identifying a subgroup whose molecular profiles changed at an accelerating rate relative to the rest of the cohort.

These findings support three principles for constructing interpretable disease manifolds. First, geometric fidelity and clinical utility may represent competing objectives; a useful manifold may need to accept modest geometric deformation in exchange for improved clinical coherence, and this trade-off should be explicitly controlled during model optimization. Second, the observed performance patterns suggest that unsupervised density-based organization may provide a useful scaffold for temporal forecasting beyond supervised label-based organization alone, because disease progression may follow continuous molecular pathways that do not align with discrete clinical boundaries. Third, the optimal task-weighting strategy appears to be non-stationary: objectives that conflict early in training may become complementary after a stable topological foundation has been established. Dynamically staged optimization may therefore provide a broadly applicable strategy for multi-task manifold learning in longitudinal biological data.

The learned manifold also suggests several potential avenues for clinical investigation. With prospective validation, the position of a newly enrolled participant on the manifold could help identify whether their molecular profile more closely follows an inflammatory fibrosis or pan-immune activation trajectory, potentially informing mechanism-based stratification beyond clinical stage alone. Kinematic clustering could similarly support prospective risk assessment by identifying participants with accelerating molecular trajectories for closer surveillance or enrollment in early-intervention studies. The pseudotime coordinate may also provide a continuous measure of molecular progression that complements discrete GOLD staging, particularly within the heterogeneous GOLD 0 population. These potential applications remain hypothesis-generating and require validation in prospective cohorts before clinical implementation.

Several directions remain for future work. This study evaluated early- and mid-fusion strategies under the constraint of complete matched cross-modal data. Future work will investigate fusion architectures that can accommodate visits observed in one modality but missing in another, a common feature of longitudinal observational cohorts. We also plan to evaluate MT-LLE in additional disease domains, including neurodegenerative and autoimmune conditions, where longitudinal molecular heterogeneity presents similar analytical challenges. Finally, the current reinforcement learning formulation learns a task-weighting policy during model training. An online-learning extension that updates the model and its task-weighting policy as additional patient visits become available could support longitudinally adaptive disease modeling as individual trajectories evolve over time.

## Supporting information

**S1 Appendix. Supplementary materials**. Detailed methods and extended analyses for MT-LLE, including the multi-omics mid-fusion architecture; extended SPIROMICS proteomics analyses (task-weighting ablation, biological characterization, static phenotyping, trajectory inference, and kinematic phenotyping); SPIROMICS metabolomics and multi-omics integration analyses; and COPDGene proteomics, metabolomics, and multi-omics integration analyses.

## Notes

### Competing Interest Statement

The authors have declared no competing interest.

### Funding

- NIH Common Fund: R01HL152735, R01HL137995, U01HL089897, U01HL089856

