## Supplementary Information for "MT-LLE: Multi-task locally linear embedding for interpretable disease modeling from longitudinal omics data"

### S1 Appendix. Supplementary materials

### Section A. Multi-omics mid-fusion architecture

While the core MT-LLE framework operates on a single omics modality, the architecture generalizes naturally to multiple modalities. In the experiments described in this paper, we instantiate this extension with two modalities, proteomics and metabolomics; however, the architecture places no constraints on the number of type of modality-specific inputs and can be applied to any set of high-dimensional omics layers with matched subjects and timepoints.

#### Modality-specific encoders

Two independent MT-LLE encoders are trained in parallel: encoder $E_{\phi_1}$ maps the first modality's input ($F_1$ features) to $z_i^{(1)} \in \mathbb{R}^d$, and encoder $E_{\phi_2}$ maps the second modality's input ($F_2$ features) to $z_i^{(2)} \in \mathbb{R}^d$. Each encoder maintains its own LLE reconstruction loss, ensuring that the neighborhood structure of each modality is independently preserved:

$$\mathcal{L}_{\text{LLE}} = \mathcal{L}_{\text{LLE}}^{(1)} + \mathcal{L}_{\text{LLE}}^{(2)} \tag{1}$$

#### Cross-modal neighborhood alignment

To enforce geometric consistency across the two modality-specific manifolds, we introduce a cross-modal alignment loss. The first modality's neighborhood structure serves as the reference anchor: samples that are neighbors in the first modality's embedding space are encouraged to also be close in the second modality's embedding space, and vice versa:

$$\mathcal{L}_{\text{align}} = \frac{1}{2}\left(\frac{1}{BK}\sum_i \sum_{j\in\mathcal{N}_i^{(1)}} \left\| z_i^{(2)} - z_j^{(2)} \right\|_2 + \frac{1}{BK}\sum_i \sum_{j\in\mathcal{N}_i^{(2)}} \left\| z_i^{(1)} - z_j^{(1)} \right\|_2 \right) \tag{2}$$

where $\mathcal{N}_i^{(1)}$ denotes the $K$ nearest neighbors of sample $i$ in the first modality's embedding space. A longitudinal cross-modal contrastive loss $\mathcal{L}_{\text{cross}}$ additionally pulls together representations of the same subject across modalities while treating different subjects as negatives, excluding same-subject pairs across different visits from the negative pool to preserve longitudinal structure. Both $\mathcal{L}_{\text{LLE}}$ and $\mathcal{L}_{\text{align}}$ are assigned fixed weights and are not controlled by the RL agent, ensuring that geometric validity is preserved independent of task scheduling decisions.

#### Bidirectional cross-attention fusion

The modality-specific embeddings are fused via a bidirectional cross-attention mechanism in which each modality attends to the other to produce context-enriched representations:

$$z^{(1\leftarrow 2)} = \text{CrossAttn}\left(Q = z^{(1)}, K = z^{(2)}, V = z^{(2)}\right) \quad z^{(2\leftarrow 1)} = \text{CrossAttn}\left(Q = z^{(2)}, K = z^{(1)}, V = z^{(1)}\right) \tag{3}$$

A learned sigmoid gate then blends the two context-enriched representations:

$$\alpha = \sigma\left(W_{\text{g}}\left[z^{(1\leftarrow 2)} \mid z^{(2\leftarrow 1)}\right]\right) \tag{4}$$

$$z_{\text{fused}} = \text{LayerNorm}\left(\alpha \odot z^{(1\leftarrow 2)} + (1-\alpha)\odot z^{(2\leftarrow 1)}\right) \tag{5}$$

The gate $\alpha \in (0,1)^d$ is detached from the gradient graph before being passed to the RL agent as a modality dominance signal, preventing the reward signal from flowing back through the fusion mechanism.

#### Extended RL agent

In the multi-omics setting, the RL agent's action and state spaces are extended relative to the single-omics formulation. For (M) modalities, the action vector becomes:

$$\boldsymbol{a}t = [w\text{pred}, w_{\text{cluster}}, w_{\text{traj}}, w_{\text{pheno}}, \alpha_1, \ldots, \alpha_M] \tag{6}$$

where $\alpha_1, \ldots, \alpha_M$ represent a soft modality blend overrides applied as a regularization signal on $z_{\text{fused}}$, allowing the agent to up-weight the more informative modality when manifold qualities diverge. In the experiments reported in this paper, $M = 2$, corresponding to proteomics and metabolomics. The state vector is extended to incorporate per-modality LLE losses, fusion gate summaries, and the cross-modal alignment loss. Task weights are sampled from a Dirichlet distribution parameterized by the policy output:

$$\boldsymbol{a}t \sim \text{Dirichlet}\left(10\pi\phi(s_t) + \epsilon\right) \tag{7}$$

The RL reward is extended to include a modality diversity bonus that penalizes fusion gate collapse (one modality dominating the fused representation):

$$R_t = (v_{\text{pred}} + v_{\text{NMI}}) - 0.5\mathcal{L}_{\text{align}} + 0.2H(\boldsymbol{\alpha}) \tag{8}$$

where $H(\boldsymbol{\alpha})$ is the entropy of the mean fusion gate values across the training set, incentivizing balanced utilization of both modalities.

#### Training protocol

A warmup period of 15 epochs is applied during which only the geometric losses ($\mathcal{L}_{\text{LLE}}$, $\mathcal{L}_{\text{align}}$, $\mathcal{L}_{\text{cross}}$) are active, the modality-specific manifolds to establish stable topological foundations before task-specific objectives are introduced. In the experiments reported in this paper, modality 1 corresponds to plasma proteomics and modality 2 corresponds to metabolomics.

### Section B. Extended analysis of the SPIROMICS proteomics dataset

#### Task weighting strategy ablation

Tables A and B report the complete quantitative results for the task weighting strategy ablation described in "Ablation of task weighting strategies" section of the main manuscript, providing exact mean ± standard deviation values across all geometric and downstream metrics for all five weighting strategies across 10 independent runs with distinct random seeds.

Table A additionally includes wall-clock time per epoch. The MT-LLE RL agent ($0.0535 \pm 0.0038$ s/epoch) adds only 0.003 seconds per epoch over the fastest comparator, Cosine Annealing ($0.0504 \pm 0.0041$ s/epoch), a 6% overhead, while GradNorm incurs the highest computational cost ($0.1010 \pm 0.0070$ s/epoch), an 89% increase relative to the RL agent despite underperforming it on every metric.

Table B confirms that the gap between Cosine Annealing and MT-LLE — 0.084 in KNN Purity, 0.062 in Classification F1, 0.057 in Forecasting F1, and 0.051 in Silhouette Score — demonstrates that the adaptive RL policy provides measurable value beyond phase ordering alone that a hand-crafted schedule cannot recover, with all margins over PCGrad supported by non-overlapping standard deviations across 10 independent random seeds.

#### Biological characterization of the latent interpolation path

Table C provides UniProt functional annotations for the top 20 most variable proteins identified along the GOLD 0 → GOLD 4 latent interpolation path. Proteins are listed in order of decreasing expression variability across the interpolation trajectory. UniProt annotations were retrieved June 2026.

**Table A. Intrinsic geometric quality (SPIROMICS proteomics).** Comparison of task weighting strategies against the MT-LLE RL agent on geometric measures and computational overhead.

| | **Intrinsic Geometric Measures** | | | |
|---|---|---|---|---|
| **Weighting Strategy** | **Reconstruction Error (↓)** *(MSE)* | **Local Neighbourhood Pres. (↑)** *(Ratio [0–1])* | **Global Structure Pres. (↑)** *(Spearman ρ [-1, 1])* | **Wall-clock (↓)** *(s/epoch)* |
| Static Uniform | 0.1250 ± 0.0085 | 0.3250 ± 0.0095 | 0.6850 ± 0.0155 | 0.0561 ± 0.0046 |
| GradNorm | 0.1120 ± 0.0080 | 0.3480 ± 0.0085 | 0.7220 ± 0.0125 | 0.1010 ± 0.0070 |
| PCGrad | 0.1085 ± 0.0075 | 0.3550 ± 0.0090 | 0.7350 ± 0.0110 | 0.0819 ± 0.0054 |
| Cosine Annealing | 0.1185 ± 0.0092 | 0.3390 ± 0.0105 | 0.7050 ± 0.0140 | **0.0504 ± 0.0041** |
| **MT-LLE (RL Agent)** | **0.0915 ± 0.0062** | **0.3805 ± 0.0075** | **0.7612 ± 0.0088** | 0.0535 ± 0.0038 |

All values reported as mean ± std across 10 independent runs with distinct random seeds. All strategies use identical encoder architecture and loss functions; only the task-weight assignment mechanism differs.

**Table B. Manifold coherence and downstream task performance (SPIROMICS proteomics).** Comparison of task weighting strategies against the MT-LLE RL agent.

| | **Clinical Coherence** | **Downstream Task Performance** | | |
|---|---|---|---|---|
| **Weighting Strategy** | **KNN Purity** *(Ratio [0–1])* | **Classification** *(F1-Macro)* | **Clustering** *(Silhouette Score)* | **Forecasting** *(F1-Macro)* |
| Static Uniform | 0.5050 ± 0.0125 | 0.4600 ± 0.0140 | 0.2350 ± 0.0120 | 0.3150 ± 0.0145 |
| GradNorm | 0.5350 ± 0.0140 | 0.4850 ± 0.0150 | 0.2620 ± 0.0130 | 0.3450 ± 0.0160 |
| PCGrad | 0.5495 ± 0.0135 | 0.4985 ± 0.0145 | 0.2750 ± 0.0125 | 0.3580 ± 0.0150 |
| Cosine Annealing | 0.5220 ± 0.0155 | 0.4720 ± 0.0165 | 0.2480 ± 0.0145 | 0.3320 ± 0.0175 |
| **MT-LLE (RL Agent)** | **0.6055 ± 0.0118** | **0.5342 ± 0.0135** | **0.2985 ± 0.0105** | **0.3885 ± 0.0142** |

All values reported as mean ± std across 10 independent runs with distinct random seeds. All strategies use identical encoder architecture and loss functions.

**Table C. Functional annotations.** Top 20 variable proteins in the SPIROMICS proteomics latent interpolation path (GOLD 0 → GOLD 4). UniProt annotations retrieved June 2025.

| **Protein** | **Gene** | **UniProt** | **UniProt Function (abridged)** |
|---|---|---|---|
| Serine/threonine-protein kinase Chk2 | CHEK2 | O96017 | Serine/threonine kinase required for checkpoint-mediated cell cycle arrest, DNA repair activation, and apoptosis in response to DNA double-strand breaks; phosphorylates CDC25A/B/C to block cell cycle progression, BRCA2 to promote homologous recombination repair, and p53/TP53 to initiate apoptosis; tumour suppressor whose absence contributes to chromosomal instability; driver of cellular senescence under chronic genotoxic stress. |
| Urocortin-3 | UCN3 | Q969E3 | Neuropeptide hormone central to whole-body stress adaptation; released by the hypothalamus in response to physical or psychological stress; acts via CRH receptor CRHR2 to regulate energy homeostasis, reduce food intake, improve insulin sensitivity, and manage blood glucose levels; does not activate the HPA axis to promote ACTH production, distinguishing it from CRH and UCN. |
| Cystatin-SN | CST1 | P01037 | Type 2 cystatin cysteine protease inhibitor; potent inhibitor of papain and dipeptidyl peptidase I; secreted in saliva, tears, and mucosal fluids; contributes to protease-antiprotease balance at mucosal surfaces. |
| Carcinoembryonic antigen-related cell adhesion molecule 7† | CEACAM7 | Q14002 | Member of the CEA family of the immunoglobulin superfamily; no functional annotation currently available in UniProtKB; expressed in gastrointestinal and respiratory epithelium. |
| Protein FAM241B (C10orf35) | FAM241B | Q96D05 | Also designated C10orf35; may play a role in lysosome homeostasis and membrane trafficking. |
| Large ribosomal subunit protein eL38† | RPL38 | P63173 | Component of the large 60S ribosomal subunit; required for IRES-mediated translation of a specific subset of developmental Hox mRNAs; selectively regulates translational output rather than global protein synthesis. |
| Ubiquitin carboxyl-terminal hydrolase 21 | USP21 | Q9UK80 | Deubiquitinase that removes ubiquitin from histone H2A, relieving epigenetic transcriptional repression and enabling di- and trimethylation of histone H3 at Lys-4; also deubiquitinates BAZ2A/TIP5 and ribosomal proteins RPS10/eS10 and RPS20/uS10; regulates transcriptional initiation and ribosome quality control. |
| Endoplasmic reticulum aminopeptidase 1 | ERAP1 | Q9NZ08 | Aminopeptidase in the ER lumen that trims peptide precursors to the 9-mer length required for MHC class I presentation; preferentially processes substrates 9–16 residues long with hydrophobic C-termini; key regulator of adaptive immune responses; may regulate blood pressure via angiotensin II inactivation. |

| Protein | Gene | UniProt | UniProt Function (abridged) |
|---|---|---|---|
| Left-right determination factor 2 | LEFTY2 | O00292 | TGF-$\beta$ superfamily member required for left-right asymmetry determination of organ systems; inhibits nodal signalling; may play a role in endometrial bleeding. |
| Nuclear pore complex protein Nup98–Nup96 | NUP98 | P52948 | Structural component of the nuclear pore complex (NPC); mediates bidirectional nucleocytoplasmic transport together with NUP96; may anchor NUP153 and TPR to the NPC; in cooperation with DHX9, activates transcription and alternative splicing of a subset of genes; frequently rearranged in haematological malignancies. |
| Pregnancy-specific beta-1-glycoprotein 9 | PSG9 | Q00887 | Binds the small latent TGF-$\beta$1 complex and activates TGF-$\beta$1; stimulates FoxP3 expression in naive $CD4^+$ T-cells, increasing regulatory T-cell numbers; induces TGF-$\beta$1 secretion from macrophages; may reduce pro-inflammatory cytokine expression by T-cells; plasma elevation likely reflects systemic immunomodulatory spillover. |
| BPI fold-containing family B member 1 | BPIFB1 | Q8TDL5 | Innate defense protein expressed in airway epithelium and submucosal glands; binds bacterial lipopolysaccharide (LPS) and modulates cellular responses to LPS; may play a role in innate immunity in mouth, nose, and lungs; downregulated in COPD and cystic fibrosis, consistent with failure of innate airway immunity. |
| Gamma-enolase | ENO2 | P09104 | Glycolytic enzyme catalysing the conversion of 2-phosphoglycerate to phosphoenolpyruvate; neuroendocrine isoform with neurotrophic and neuroprotective properties; binds cultured neocortical neurons in a calcium-dependent manner; serum biomarker of neuronal injury and neuroendocrine differentiation of the airway epithelium. |
| Thioredoxin-related transmembrane protein 1 | TMX1 | Q9H3N1 | ER-resident oxidoreductase that catalyses dithiol-disulfide exchange reactions; inhibits the alternative triglyceride biosynthesis pathway via TMEM68/DIESL; mediates disulfide bond formation in transmembrane proteins; involved in ER-associated degradation (ERAD) and regulation of ER-mitochondria contact sites and $Ca^{2+}$ flux; also designated TXNDC1. |
| PDZ domain-containing protein GIPC1[†] | GIPC1 | O14908 | Scaffold protein containing a central PDZ domain; regulates G protein-linked signalling via interaction with RGS-GAIP; involved in vesicular trafficking and receptor internalisation. |

| Protein | Gene | UniProt | UniProt Function (abridged) |
|---|---|---|---|
| Ubiquitin-conjugating enzyme E2 W | UBE2W | Q96B02 | E2 ubiquitin-conjugating enzyme that specifically monoubiquitinates the N-terminus of disordered substrate proteins including ATXN3, MAPT/TAU, and STUB1/CHIP; mediates monoubiquitination of FANCD2 in the DNA damage response; involved in protein quality control and degradation of misfolded chaperone substrates. |
| Disintegrin and metalloproteinase domain-containing protein 7 | ADAM7 | Q9H2U9 | Non-catalytic metalloprotease-like protein required for normal male fertility via maintenance of caput epididymis epithelial morphology and lumen structure; plays a role in sperm motility, flagella morphology, and tyrosine phosphorylation during capacitation; also expressed in airway epithelium and regulated by androgens. |
| Rho GTPase-activating protein 1 | ARHGAP1 | Q07960 | GTPase-activating protein for Rho, Rac, and Cdc42; converts active GTP-bound Rho GTPases to the inactive GDP-bound state; preferred substrate is Cdc42; regulates actin cytoskeleton dynamics and cell polarity. |
| Bcl-2-modifying factor† | BMF | Q96LC9 | Pro-apoptotic BH3-only protein of the Bcl-2 family, with isoform 1 as the principal apoptotic initiator; sequesters anti-apoptotic Bcl-2 family members; mediates anoikis and oxidative-stress-induced apoptosis. |
| Lysozyme g-like protein 2† | LYG2 | Q86SG7 | Probable antibacterial protein of the innate mucosal immune system; expressed in secretory cells of the respiratory tract; contributes to airway antimicrobial defence. |

† UniProt functional annotation is sparse or absent for this entry; the abridged description incorporates established literature consistent with the annotation approach used in Tables F–H.

#### Case study I: Static phenotyping and functional annotation

Fig A and Fig B show the clinical and demographic profiles of regions identified by the supervised (LLE + classification) and unsupervised (LLE + clustering) manifolds, respectively, providing the supporting clinical context for the phenotypic descriptions in the main manuscript. Regions excluded by the pre-specified quality criteria are displayed but not interpreted.

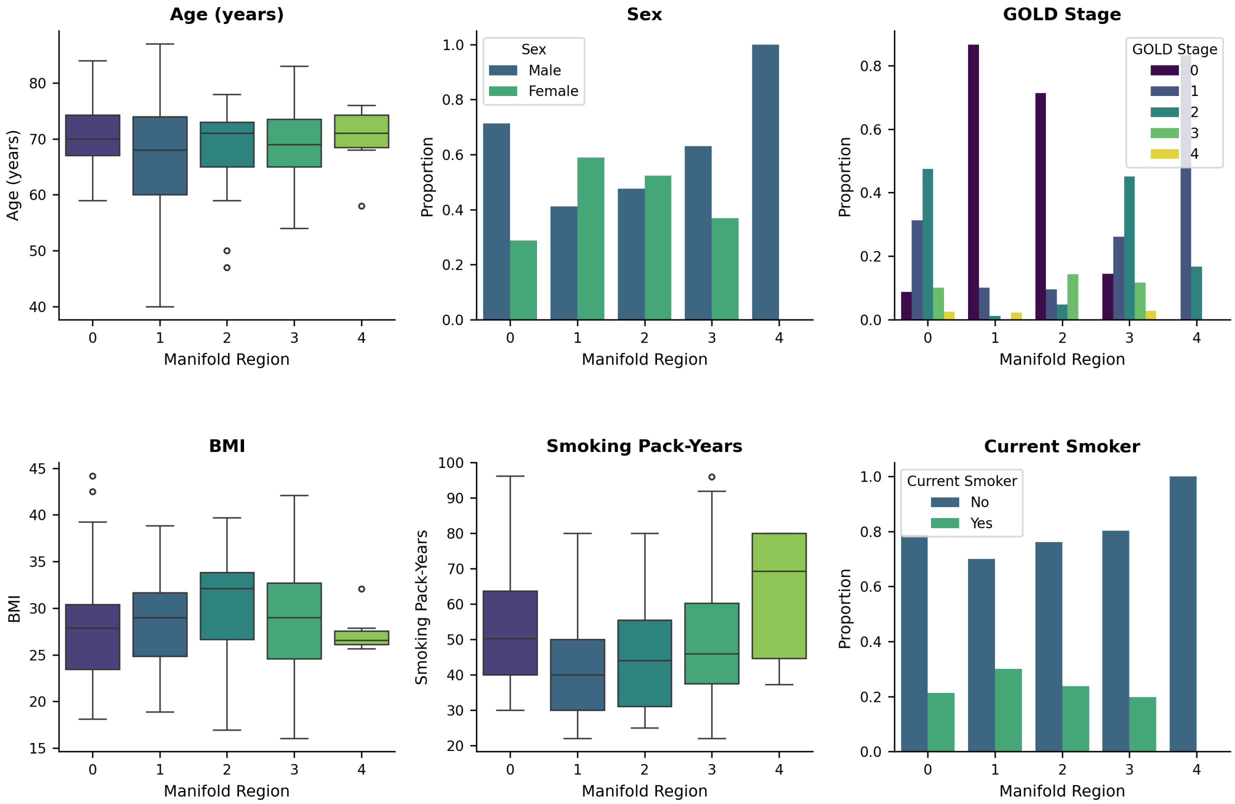


**Fig A. Clinical profiling of regions identified by the supervised manifold (LLE + classification).** Clinical and demographic characteristics are summarised across manifold regions 0–4. (A) Age, (B) sex, (C) Global Initiative for Chronic Obstructive Lung Disease (GOLD) stage, (D) body mass index (BMI), (E) cumulative smoking exposure in pack-years, and (F) current smoking status. Boxplots show the median and interquartile range, with whiskers extending to 1.5 times the interquartile range; individual points denote observations beyond the whiskers. Bar plots show the within-region proportions for categorical variables. Regions 0–3 are usable; Region 4 ($n = 6$; maximum $Z$-score = 4.9) was excluded by the pre-specified minimum size criterion and is shown for completeness but not narratively interpreted.

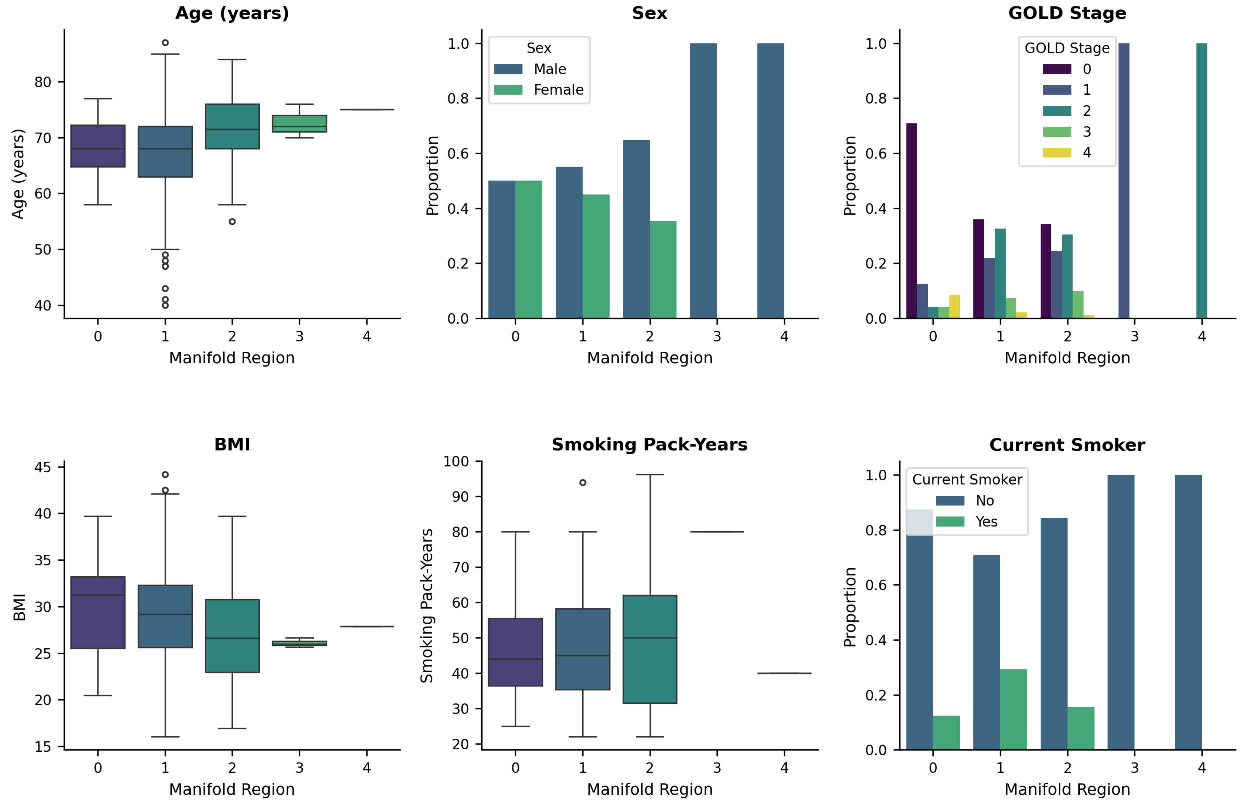


**Fig B. Clinical profiling of regions identified by the unsupervised manifold (LLE + clustering).** Clinical and demographic characteristics are summarised across manifold regions 0–4. (A) Age, (B) sex, (C) Global Initiative for Chronic Obstructive Lung Disease (GOLD) stage, (D) body mass index (BMI), (E) cumulative smoking exposure in pack-years, and (F) current smoking status. Boxplots show the median and interquartile range, with whiskers extending to 1.5 times the interquartile range; individual points denote observations beyond the whiskers. Bar plots show the within-region proportions for categorical variables. Regions 0–2 are usable; Regions 3 ($n = 1$) and 4 ($n = 3$), both with maximum $Z$-scores exceeding 9, were excluded by the pre-specified size and anomaly thresholds and are shown for completeness but not narratively interpreted.

Tables D, E, and F report the top-10 differentially abundant proteins per usable region for the supervised, unsupervised, and hybrid manifold configurations, respectively. For each region, proteins are ranked by mean Z-score computed as (region mean − global mean) / global SD on the original high-dimensional feature space. The functional annotations were retrieved from UniProtKB (June 2026) and regions excluded by the pre-specified quality criteria are omitted from all three tables.

**Table D. Regional proteomic profiles.** Top-10 differentially abundant proteins per usable region in the LLE + Classification manifold.

| Ensembl ID | Symbol | Z | UniProt Function (abridged) |
|---|---|---|---|
| ***Region 0 ($n = 44$, max Z = 0.65) — Mixed GOLD; immune exhaustion, bone remodelling, metabolic reprogramming*** | | | |
| ENSG00000211898 | IGHD | 0.648 | Constant region of the IgD heavy chain; mediates IgD-class antibody effector functions in B cell activation. |
| ENSG00000198759 | EGFL6 | 0.562 | EGF-like domain protein; promotes endothelial cell proliferation and angiogenesis; expressed in vascular stroma and foetal tissues. |
| ENSG00000125999 | BGLAP | 0.549 | Osteocalcin; bone matrix protein secreted by osteoblasts; regulates mineralisation, energy metabolism, and insulin secretion; elevated in COPD-associated osteoporosis. |
| ENSG00000105755 | ETHE1 | 0.520 | Mitochondrial persulfide dioxygenase; catabolises $H_2S$ via the sulfide oxidation pathway; loss-of-function causes ethylmalonic encephalopathy. |
| ENSG00000092621 | PHGDH | 0.516 | Phosphoglycerate dehydrogenase; catalyses the first step of L-serine biosynthesis from 3-phosphoglycerate; upregulated in oxidative stress and one-carbon metabolic reprogramming. |
| ENSG00000181847 | TIGIT | 0.496 | T cell immunoreceptor with Ig and ITIM domains; inhibitory checkpoint suppressing T and NK cell activation upon binding poliovirus receptor (PVR). |
| ENSG00000120324 | PCDHB10 | 0.462 | Protocadherin beta-10; calcium-dependent cell-cell adhesion glycoprotein expressed in neurons; regulates axonal connectivity. |
| ENSG00000161911 | TREML1 | 0.432 | Triggering receptor expressed on myeloid cells-like 1; activates platelet and myeloid cell signalling; modulates inflammatory responses. |
| ENSG00000004059 | ARF5 | 0.424 | ADP ribosylation factor 5; GTPase regulating Golgi-to-endosome vesicular transport and organelle membrane dynamics. |
| ENSG00000173698 | ADGRG2 | 0.423 | Adhesion G-protein coupled receptor G2; receptor for steroid hormones; regulates cAMP levels in efferent ductules; involved in male reproductive tract function. |
| ***Region 1 ($n = 24$, max Z = 0.93) — GOLD 0; lectin complement pathway activation*** | | | |
| ENSG00000103043 | VAC14 | 0.928 | Scaffold component of the PIKfyve lipid kinase complex; regulates endosomal PI(3,5)$P_2$ levels and lysosomal membrane trafficking. |
| ENSG00000069206 | ADAM7 | 0.927 | ADAM metallopeptidase domain 7; membrane-anchored ECM sheddase; expressed in testicular epithelium during sperm maturation; also found in airway epithelium. |

| Ensembl ID | Symbol | Z | UniProt Function (abridged) |
|---|---|---|---|
| ENSG00000137947 | GTF2B | 0.869 | General transcription factor IIB; positions RNA polymerase II at the transcription start site; required for basal and activated Pol II transcription initiation. |
| ENSG00000088387 | DOCK9 | 0.861 | Dedicator of cytokinesis 9; Cdc42-specific guanine nucleotide exchange factor; regulates actin cytoskeleton, filopodia formation, and cellular polarity. |
| ENSG00000127241 | MASP1 | 0.859 | Mannan-binding lectin serine protease 1; initiates the lectin pathway of complement by activating MASP-2; cleaves C2, C3, fibrinogen, and factor XIII. |
| ENSG00000170632 | ARMC10 | 0.819 | Armadillo repeat containing 10; regulates mitochondrial network dynamics; interacts with proteasomal subunits; ubiquitously expressed. |
| ENSG00000113387 | SUB1 | 0.818 | Transcriptional coactivator PC4; facilitates RNA Pol II assembly and transcription stimulation; involved in nucleotide excision repair. |
| ENSG00000131069 | ACSS2 | 0.817 | Acyl-CoA synthetase short chain family member 2; converts acetate to acetyl-CoA for histone acetylation and lipid synthesis; upregulated under metabolic stress and nutrient limitation. |
| ENSG00000152939 | MARVELD2 | 0.794 | Marvel domain-containing protein 2; plays a role in the formation of tricellular tight junctions between epithelial cells; required for cochlear hair cell survival and hearing. |
| ENSG00000117322 | CR2 | 0.780 | Complement receptor 2 (CD21); B cell co-receptor binding C3d and Epstein-Barr virus gp350; lowers B cell activation threshold via BCR co-receptor complex with CD19 and CD81. |
| ***Region 2 ($n = 78$, max Z = 0.51) — GOLD 0; innate mucosal defence and barrier signals*** | | | |
| ENSG00000079257 | LXN | 0.513 | Latexin; endogenous serine carboxypeptidase inhibitor; regulates haematopoietic stem cell pool size; anti-inflammatory and putative tumour suppressor. |
| ENSG00000129221 | AIPL1 | 0.457 | Aryl hydrocarbon receptor interacting protein-like 1; co-chaperone in the HSP90 complex; essential for photoreceptor viability and farnesylated protein stability. |
| ENSG00000198873 | GRK5 | 0.424 | G protein-coupled receptor kinase 5; serine/threonine kinase preferentially phosphorylating agonist-occupied GPCRs; regulates $\beta$-adrenergic receptor desensitisation and hypertrophic signalling. |
| ENSG00000113369 | ARRDC3 | 0.341 | Arrestin domain containing 3; negative regulator of $\beta$-adrenergic signalling; promotes receptor ubiquitination; regulates adiposity and GPCR internalisation. |
| ENSG00000203970 | DEFB110 | 0.339 | Defensin beta 110; antimicrobial peptide with antibacterial activity; component of mucosal innate immune defence. |
| ENSG00000229453 | SPINK8 | 0.338 | Serine protease inhibitor Kazal type 8; probable serine protease inhibitor; likely involved in airway protease–antiprotease balance. |

| Ensembl ID | Symbol | Z | UniProt Function (abridged) |
|---|---|---|---|
| ENSG00000177283 | FZD8 | 0.335 | Frizzled-8; receptor for Wnt proteins; component of the Wnt–Fzd–LRP5–LRP6 signalling complex; regulates cell fate determination, proliferation, and polarity. |
| ENSG00000174992 | ZG16 | 0.325 | Zymogen granule protein 16; probable lectin involved in protein trafficking and secretory pathway organisation in exocrine glands and mucosal surfaces. |
| ENSG00000104343 | UBE2W | 0.300 | Ubiquitin conjugating enzyme E2 W; accepts ubiquitin from the E1 complex and transfers it to substrates; involved in N-terminal ubiquitination. |
| ENSG00000197191 | CYSRT1 | 0.299 | Cysteine-rich tail protein 1; antimicrobial epidermal protein located in the cornified envelope; interacts with late cornified envelope proteins; involved in establishment of epithelial barrier integrity and host-pathogen mucosal defence. |
| ***Region 3 ($n = 156$, max Z = 0.24) — GOLD 2–3 centroid; broad low-magnitude disease signals*** | | | |
| ENSG00000115170 | ACVR1 | 0.242 | Activin receptor type 1 (ALK2); BMP type I receptor serine/threonine kinase; transduces TGF-$\beta$/BMP signals; regulates osteogenesis and vascular calcification; gain-of-function mutations cause fibrodysplasia ossificans progressiva. |
| ENSG00000127951 | FGL2 | 0.232 | Fibrinogen-like protein 2; may play a role in physiologic lymphocyte functions at mucosal surfaces; fibroleukin activity; expressed in macrophages and T cells. |
| ENSG00000242252 | BGLAP | 0.230 | Osteocalcin (alternate locus); bone matrix protein secreted by osteoblasts; regulates bone mineralisation and energy metabolism; also functions as a hormone. |
| ENSG00000178473 | UCN3 | 0.225 | Urocortin 3; neuropeptide of the CRF family; suppresses food intake, delays gastric emptying, and regulates HPA-axis stress responses. |
| ENSG00000172809 | RPL38 | 0.179 | Ribosomal protein L38; component of the large (60S) ribosomal subunit; required for internal ribosome entry site-mediated translation of a subset of Hox mRNAs. |
| ENSG00000166311 | SMPD1 | 0.141 | Sphingomyelin phosphodiesterase 1 (acid sphingomyelinase); converts sphingomyelin to ceramide; implicated in cigarette smoke-induced ceramide-mediated apoptosis in lung epithelium. |
| ENSG00000151651 | ADAM8 | 0.136 | ADAM metallopeptidase domain 8; involved in extravasation of leukocytes during inflammation; cleaves EGFR and L-selectin; elevated in COPD airways. |
| ENSG00000037757 | MRI1 | 0.117 | Methylthioribose-1-phosphate isomerase 1; catalyses the interconversion of methylthioribose-1-phosphate in the methionine salvage pathway. |
| ENSG00000104081 | BMF | 0.087 | Bcl-2 modifying factor; pro-apoptotic BH3-only protein; may regulate apoptosis by sequestering Bcl-2 family members; implicated in anoikis and oxidative-stress-induced cell death. |
| ENSG00000185674 | LYG2 | 0.081 | Lysozyme G2; potent antibacterial protein involved in innate mucosal immune defence; expressed in secretory cells of the respiratory and gastrointestinal tracts. |

**Table E. Regional proteomic profiles.** Top-10 differentially abundant proteins per usable region in the LLE + Clustering manifold.

| **Ensembl ID** | **Symbol** | **Z** | **UniProt Function (abridged)** |
|---|---|---|---|
| ***Region 0 ($n = 61$, max Z = 0.82) — Pre-obstructive; innate mucosal defence and barrier signals*** | | | |
| ENSG00000129221 | AIPL1 | 0.816 | Aryl hydrocarbon receptor interacting protein-like 1; co-chaperone in HSP90 complex required for photoreceptor viability; interacts with farnesylated proteins and FKBP51/52. |
| ENSG00000079257 | LXN | 0.681 | Latexin; endogenous serine carboxypeptidase inhibitor; regulates haematopoietic stem cell number; suppresses tumourigenesis and inflammation. |
| ENSG00000203970 | DEFB110 | 0.661 | Defensin beta 110; antimicrobial peptide of the $\beta$-defensin family; mediates innate mucosal immunity against bacteria. |
| ENSG00000174992 | ZG16 | 0.649 | Zymogen granule protein 16; probable lectin involved in protein trafficking; may act as a linker between secretory granule membranes and mucin polymers at mucosal surfaces. |
| ENSG00000170632 | ARMC10 | 0.592 | Armadillo repeat containing 10; regulates mitochondrial dynamics; interacts with the proteasome; expressed ubiquitously. |
| ENSG00000069206 | ADAM7 | 0.573 | ADAM metallopeptidase 7; membrane-anchored ECM sheddase; also expressed in respiratory epithelium; regulated by androgens. |
| ENSG00000143258 | USP21 | 0.486 | Ubiquitin specific peptidase 21; deubiquitinates histone H2A, reversing an epigenetic mark for gene silencing; also deubiquitinates BRCA2 and PCNA. |
| ENSG00000229453 | SPINK8 | 0.478 | Serine protease inhibitor Kazal type 8; probable serine protease inhibitor implicated in mucosal protease–antiprotease balance. |
| ENSG00000103043 | VAC14 | 0.461 | Scaffold of the PIKfyve complex; regulates endosomal PI(3,5)$P_2$ and lysosomal membrane trafficking. |
| ENSG00000139921 | TMX1 | 0.402 | Thioredoxin-related transmembrane protein 1; oxidoreductase participating in protein disulphide bond formation in the endoplasmic reticulum; involved in ER-stress responses. |
| ***Region 1 ($n = 194$, max Z = 0.20) — Undifferentiated background; all GOLD stages; near-zero proteomic signal*** | | | |
| ENSG00000115170 | ACVR1 | 0.200 | Activin A receptor type 1 (ALK2); BMP/TGF-$\beta$ type I receptor; regulates osteogenesis, vascular calcification, and innate immune pathway activation. |
| ENSG00000242252 | BGLAP | 0.196 | Osteocalcin (alternate locus); bone matrix protein secreted by osteoblasts; regulates bone mineralisation and functions as a hormone. |
| ENSG00000127951 | FGL2 | 0.157 | Fibrinogen-like protein 2; involved in physiologic lymphocyte functions at mucosal surfaces; expressed in macrophages and regulatory T cells. |
| ENSG00000178473 | UCN3 | 0.145 | Urocortin 3; CRF-family neuropeptide; suppresses food intake, delays gastric emptying, and mediates HPA-axis stress responses. |

| Ensembl ID | Symbol | Z | UniProt Function (abridged) |
|---|---|---|---|
| ENSG00000172809 | RPL38 | 0.132 | Ribosomal protein L38; component of the 60S ribosomal subunit; required for IRES-mediated translation of a subset of developmental Hox mRNAs. |
| ENSG00000170323 | FABP4 | 0.116 | Fatty acid binding protein 4 (A-FABP); intracellular lipid transporter in adipocytes and macrophages; regulates lipid metabolism, inflammatory signalling, and insulin resistance. |
| ENSG00000121769 | FABP3 | 0.109 | Fatty acid binding protein 3 (H-FABP); transports long-chain fatty acids in muscle and myocardium; serum biomarker of myocardial injury. |
| ENSG00000151651 | ADAM8 | 0.101 | ADAM metallopeptidase domain 8; involved in leukocyte extravasation; cleaves EGFR and L-selectin; elevated in COPD airways and associated with disease severity. |
| ENSG00000128965 | CHAC1 | 0.087 | ChaC glutathione-specific gamma-glutamylcyclotransferase 1; cleaves glutathione, reducing cellular antioxidant capacity; pro-apoptotic ER stress effector downstream of ATF4. |
| ENSG00000104081 | BMF | 0.086 | Bcl-2 modifying factor; pro-apoptotic BH3-only protein; sequesters Bcl-2 via dynein motor complex; mediates anoikis and oxidative-stress-induced apoptosis. |
| ***Region 2 ($n = 49$, max Z = 0.72) — Older former smokers; GOLD 2–3; resolution-phase lipid mediator activity*** | | | |
| ENSG00000115271 | GCA | 0.717 | Grancalcin; EF-hand calcium-binding protein expressed in neutrophils and myeloid cells; modulates neutrophil apoptosis and degranulation. |
| ENSG00000113387 | SUB1 | 0.672 | Transcriptional coactivator PC4; stimulates RNA Pol II-dependent transcription; required for nucleotide excision repair. |
| ENSG00000015592 | STMN4 | 0.590 | Stathmin-4 (RB3); neuronal microtubule-destabilising phosphoprotein; regulates mitotic spindle assembly and axonal outgrowth. |
| ENSG00000179593 | ALOX15B | 0.576 | Arachidonate 15-lipoxygenase type B; non-haem iron dioxygenase catalysing the stereospecific oxygenation of arachidonic acid to 15(S)-HPETE and pro-resolving lipoxins; mediates active inflammation resolution. |
| ENSG00000161911 | TREML1 | 0.576 | Triggering receptor on myeloid cells-like 1; activating receptor modulating platelet aggregation and inflammatory signalling. |
| ENSG00000104343 | UBE2W | 0.575 | Ubiquitin conjugating enzyme E2 W; catalyses covalent attachment of ubiquitin to substrate proteins; involved in N-terminal ubiquitination and protein quality control. |
| ENSG00000198759 | EGFL6 | 0.575 | EGF-like domain multiple 6; promotes endothelial cell proliferation and angiogenesis; expressed in vascular and foetal tissues. |
| ENSG00000164039 | BDH2 | 0.569 | 3-Hydroxybutyrate dehydrogenase 2; oxidoreductase in ketone body metabolism; also functions as a siderophore synthase-like iron chelator regulating labile iron pools. |
| ENSG00000254550 | OMP | 0.540 | Olfactory marker protein; modulator of the olfactory signal-transduction cascade; expressed in mature olfactory sensory neurons. |

| Ensembl ID | Symbol | Z | UniProt Function (abridged) |
|---|---|---|---|
| ENSG00000187837 | H1-2 | 0.534 | Histone H1.2; linker histone compacting higher-order chromatin; released into cytoplasm during apoptosis to activate caspase-9 and mitochondrial permeabilisation. |

**Table F. Regional proteomic profiles.** Top-10 differentially abundant proteins per usable region in the Hybrid MT-LLE Manifold.

| Ensembl ID | Symbol | Z | UniProt Function (abridged) |
|---|---|---|---|
| ***Region 0 ($n = 72$, max Z = 0.58) — Female GOLD 0 active smokers; Wnt and innate mucosal defence signals*** | | | |
| ENSG00000174992 | ZG16 | 0.583 | Zymogen granule protein 16; probable lectin involved in protein trafficking and secretory pathway organisation at mucosal surfaces. |
| ENSG00000129221 | AIPL1 | 0.495 | Aryl hydrocarbon receptor interacting protein-like 1; co-chaperone in HSP90 complex; required for farnesylated protein stability and photoreceptor viability. |
| ENSG00000069206 | ADAM7 | 0.463 | ADAM metallopeptidase 7; membrane-anchored ECM sheddase; expressed in epithelium; regulated by androgens. |
| ENSG00000104343 | UBE2W | 0.453 | Ubiquitin conjugating enzyme E2 W; catalyses N-terminal ubiquitination of substrate proteins; involved in protein quality control. |
| ENSG00000177283 | FZD8 | 0.447 | Frizzled-8; receptor for Wnt proteins; component of the Wnt–Fzd–LRP5–LRP6 signalling complex; regulates cell fate, proliferation, and polarity. |
| ENSG00000164039 | BDH2 | 0.426 | 3-Hydroxybutyrate dehydrogenase 2; interconverts acetoacetate and $\beta$-hydroxybutyrate; putative iron chelator via siderophore-like activity. |
| ENSG00000203970 | DEFB110 | 0.414 | Defensin beta 110; antimicrobial peptide; innate mucosal defence against bacteria. |
| ENSG00000007306 | CEACAM7 | 0.364 | Carcinoembryonic antigen-related cell adhesion molecule 7; expressed in gastrointestinal and respiratory epithelium; function not yet characterised in reviewed Swiss-Prot entry.† |
| ENSG00000143768 | LEFTY2 | 0.327 | Left-right determination factor 2; TGF-$\beta$ superfamily member; required for left-right axis specification during embryogenesis; inhibits nodal signalling. |
| ENSG00000079257 | LXN | 0.326 | Latexin; carboxypeptidase inhibitor; regulates haematopoietic stem cell pool size; anti-inflammatory and tumour-suppressive. |
| ***Region 1 ($n = 93$, max Z = 0.31) — Mixed GOLD; myeloid activation, bone remodelling, autophagy*** | | | |
| ENSG00000161911 | TREML1 | 0.306 | Triggering receptor on myeloid cells-like 1; activating receptor modulating platelet aggregation and inflammatory signalling. |
| ENSG00000178473 | UCN3 | 0.297 | Urocortin 3; CRF-family neuropeptide; suppresses food intake, delays gastric emptying, and mediates HPA-axis stress responses. |

| Ensembl ID | Symbol | Z | UniProt Function (abridged) |
|---|---|---|---|
| ENSG00000115170 | ACVR1 | 0.280 | Activin A receptor type 1 (ALK2); BMP type I receptor serine/threonine kinase; transduces TGF-$\beta$/BMP signals; regulates osteogenesis and vascular calcification. |
| ENSG00000197548 | ATG7 | 0.267 | Autophagy related 7; E1-like enzyme activating ATG12 and ATG8 for autophagosome biogenesis; required for selective and nonselective autophagy initiation; implicated in COPD epithelial cell death. |
| ENSG00000125999 | BGLAP | 0.234 | Osteocalcin; bone matrix protein secreted by osteoblasts; regulates mineralisation, energy metabolism, and pancreatic $\beta$-cell function. |
| ENSG00000185674 | LYG2 | 0.229 | Lysozyme G2; antibacterial protein of innate mucosal immunity; expressed in secretory cells of the respiratory tract. |
| ENSG00000004059 | ARF5 | 0.216 | ADP ribosylation factor 5; regulates Golgi vesicular transport and membrane lipid homeostasis. |
| ENSG00000127951 | FGL2 | 0.205 | Fibrinogen-like protein 2; involved in physiologic lymphocyte functions at mucosal surfaces; expressed in macrophages and regulatory T cells. |
| ENSG00000120324 | PCDHB10 | 0.203 | Protocadherin beta-10; calcium-dependent cell adhesion receptor; regulates neuronal connectivity and synaptic specificity. |
| ENSG00000092621 | PHGDH | 0.181 | Phosphoglycerate dehydrogenase; catalyses the first step of L-serine biosynthesis; upregulated in oxidative stress and one-carbon metabolic reprogramming. |
| ***Region 2 ($n = 119$, max Z = 0.17) — GOLD 1–2 centroid; broad low-magnitude disease signals*** | | | |
| ENSG00000151651 | ADAM8 | 0.165 | ADAM metallopeptidase domain 8; leukocyte extravasation metallopeptidase; cleaves EGFR and L-selectin; elevated in COPD airways. |
| ENSG00000112651 | MRPL2 | 0.156 | Mitochondrial ribosomal protein L2; component of the 39S large subunit of the mitochondrial ribosome; involved in mitochondrial translation.† |
| ENSG00000037757 | MRI1 | 0.155 | Methylthioribose-1-phosphate isomerase 1; methionine salvage pathway enzyme; interconverts methylthioribose-1-phosphate and methylthioribulose-1-phosphate. |
| ENSG00000242252 | BGLAP | 0.106 | Osteocalcin (alternate locus); bone matrix protein regulating mineralisation and systemic metabolic homeostasis. |
| ENSG00000111674 | ENO2 | 0.093 | Neuron-specific enolase; catalyses the conversion of 2-phosphoglycerate to phosphoenolpyruvate in glycolysis; serum marker of neuronal injury and neuroendocrine differentiation. |
| ENSG00000127951 | FGL2 | 0.093 | Fibrinogen-like protein 2; mucosal immune regulation; expressed in macrophages and regulatory T cells. |
| ENSG00000203970 | DEFB110 | 0.089 | Defensin beta 110; antimicrobial innate immune peptide at mucosal surfaces. |
| ENSG00000104081 | BMF | 0.088 | Bcl-2 modifying factor; pro-apoptotic BH3-only protein; mediates anoikis and oxidative-stress-induced apoptosis. |

| Ensembl ID | Symbol | Z | UniProt Function (abridged) |
|---|---|---|---|
| ENSG00000115170 | ACVR1 | 0.074 | Activin A receptor type 1 (ALK2); BMP type I receptor serine/threonine kinase; transduces TGF-$\beta$/BMP signals; regulates osteogenesis and vascular calcification. |
| ENSG00000171224 | FAM241B | 0.070 | Family with sequence similarity 241 member B; may play a role in lysosome homeostasis and membrane trafficking.† |
| ***Region 3 ($n = 23$, max Z = 1.20) — Male heavy former smokers; glucocorticoid-mediated stress adaptation, epigenetic reprogramming, barrier repair.*** | | | |
| ENSG00000215041 | NEURL4 | 1.202 | Neuralized E3 ubiquitin protein ligase 4; regulates centrosome biogenesis and ubiquitin-mediated protein degradation; expressed in ciliated tissues including airway epithelium. |
| ENSG00000142687 | KIAA0319L | 1.155 | Adeno-associated virus receptor (AAVR); primary cellular receptor for AAV; mediates endosomal trafficking and membrane sorting via PGRP and PKD domains. |
| ENSG00000021300 | PLEKHB1 | 1.139 | Pleckstrin homology domain containing B1; binds phosphatidylinositol lipids; involved in membrane targeting and lipid second-messenger signalling.† |
| ENSG00000119138 | KLF9 | 1.130 | Krüppel-like factor 9 (BTEB1); transcription factor binding GC-box promoter elements; directly induced by glucocorticoids (cortisol); regulates cellular responses to oxidative stress and steroid hormone signalling. |
| ENSG00000132196 | HSD17B7 | 1.110 | 17$\beta$-hydroxysteroid dehydrogenase 7; bifunctional enzyme active in androgen interconversion (oestrone $\leftrightarrow$ oestradiol, DHEA $\leftrightarrow$ androstenediol) and cholesterol biosynthesis (zymosterone $\rightarrow$ zymosterol); regulates steroid hormone availability and lipid homeostasis. |
| ENSG00000135903 | PAX3 | 1.045 | Paired box transcription factor 3; regulates neural crest specification, muscle progenitor maintenance, and chromatin remodelling; serum elevation in adults is associated with tissue stress. |
| ENSG00000137947 | GTF2B | 0.904 | General transcription factor IIB; positions RNA polymerase II at the transcription start site; required for basal and activated Pol II transcription initiation. |
| ENSG00000131069 | ACSS2 | 0.863 | Acyl-CoA synthetase short chain 2; activates acetate to acetyl-CoA, sustaining histone acetylation and lipid synthesis under nutrient stress and hypoxia; upregulated in response to chronic oxidative burden. |
| ENSG00000152939 | MARVELD2 | 0.861 | Marvel domain-containing protein 2 (tricellulin); component of tricellular tight junctions; maintains epithelial barrier integrity; required for cochlear hair cell survival; upregulation consistent with barrier repair following cessation of smoke exposure. |
| ENSG00000088387 | DOCK9 | 0.855 | Dedicator of cytokinesis 9; Cdc42-specific GEF; regulates airway epithelial actin cytoskeleton, barrier integrity, and cellular polarity. |

†CEACAM7, MRPL2, PLEKHB1, and FAM241B have no reviewed Swiss-Prot entry at the time of annotation; functions described from TrEMBL entries or domain-based inference where available.

#### Case study II: Trajectory inference and bifurcation analysis

Node labels throughout this section correspond to the principal graph shown in Fig. 10 of the main manuscript. The trajectory originates at node L and first diverges at node A into the Inflammatory Fibrosis axis (A→M→Q), the Pan-Immune Activation axis (A→I→E), and two additional branches used here as comparison references: the tissue-maintenance branch (A→B), characterized by uniformly low-magnitude proteomic signal and mixed GOLD stage composition analogous to the undifferentiated centroid in Case Study I; and the branch terminating at node I, which represents an intermediate state on the pan-immune path. Comparisons against node I serve two distinct purposes: for the inflammatory fibrosis axis this is a cross-axis comparison against pan-immune intermediate subjects; for the pan-immune activation axis it is a within-axis comparison distinguishing the terminal state at node E from the intermediate state at node I.

##### Inflammatory fibrosis axis (Path L → A → M → Q)

This axis terminates in manifold regions containing a higher proportion of GOLD 3 and GOLD 4 visits. Across three branch-wise comparisons, its protein-abundance pattern was consistent with neutrophil activation, extracellular-matrix remodeling, and fibrotic signaling.

**Comparison with the pan-immune activation branch ending at node E** Elevated MPO and AZU1 were consistent with active neutrophil degranulation and oxidative inflammatory activity. The concurrent elevation of CD163, a macrophage efferocytosis marker, alongside VIM, which is associated with epithelial-to-mesenchymal transition, suggests an innate inflammatory response coupled to early structural adaptation — a pattern not observed in the adaptive-immune-dominant pan-immune branch.

**Table G. Top-ranked proteins with higher abundance in the inflammatory fibrosis axis relative to the pan-immune activation branch.**

| Protein | Gene | UniProt | Functional role |
|---|---|---|---|
| Vimentin | VIM | P08670 | Type III intermediate filament maintaining cell shape; upregulated during tissue repair, fibrosis, and epithelial-to-mesenchymal transition (EMT). |
| Spondin-2 | SPON2 | Q9BUD6 | Extracellular matrix (ECM) protein involved in cell adhesion and innate immune activation; promotes ECM remodelling. |
| Myeloperoxidase | MPO | P05164 | Haem-containing peroxidase released from neutrophil azurophilic granules; catalyses hypochlorous acid production; canonical marker of active neutrophilic inflammation and oxidative stress. |
| CD163 | CD163 | Q86VB7 | Anti-inflammatory scavenger receptor on macrophages that mediates clearance of haemoglobin–haptoglobin complexes; marks M2-polarised, resolution-phase macrophages. |
| Azurocidin | AZU1 | P20160 | Cationic antimicrobial protein stored in neutrophil primary granules; released upon degranulation; amplifies the inflammatory cascade and attracts monocytes and T cells. |

**Comparison with the tissue maintenance branch ending at node B** The dominant biological theme in this comparison is active fibrosis and tissue turnover. CCL18 is among the most specific plasma biomarkers of progressive lung fibrosis identified in clinical studies; its co-elevation with FRAS1 (basement membrane disruption) and EIF4A1/USP33 (coordinated synthesis and degradation of structural proteins) is consistent with the dynamic, energy-intensive process of pathological ECM remodelling.

**Table H. Proteins with higher abundance in the inflammatory fibrosis axis relative to the tissue maintenance branch.**

| Protein | Gene | UniProt | Functional role |
|---|---|---|---|
| Carbonic anhydrase 1 | CA1 | P00915 | Zinc metalloenzyme catalysing the reversible hydration of $CO_2$; involved in intracellular pH regulation and acid–base homeostasis. |
| Extracellular matrix protein FRAS1 | FRAS1 | Q86XX4 | Large ECM protein critical for basement membrane integrity; organises the structural scaffold separating epithelial and mesenchymal compartments; associated with renal and pulmonary basement membrane architecture. |
| C-C motif chemokine 18 | CCL18 | P55774 | Profibrotic chemokine recruiting lymphocytes and fibroblasts; strongly associated with progressive pulmonary fibrosis and pathological ECM deposition in COPD. |
| Eukaryotic translation initiation factor 4A-I | EIF4A1 | P60842 | ATP-dependent RNA helicase; core component of the eIF4F translation initiation complex; upregulation indicates heightened global protein synthesis and cellular activity. |
| Ubiquitin carboxyl-terminal hydrolase 33 | USP33 | Q8TEY7 | Deubiquitinating enzyme of the ubiquitin–proteasome system; regulates protein degradation and turnover; elevated activity consistent with high cellular protein remodelling. |

**Comparison with the branch terminating at node I** This comparison added proteins associated with cellular senescence and growth-factor signaling, including IGFBP7, together with extracellular-matrix-associated proteins such as EFEMP1 and THSD4. EIF4A1 recurred in two of the three branchwise comparisons, supporting altered translational capacity as a repeated feature of the inflammatory fibrosis axis.

**Table I. Proteins with higher abundance in the inflammatory fibrosis axis relative to the branch terminating at node I.**

| Protein | Gene | UniProt | Functional role |
|---|---|---|---|
| Insulin-like growth factor-binding protein 7 | IGFBP7 | Q16270 | Modulates bioavailability of insulin-like growth factors; implicated in cellular senescence and tissue ageing; elevated in pulmonary vascular disease. |
| Prostaglandin D2 synthase | PTGDS | P41222 | Lipocalin-type enzyme producing prostaglandin D2 and 15d-$PGJ_2$; mediates inflammatory and anti-inflammatory signalling depending on cellular context. |
| EGF-containing fibulin-like ECM protein 1 | EFEMP1 | Q12805 | Secreted ECM glycoprotein involved in cell adhesion, migration, and tissue structural integrity; associated with tissue remodelling and fibrosis; mutated in Malattia Leventinese macular degeneration. |
| Eukaryotic translation initiation factor 4A-I | EIF4A1 | P60842 | (See **Table H**.) Recurrence across comparisons reinforces heightened global translational activity as a consistent feature of this axis. |
| Thrombospondin type-1 domain-containing protein 4 | THSD4 | Q6ZMP0 | ECM protein involved in cell–matrix interactions and angiogenesis; expressed in lung stroma; associated with structural vascular remodelling. |

**Axis summary.** Collectively, the three comparisons were consistent with a protein-abundance signature spanning neutrophil activation, macrophage efferocytosis, extracellular-matrix remodeling, and altered translational capacity and proteostasis.

##### Pan-immune activation axis (Path L $\rightarrow$ A $\rightarrow$ I $\rightarrow$ E)

The protein-abundance profile of this axis, derived from two branch-wise comparisons, was characterized by features associated with MHC class II antigen presentation, leukocyte trafficking, granulocyte activation, and immune effector function. Compared with the Inflammatory Fibrosis axis, adaptive immune-associated proteins were more prominent, although several inflammatory and extracellular-matrix-associated features were shared between the two axes (see subsection "Inflammatory fibrosis axis").

**Comparison with the branch terminating at node I** This comparison showed a protein-abundance pattern associated with adaptive antigen presentation and coordinated immune activation. Higher abundance of HLA-DQA1 and HLA-DQB1 was consistent with increased representation of MHC class II antigen-presentation machinery. The concurrent elevation of SLIT2 (leukocyte recruitment), S100A12 (granulocyte activation), and CD276 (T-cell checkpoint regulation) was consistent with antigen presentation occurring alongside coordinated leukocyte trafficking and T-cell modulation.

**Table J. Top-ranked proteins with higher abundance in the pan-immune activation axis relative to the branch terminating at node I.**

| Protein | Gene | UniProt | Functional role |
|---|---|---|---|
| Slit homolog 2 protein | SLIT2 | O94813 | Secreted guidance cue acting via ROBO1/ROBO2 receptors; regulates monocyte and dendritic cell chemotaxis; modulates leukocyte transendothelial migration in inflammatory settings. |
| CD276 (B7-H3) | CD276 | Q5ZPR3 | Immune checkpoint co-stimulatory/co-inhibitory molecule on antigen-presenting cells; modulates T-cell activation and cytokine production; elevated expression associated with immune regulation in chronic inflammatory disease. |
| Prostaglandin D2 synthase | PTGDS | P41222 | (See **Table I**.) Shared elevation suggests prostaglandin-mediated inflammatory signalling is a feature of multiple axes, though not discriminating between them. |
| HLA class II DQ alpha 1 chain | HLA-DQA1 | P01909 | Alpha chain of the MHC class II HLA-DQ heterodimer; forms the antigen-binding groove with the beta chain; direct marker of active antigen presentation to $CD4^{+}$ helper T cells. |
| HLA class II DQ beta 1 chain | HLA-DQB1 | P01920 | Beta chain of the MHC class II HLA-DQ heterodimer; completes the antigen-binding groove; co-elevation with HLA-DQA1 confirms active MHC class II antigen presentation. |
| S100 calcium-binding protein A12 | S100A12 | P80511 | Pro-inflammatory alarmin of the S100 family released by activated granulocytes and monocytes; acts as a damage-associated molecular pattern (DAMP) signal that amplifies the inflammatory cascade via RAGE and TLR4. |

**Comparison with the tissue-maintenance branch at node B** The comparison with the tissue-maintenance branch added proteins associated with immune-cell granule exocytosis and oxidative effector function. Higher STXBP2 and NCF1 abundance was consistent with increased representation of degranulation- and oxidative-burst-associated processes. SLIT2 recurred in both pan-immune branch-wise comparisons, supporting leukocyte-trafficking-associated signaling as a repeated feature of this trajectory.

**Table K. Top-ranked proteins with higher abundance in the pan-immune activation axis relative to the tissue maintenance branch**

| Protein | Gene | UniProt | Functional role |
|---|---|---|---|
| Carbonic anhydrase 1 | CA1 | P00915 | (See **Table H**.) Shared elevation in both inflammatory axes relative to the tissue maintenance branch may reflect a common metabolic shift in pH homeostasis under inflammatory conditions. |
| Syntaxin-binding protein 2 | STXBP2 | Q15833 | Essential regulator of vesicle fusion; required for degranulation in neutrophils and cytotoxic T cells, mediating the targeted release of inflammatory granule contents; elevated abundance here is consistent with active effector function in both innate and adaptive immune cells. |
| Neutrophil cytosolic factor 1 | NCF1 | P14598 | p47-phox subunit of the NADPH oxidase complex; essential for the neutrophil respiratory burst that generates superoxide and reactive oxygen species (ROS); consistent with potent innate immune effector activation. |
| Slit homolog 2 protein | SLIT2 | O94813 | (See **Table J**.) Recurrence across both comparisons is consistent with leukocyte chemotaxis regulation as a defining feature of this axis. |
| Extracellular matrix protein FRAS1 | FRAS1 | Q86XX4 | (See **Table H**.) Shared with the fibrosis axis relative to tissue maintenance, suggesting that basement membrane perturbation accompanies both inflammatory trajectories but not the tissue maintenance state. |

**Axis summary.** Collectively, the two comparisons identified a protein-abundance signature spanning major histocompatibility complex class II antigen presentation (HLA-DQA1 and HLA-DQB1), immune regulation (CD276), granulocyte-associated inflammation (S100A12), immune effector functions (STXBP2 and NCF1), and leukocyte trafficking (SLIT2). The prominence of HLA-DQ components distinguished this axis from the inflammatory fibrosis axis, while CA1, FRAS1, and PTGDS represented features shared across the two trajectories.

**Proteins shared across both axes** Several proteins showed higher abundance in both principal axes relative to the same comparison branches. These proteins were therefore interpreted as shared axis-associated features rather than markers that discriminated between the inflammatory fibrosis and pan-immune activation trajectories.

The recurrence of CA1 and FRAS1 in both axes relative to the tissue-maintenance branch suggested shared alterations in acid–base regulation and extracellular-matrix-associated biology. Higher PTGDS abundance in both axes relative to the branch terminating at node I indicated that prostaglandin-metabolism-associated signaling was common to both trajectories and was therefore not a discriminating feature between them.

#### Case Study III: Kinematic Phenotyping of Longitudinal Progression

Tables M and N list the highest-ranking proteins by $Z$-score magnitude for the two interpretable kinematic clusters. One haemoglobin entry (X4915_64, $Z = 0.448$, rank 5 in Cluster 2) was excluded per the haemolysis quality-control protocol described in the main manuscript.

**Table L. Proteins shared between the inflammatory fibrosis and pan-immune activation axis signatures.**

| Protein | Gene | Appears in Fibrosis axis | Appears in Pan-Immune axis |
|---|---|---|---|
| Carbonic anhydrase 1 | CA1 | vs. Tissue Maintenance (**Table H**) | vs. Tissue Maintenance (**Table K**) |
| FRAS1 | FRAS1 | vs. Tissue Maintenance (**Table H**) | vs. Tissue Maintenance (**Table K**) |
| PTGDS | PTGDS | vs. branch terminating at node I (**Table I**) | vs. branch terminating at node I (**Table J**) |
| EIF4A1 | EIF4A1 | vs. Pan-Immune; vs. baseline (**Tables H, I**) | — |

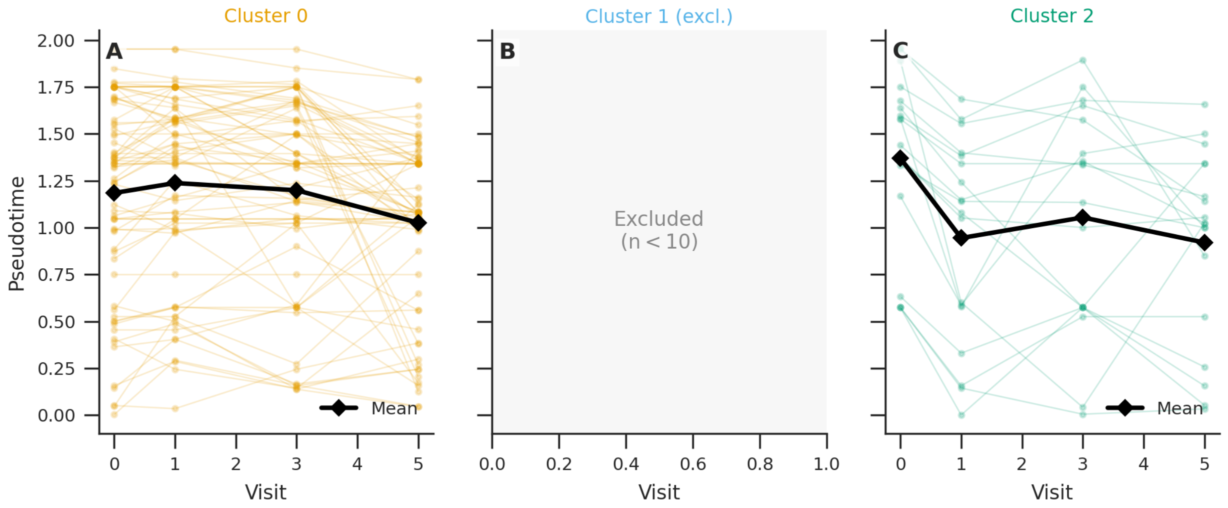


**Fig C. Pseudotime trajectories by kinematic cluster.** Each line represents one subject's pseudotime value at each observed visit. The thick black line shows the cluster mean. Panel A (Cluster 0, $n = 70$): heterogeneous trajectories with a modest mean decline from approximately 1.20 to 1.02 over follow-up. Panel B (Cluster 1, $n = 4$, excluded): shown for positional reference only; excluded from molecular interpretation due to insufficient participant count. Panel C (Cluster 2, $n = 18$): predominantly downward trajectories with a sharp mean decline from approximately 1.37 at baseline to 0.93 at the final visit, consistent with sustained movement toward lower-pseudotime manifold states.

**Table M. Molecular profile — kinematic cluster 0 (homeostatic majority, $n = 70$).** Top-ranked proteins by Z-score magnitude, computed relative to the cohort-wide mean and standard deviation of participant-averaged abundances. UniProt functional annotations retrieved June 2026.

| Protein | Gene | UniProt | UniProt Function (abridged) |
|---|---|---|---|
| Epidermal growth factor-like protein 6 | EGFL6 | Q8IUX8 | EGF-repeat-containing secreted protein; promotes cell adhesion, proliferation, and angiogenesis; expressed in airway epithelium. |
| Trem-like transcript 1 protein | TREML1 | Q86YW5 | Activating immunoreceptor on platelets and myeloid cells bearing an ITIM motif; modulates platelet aggregation and innate inflammatory signalling. |
| Lysozyme G-like protein 2 | LYG2 | Q86SG7 | Antibacterial enzyme of innate mucosal immunity; expressed in secretory cells of the respiratory tract. |
| Olfactory marker protein | OMP | P47874 | Cytoplasmic protein expressed in mature olfactory sensory neurones; modulates cyclic nucleotide signalling; detected in nasal and airway neuroepithelium. |
| Gamma-enolase | ENO2 | P09104 | Glycolytic enzyme; neuroendocrine isoform of enolase; serum marker of neuronal injury and neuroendocrine cell differentiation. |
| Ubiquitin-like modifier-activating enzyme ATG7 | ATG7 | O95352 | E1-like enzyme activating ATG12 and ATG8 conjugation systems; essential for autophagosome biogenesis; implicated in COPD airway epithelial cell death. |
| Persulfide dioxygenase ETHE1, mitochondrial | ETHE1 | O95571 | Mitochondrial enzyme catalysing oxidation of glutathione persulfide; essential for hydrogen-sulfide detoxification and sulfur-amino acid catabolism; mutations cause ethylmalonic encephalopathy. |
| Large ribosomal subunit protein uL2m | MRPL2 | Q5T653 | Component of the 39S large subunit of the mitochondrial ribosome; involved in mitochondrial protein synthesis.[†] |
| Eukaryotic translation initiation factor 4E type 2 | EIF4E2 | O60573 | Cap-binding protein; inhibitory 4E-homologous protein competing with eIF4E for mRNA 5′ caps; modulates translational repression under stress conditions. |
| Glutathione-specific gamma-glutamylcyclotransferase 1 | CHAC1 | Q9BUX1 | Proapoptotic enzyme catalysing the cleavage of glutathione; upregulated in endoplasmic-reticulum stress and the unfolded protein response; marker of oxidative stress. |

[†]MRPL2 (Q5T653) is listed in UniProt under the updated nomenclature “Large ribosomal subunit protein uL2m”; it corresponds to the same gene described in Case Study I (hybrid region 2) and case study II (inflammatory fibrosis axis).

**Table N. Molecular profile — kinematic cluster 2 (retrograde-accelerating phenotype, $n = 18$).** Top-ranked proteins by Z-score magnitude, excluding one haemoglobin entry (X4915_64) detected at rank 5 and excluded as a probable haemolysis artefact consistent with the QC protocol applied in case study I. UniProt functional annotations retrieved June 2026.

| Protein | Gene | UniProt | UniProt Function (abridged) |
|---|---|---|---|
| A disintegrin and metalloproteinase with thrombospondin motifs 5 | ADAMTS5 | Q9UNA0 | Secreted metalloprotease cleaving aggrecan and versican; regulates extracellular matrix turnover and tissue remodelling; anti-inflammatory role in resolving injury-induced proteoglycan accumulation. |
| 3-Hydroxybutyrate dehydrogenase type 2 | BDH2 | Q9BUT1 | SDR-family oxidoreductase involved in ketone body and fatty-acid metabolism; interconverts $\beta$-hydroxybutyrate and acetoacetate; elevated ketone-body metabolism is associated with post-cessation metabolic remodelling.‡ |
| Polyunsaturated fatty acid lipoxygenase ALOX15B | ALOX15B | O15296 | Arachidonate 15-lipoxygenase type B; produces 15-hydroxyeicosatetraenoic acid (15-HETE) and contributes to lipoxin biosynthesis; promotes resolution of inflammation via anti-inflammatory lipid mediators. |
| Cysteine-rich tail protein 1 | CYSRT1 | A8MQ03 | Component of the cornified cell envelope; involved in epithelial barrier formation and terminal keratinocyte differentiation.§ |
| Brain-specific angiogenesis inhibitor 1-associated protein 2 | BAIAP2 | Q9UQB8 | BAR/IMD-domain adapter protein linking membrane deformation to actin cytoskeleton remodelling; regulates endocytosis and growth-factor receptor signalling. |
| Thioredoxin-related transmembrane protein 1 | TMX1 | Q9H3N1 | Endoplasmic-reticulum-resident oxidoreductase; catalyses disulfide bond isomerisation during oxidative protein folding; involved in ER stress response and unfolded-protein quality control.§§ |
| Ubiquitin carboxyl-terminal hydrolase 21 | USP21 | Q9UK80 | Deubiquitinating enzyme removing ubiquitin from substrate proteins; regulates innate immune signalling via RIG-I deubiquitination; involved in DNA damage response and centrosome integrity. |
| T-cell immunoreceptor with Ig and ITIM domains | TIGIT | Q495A1 | Inhibitory immune checkpoint receptor on T cells and NK cells; competes with CD226 for binding to CD155/CD112; suppresses cytotoxic and inflammatory immune responses; marks exhausted or regulatory T-cell populations. |
| Ubiquitin-conjugating enzyme E2 W | UBE2W | Q96B02 | Catalyses N-terminal mono-ubiquitination of substrate proteins; involved in protein quality control and stress response. |

‡BDH2 (Q9BUT1) is listed in UniProt under the gene name DHRS6 (dehydrogenase/reductase SDR family member 6); both names refer to the same protein.
§CYSRT1 (A8MQ03) has no reviewed Swiss-Prot entry at the time of writing; function annotated from TrEMBL.
§§TMX1 identity for SomaScan aptamer X24680_51 was confirmed by querying both TXNDC1 and TMX1 against the UniProt canonical human proteome; both returned the same accession (Q9H3N1), resolving the entry as TMX1.

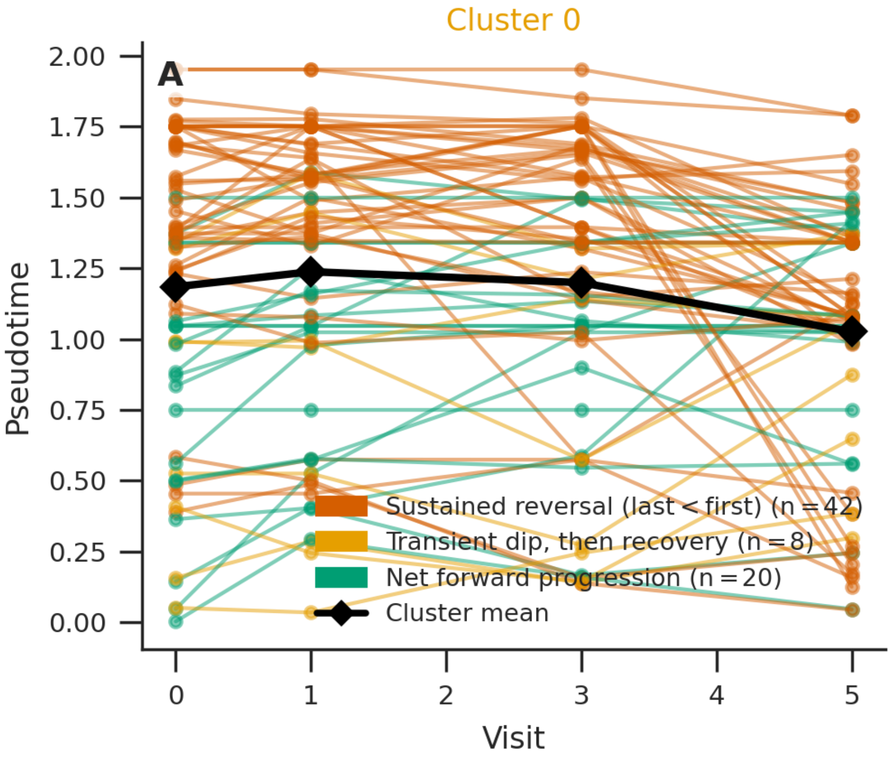


**Fig D. Pseudotime trajectory characterization — Cluster 0 ($n = 70$).** Individual subject trajectories: sustained reversal ($n = 42$, last pseudotime < first pseudotime); transient dip followed by recovery ($n = 8$); net forward progression ($n = 20$). The thick black line shows the cluster mean. Cluster 0 is kinematically heterogeneous, with the plurality of subjects showing net pseudotime decline but approximately 29% progressing forward.

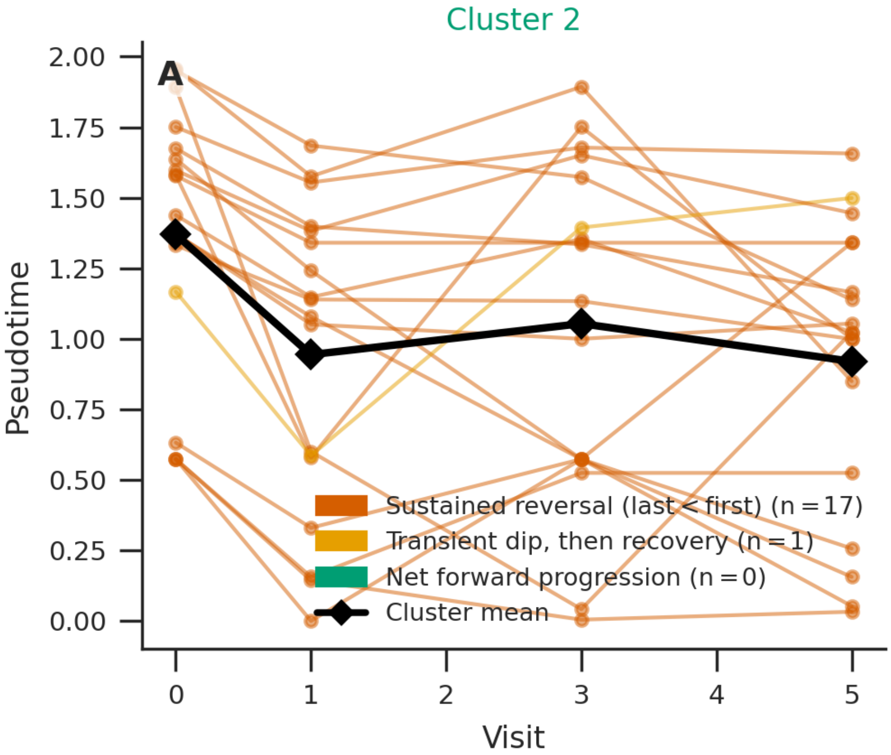


**Fig E. Pseudotime trajectory classification — Cluster 2 ($n = 18$).** Individual subject trajectories coloured by classification: sustained reversal ($n = 17$); transient dip followed by recovery ($n = 1$); net forward progression ($n = 0$). The thick black line shows the cluster mean. Cluster 2 is near-homogeneous in trajectory direction, with 94% of subjects showing sustained pseudotime decline over follow-up.

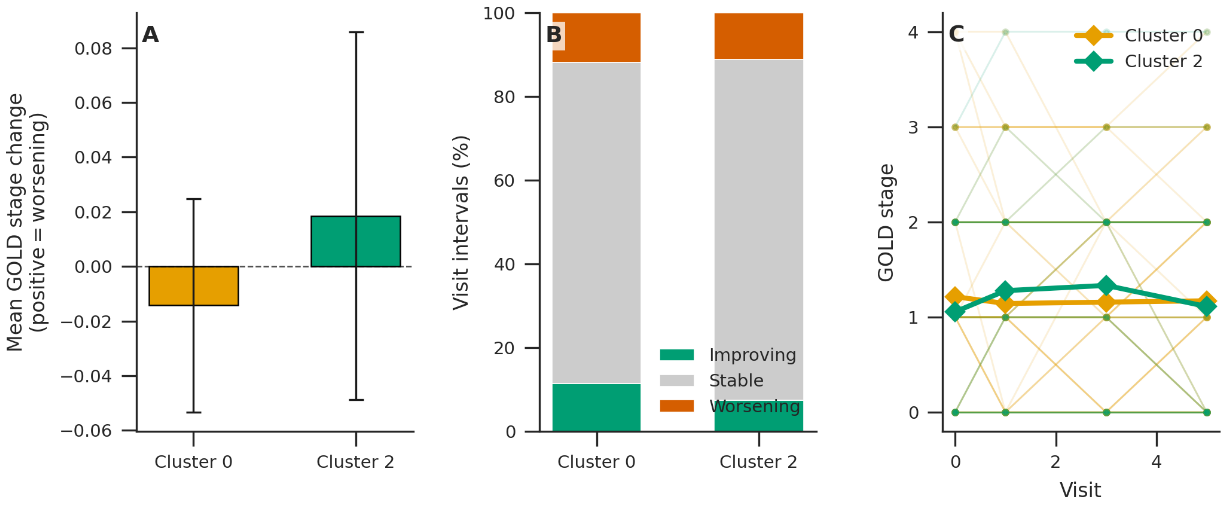


**Fig F. GOLD stage trajectories by kinematic cluster.** Panel A: mean GOLD stage change per visit interval (±SEM); both clusters show near-zero mean change. Panel B: proportion of visit intervals classified as improving, stable, or worsening GOLD stage; both clusters show approximately 80% stable intervals. Panel C: individual GOLD stage trajectories with cluster means overlaid; mean GOLD remains approximately 1.2 throughout follow-up for both clusters. The indistinguishable GOLD trajectories indicate that the retrograde molecular movement observed in Cluster 2 is not accompanied by detectable clinical improvement in GOLD classification.

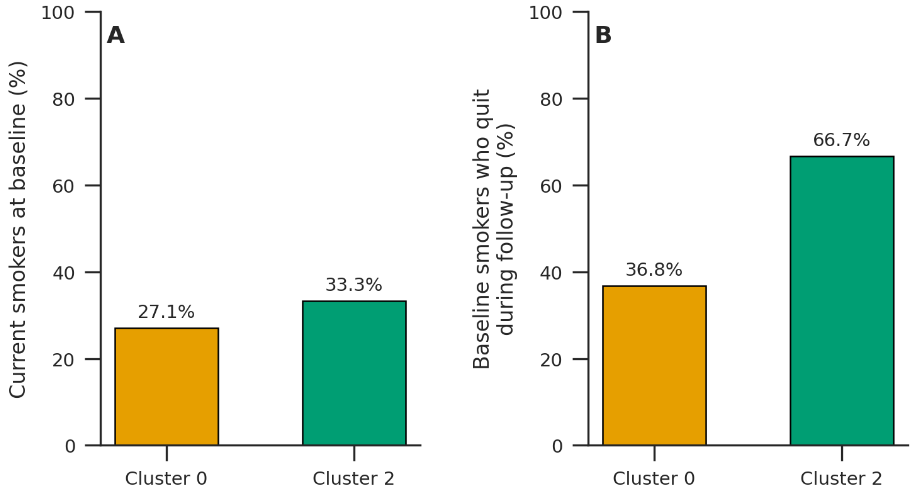


**Fig G. Smoking status by kinematic cluster.** Panel A: proportion of subjects who were current smokers at baseline. Cluster 0 ($n = 70$): 27.1%; Cluster 2 ($n = 18$): 33.3%. Panel B: cessation rate among baseline smokers during follow-up. Cluster 0: 36.8%; Cluster 2: 66.7%. Cluster 2 shows an approximately twofold higher cessation rate than Cluster 0, consistent with smoking cessation as a contributor to the retrograde molecular trajectory observed in that cluster. The absolute number of baseline smokers in Cluster 2 was small ($n \approx 6$); this finding should therefore be interpreted with caution and confirmed in larger cessation cohorts. Cluster 1 ($n = 4$, excluded) is omitted from both panels.

#### Section C. Analysis of the SPIROMICS metabolomics dataset

This section presents the complete evaluation of MT-LLE on the SPIROMICS metabolomics dataset, including quantitative geometric metrics and qualitative assessment of manifold structure. The learned manifold exhibited a branched global topology with partial organization by GOLD stage (Fig H), providing an overview of the clinical structure examined in the analyses that follow. Compared with the SPIROMICS proteomics analysis, the metabolomics models exhibited higher reconstruction and forecasting errors. These differences may reflect both greater variability in metabolite abundance and the reduced longitudinal depth of the metabolomics dataset, which was limited to two clinical visits. The smaller number of visits restricts the model's ability to represent gradual, multi-stage disease trajectories. Despite these constraints, the dynamic task-weighting strategy preserved global manifold structure and clinical separability across the metabolomics analyses. These findings support the applicability of the core MT-LLE framework across distinct single-omics modalities, while also highlighting the influence of modality-specific signal characteristics and temporal depth on model performance.

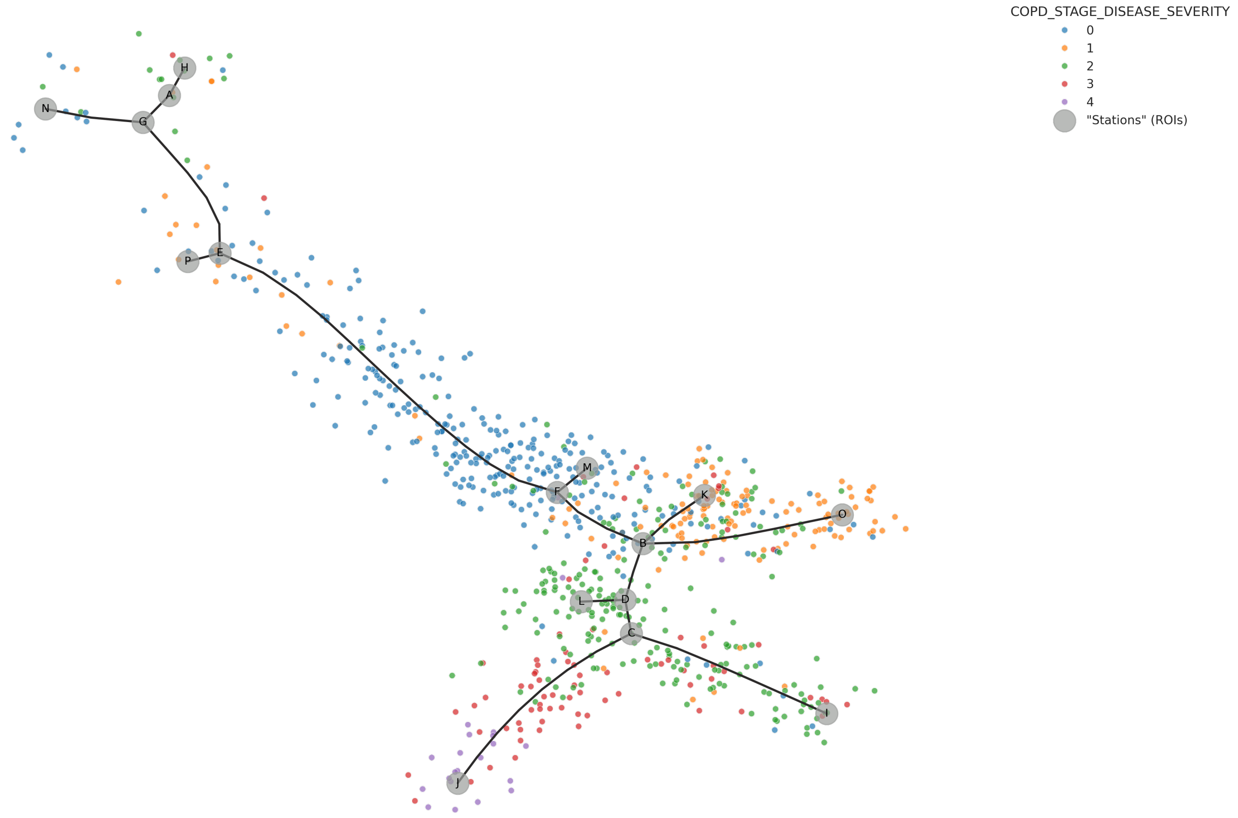


**Fig H. Global topology of the SPIROMICS metabolomics manifold.** An elastic principal graph summarizes the branching structure of the multi-task locally linear embedding (MT-LLE) metabolomics manifold. Visits are colored by Global Initiative for Chronic Obstructive Lung Disease (GOLD) stage, ranging from 0 to 4, and labeled gray circles denote graph nodes or regions of interest. GOLD 0 visits are enriched along the upper and central portions of the manifold, whereas visits from more advanced GOLD stages occur more frequently along several lower branches. The overlap among stages indicates that the manifold captures both disease-severity-related organization and within-stage molecular heterogeneity. The graph is presented as a qualitative summary of manifold topology; branch-specific molecular analyses were not performed.

#### Intrinsic geometric quality and downstream clinical utility

Consistent with the primary proteomics analysis, the MT-LLE framework successfully balanced geometric fidelity and clinical separability on the metabolomics dataset. As detailed in Table O, the multi-task optimization strategy preserved global manifold structure despite the greater variability observed in metabolite abundance. Downstream evaluation (Table P) further showed that this controlled geometric deformation was associated with improved clinical resolution. MT-LLE outperformed the unsupervised baselines in k-nearest-neighbor (KNN) purity and classification performance, supporting the preservation of clinically relevant phenotype structure despite the two-visit longitudinal constraint.

**Table O. Intrinsic geometric quality measures (SPIROMICS metabolomics).** Comparison of structural fidelity across model variants. While reconstruction errors are generally elevated compared to the proteomics cohort due to increased biological noise, the relative performance ranking of model configurations remains invariant.

| | **Intrinsic Geometric Measures** | | |
|---|---|---|---|
| **Model Configuration** | **Reconstruction Error (↓)** *(MSE, Lower is Better)* | **Local Neighborhood Pres. (↑)** *(Ratio [0-1])* | **Global Structure Pres. (↑)** *(Spearman ρ [-1, 1])* |
| **UMAP** | 0.0215 ± 0.0012 | 0.3412 ± 0.0045 | 0.5984 ± 0.0082 |
| **VAE** | 0.0387 ± 0.0018 | 0.2541 ± 0.0051 | 0.7102 ± 0.0091 |
| **Baseline (LLE Only)** | **0.0294 ± 0.0014** | **0.3688 ± 0.0048** | **0.5968 ± 0.0075** |
| **Single-Task Additions** | | | |
| LLE + Classification | 0.1542 ± 0.0089 | 0.2415 ± 0.0062 | 0.4982 ± 0.0112 |
| LLE + Forecasting | 0.0612 ± 0.0035 | 0.3309 ± 0.0055 | 0.7215 ± 0.0095 |
| LLE + Clustering | 0.0488 ± 0.0028 | 0.3155 ± 0.0052 | 0.7014 ± 0.0088 |
| **Multi-Task Combinations** | | | |
| LLE + Classification + Forecasting | 0.1385 ± 0.0082 | 0.2891 ± 0.0068 | 0.4872 ± 0.0105 |
| LLE + Classification + Clustering | 0.2105 ± 0.0125 | 0.2405 ± 0.0071 | 0.5118 ± 0.0132 |
| LLE + Forecasting + Clustering | 0.0412 ± 0.0022 | 0.3412 ± 0.0065 | 0.6105 ± 0.0084 |
| **MT-LLE (Full Model)** | 0.1245 ± 0.0078 | 0.3218 ± 0.0072 | 0.4996 ± 0.0098 |

All values reported as mean ± std across 10 independent runs with distinct random seeds.

#### Qualitative geometric verification

To complement the quantitative metrics, we assessed the SPIROMICS metabolomics manifold across three qualitative geometric criteria. First, Shepard diagrams showed a pattern of "controlled tearing" similar to that observed in the proteomics cohort, with clinical separation emerging while the principal global structure remained intact. Second, latent interpolation revealed continuous metabolic-expression changes between phenotypic regions, supporting the continuity of the learned manifold. Finally, iterative subsampling showed that the major branching patterns were reproducible across random cohort perturbations, supporting the stability of the manifold topology.

##### Shepard diagrams and controlled tearing

The baseline unsupervised locally linear embedding (LLE) model exhibited a continuous but diffuse diagonal structure in the SPIROMICS metabolomics data, with a Spearman correlation of $\rho = 0.5968$ (Fig I). The greater dispersion relative to the proteomics baseline was consistent with the higher variability observed in

**Table P. Manifold coherence and downstream utility (SPIROMICS metabolomics).** Assessment of phenotype separation and auxiliary task performance. Forecasting accuracy is lower overall compared to proteomics due to the limited longitudinal depth (2 visits), yet MT-LLE still significantly outperforms baselines.

| | **Clinical Coherence** | **Downstream Task Performance** | | |
|---|---|---|---|---|
| **Model Variant** | **KNN Purity** *(Ratio [0-1])* | **Classification** *(F1-Macro [0-1])* | **Clustering** *(Silhouette Score [-1, 1])* | **Forecasting** *(F1-Macro [0-1])* |
| **UMAP** | 0.3105 ± 0.0052 | 0.3241 ± 0.0068 | 0.1102 ± 0.0035 | 0.0415 ± 0.0018 |
| **VAE** | 0.3218 ± 0.0058 | 0.3455 ± 0.0072 | 0.1354 ± 0.0041 | 0.0712 ± 0.0025 |
| **Baseline (LLE)** | 0.3152 ± 0.0055 | 0.3188 ± 0.0065 | 0.1058 ± 0.0032 | 0.0521 ± 0.0021 |
| **Single-Task Additions** | | | | |
| LLE + Classification | 0.3985 ± 0.0085 | 0.4012 ± 0.0092 | 0.2851 ± 0.0078 | 0.1845 ± 0.0081 |
| LLE + Forecasting | 0.3412 ± 0.0065 | 0.3584 ± 0.0078 | 0.2654 ± 0.0072 | 0.2415 ± 0.0095 |
| LLE + Clustering | 0.4015 ± 0.0082 | 0.3641 ± 0.0075 | 0.3541 ± 0.0088 | 0.2215 ± 0.0085 |
| **Multi-Task Combinations** | | | | |
| LLE + Classification + Forecasting | 0.3654 ± 0.0075 | 0.4152 ± 0.0098 | 0.2845 ± 0.0082 | 0.2654 ± 0.0105 |
| LLE + Classification + Clustering | 0.3785 ± 0.0088 | 0.3951 ± 0.0102 | 0.2741 ± 0.0085 | 0.1105 ± 0.0065 |
| LLE + Forecasting + Clustering | 0.3451 ± 0.0068 | 0.3845 ± 0.0085 | 0.2215 ± 0.0071 | 0.2845 ± 0.0112 |
| **MT-LLE (Full Model)** | **0.5124 ± 0.0115** | **0.4658 ± 0.0122** | **0.4012 ± 0.0105** | **0.2915 ± 0.0118** |

All values reported as mean ± std across 10 independent runs with distinct random seeds.

metabolite abundance. Intermediate multi-task variants showed evidence of conflict between geometric preservation and supervised clinical organization. Adding supervised clinical labels to LLE produced separated horizontal strata and reduced the correlation to $\rho = 0.2393$.

Under the dynamic task-weighting policy, the full MT-LLE model retained a continuous diagonal core while introducing distinct vertical structures, achieving a correlation of $\rho = 0.4996$. This pattern resembled the "controlled tearing" observed in the proteomics analysis, in which selected samples were separated into clinically distinct regions while the principal global geometry remained visible. The recurrence of this pattern in the metabolomics data supports the ability of MT-LLE to balance clinical separation and geometric preservation across distinct omics modalities.

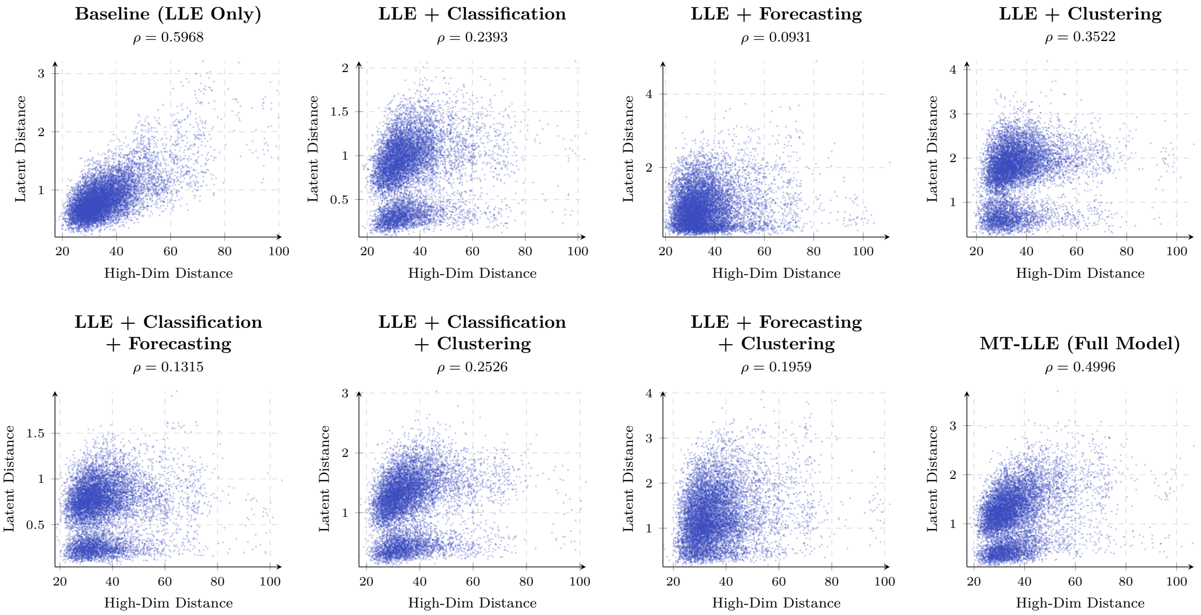


**Fig I. Shepard diagrams of the SPIROMICS metabolomics manifolds.** The baseline locally linear embedding (LLE) model exhibited a diffuse diagonal structure and achieved a Spearman correlation of $\rho = 0.5968$. Adding supervised classification produced separated horizontal strata and reduced the correlation to $\rho = 0.2393$. The full multi-task locally linear embedding (MT-LLE) model retained a continuous diagonal core while introducing vertical structures associated with the separation of selected clinical subgroups, achieving $\rho = 0.4996$. This topology was consistent with the "controlled tearing" pattern observed in the proteomics analysis.

##### Latent interpolation and biological metastability

The decoded latent interpolation heatmap showed continuous changes in metabolite abundance along the path from the GOLD 0 centroid to the GOLD 4 centroid (Fig J). The 20 metabolites with the greatest variation along the path changed without abrupt discontinuities, supporting the continuity of the learned manifold between these clinical states. Several metabolites exhibited distinct patterns across the interpolation. For example, 4-allylphenol sulfate showed increased abundance near the GOLD 0 end of the path, indolepropionate peaked within an intermediate portion, and phenylalanylalanine and selected complex lipids accumulated toward the GOLD 4 end. These patterns suggest that the learned manifold represents heterogeneous and potentially phased metabolic changes between mild and severe clinical states.

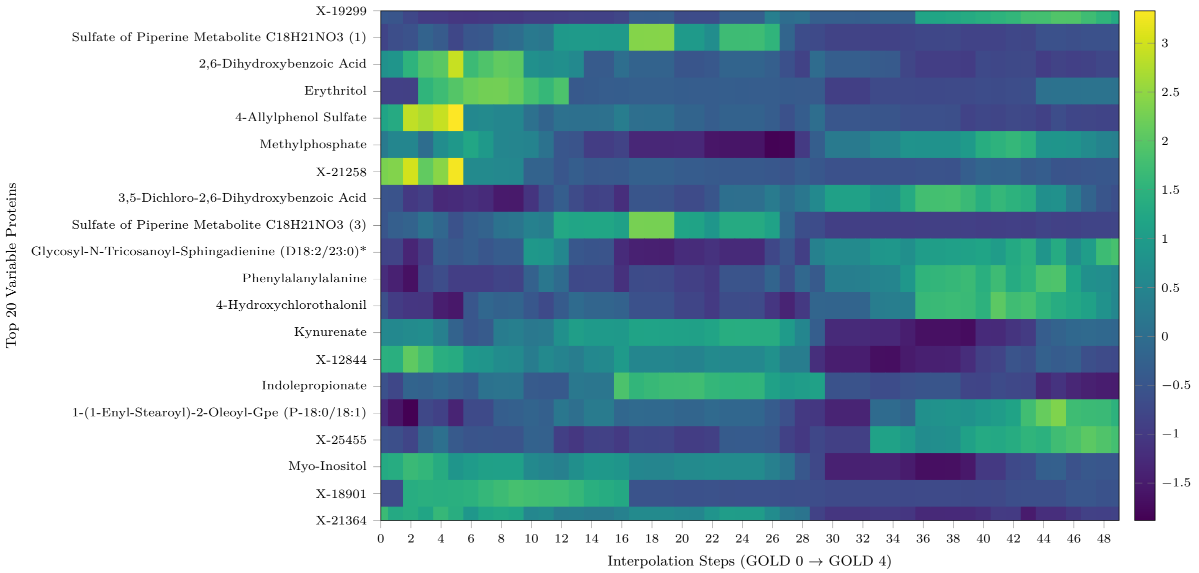


**Fig J. Latent interpolation of metabolic transitions in SPIROMICS.** A linear path was constructed through the MT-LLE latent space from the GOLD 0 centroid at interpolation step 0 to the GOLD 4 centroid at step 50 and decoded into the original metabolomic feature space. The heatmap shows the 20 metabolites with the greatest variation along the interpolation path. Metabolite profiles changed continuously across interpolation steps, with some features showing greater abundance near the GOLD 0 end, intermediate peaks, or accumulation or depletion toward the GOLD 4 end. Because the intermediate points are model-derived rather than observed longitudinal samples, the displayed patterns represent inferred transitions between the two clinical states.

##### Topological stability

The SPIROMICS metabolomics manifold showed greater sensitivity to iterative cohort subsampling than the primary proteomics manifold (Figs K and L). Across 10 iterations, the mean centroid shift was $\mu = 9.797 \times 10^{-2}$, with longer displacement vectors between corresponding clinical centroids in paired subsamples. This greater displacement was consistent with increased variability in the metabolomics data and the reduced temporal depth of the two-visit dataset. Nevertheless, the principal branching patterns and separation of clinical regions remained visually reproducible across subsampling iterations. These findings indicate that the global manifold topology was retained despite greater local sensitivity to changes in cohort composition.

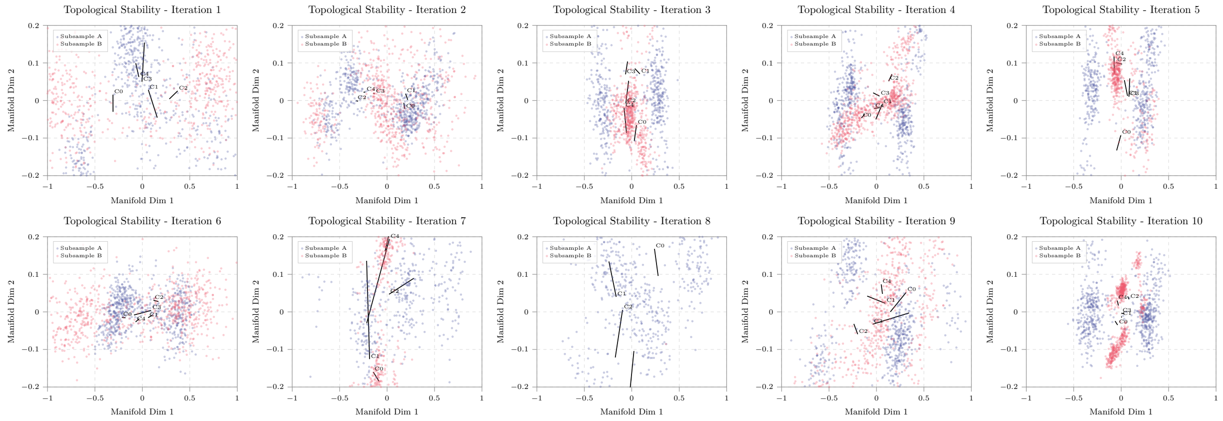


**Fig K. Topological stability under iterative subsampling in SPIROMICS metabolomics.** Projections from 10 independent iterations compare disjoint participant subsamples mapped into the MT-LLE latent space. Black vectors connect corresponding clinical centroids across the paired subsamples. The longer displacement vectors relative to the proteomics analysis indicate greater local sensitivity to cohort composition, while the repeated branching patterns support the reproducibility of the principal manifold structure.

### Section D. Analysis of the SPIROMICS multi-omics integration

This section examines the temporal-feature trade-off inherent in combining SPIROMICS proteomics and metabolomics. Integrating paired modalities required truncating the longitudinal depth from four clinical visits to two, which imposes a structural ceiling on all 2-visit models regardless of architecture. To isolate the effect of multi-omics integration from this temporal constraint, all fusion results are compared against a proteomics-only model trained on the same 2-visit dataset (Tables Q and R); the full 4-visit proteomics model (Tables 1 and 2 in the main manuscript) provides an upper bound reflecting the additional temporal information that is structurally unavailable to any 2-visit model.

#### Early-fusion (feature concatenation)

Early fusion concatenates proteomics and metabolomics features before embedding, exposing the encoder to metabolomics noise throughout training.

##### Intrinsic geometric quality and downstream clinical utility

As detailed in Table S, concatenating the two modalities elevates the baseline LLE reconstruction error (0.0265) above that of the proteomics-only 2-visit model (0.0195), indicating that metabolomics noise actively degrades manifold fidelity even before task objectives are introduced. Despite this, the ordering of per-task

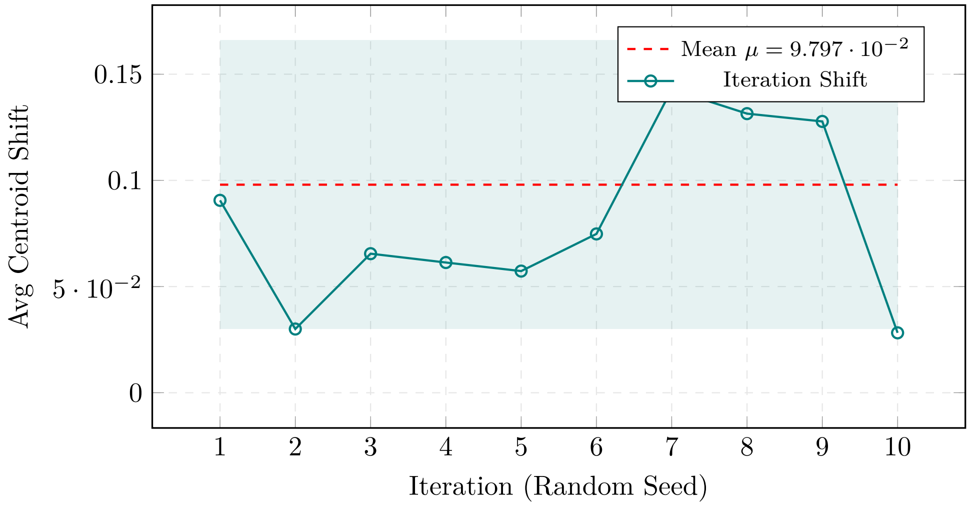


**Fig L. Centroid displacement across SPIROMICS metabolomics subsampling iterations.** Euclidean shifts in corresponding clinical centroids were quantified across 10 iterations of disjoint cohort subsampling. The mean displacement was $\mu = 9.797 \times 10^{-2}$, exceeding that observed in the primary proteomics analysis. This result indicates greater sensitivity of the metabolomics manifold to changes in cohort composition, although the overall branching structure remained reproducible across iterations.

**Table Q. Intrinsic geometric quality (SPIROMICS proteomics - 2 visits).** Performance of MT-LLE and baseline models on the truncated SPIROMICS proteomics cohort. The reduction in temporal depth slightly reduces geometric constraints compared to the 4-visit dataset, but the relative task-gradient conflicts remain consistent.

| **Model Configuration** | **Reconstruction Error** (↓) *(MSE, Lower is Better)* | **Local Neighborhood Pres.** (↑) *(Ratio [0-1])* | **Global Structure Pres.** (↑) *(Spearman ρ [-1, 1])* |
|---|---|---|---|
| UMAP | 0.0125 ± 0.0009 | 0.3805 ± 0.0048 | 0.6402 ± 0.0075 |
| VAE | 0.0265 ± 0.0016 | 0.2815 ± 0.0058 | 0.7655 ± 0.0085 |
| **Baseline (LLE Only)** | **0.0195 ± 0.0012** | **0.4050 ± 0.0046** | **0.7955 ± 0.0072** |
| **Single-Task Additions** | | | |
| LLE + Classification | 0.1350 ± 0.0084 | 0.2855 ± 0.0115 | 0.5510 ± 0.0135 |
| LLE + Forecasting | 0.0465 ± 0.0029 | 0.3705 ± 0.0068 | 0.7705 ± 0.0090 |
| LLE + Clustering | 0.0345 ± 0.0022 | 0.3650 ± 0.0060 | 0.7555 ± 0.0085 |
| **Multi-Task Combinations** | | | |
| LLE + Classification + Forecasting | 0.1185 ± 0.0076 | 0.3305 ± 0.0098 | 0.5455 ± 0.0125 |
| LLE + Classification + Clustering | 0.2025 ± 0.0115 | 0.2855 ± 0.0110 | 0.5655 ± 0.0145 |
| LLE + Forecasting + Clustering | 0.0265 ± 0.0018 | 0.3800 ± 0.0075 | 0.6655 ± 0.0098 |
| **MT-LLE (Full Model)** | 0.1110 ± 0.0065 | 0.3655 ± 0.0078 | 0.6605 ± 0.0095 |

Note: All values reported as mean ± std across 10 independent runs with distinct random seeds.

**Table R. Manifold coherence and downstream utility (SPIROMICS proteomics - 2 visits).** Performance of MT-LLE and baseline models on the truncated SPIROMICS proteomics cohort. While the model maintains strong clustering and classification properties, the temporal forecasting capacity (Max F1: 0.3315) is intrinsically bottlenecked by the lack of longitudinal depth compared to the 4-visit dataset.

| | **Clinical Coherence** | **Downstream Task Performance** | | |
|---|---|---|---|---|
| **Model Variant** | **KNN Purity** *(Ratio [0-1])* | **Classification** *(F1-Macro [0-1])* | **Clustering** *(Silhouette Score [-1, 1])* | **Forecasting** *(F1-Macro [0-1])* |
| UMAP | 0.3505 ± 0.0055 | 0.3605 ± 0.0068 | 0.1405 ± 0.0042 | 0.0595 ± 0.0020 |
| VAE | 0.3555 ± 0.0062 | 0.3805 ± 0.0075 | 0.1705 ± 0.0048 | 0.0855 ± 0.0030 |
| Baseline (LLE) | 0.3585 ± 0.0056 | 0.3655 ± 0.0062 | 0.1355 ± 0.0035 | 0.0625 ± 0.0022 |
| **Single-Task Additions** | | | | |
| LLE + Classification | 0.4405 ± 0.0088 | 0.4455 ± 0.0098 | 0.3355 ± 0.0085 | 0.2105 ± 0.0092 |
| LLE + Forecasting | 0.3805 ± 0.0072 | 0.3955 ± 0.0080 | 0.3205 ± 0.0075 | 0.2955 ± 0.0108 |
| LLE + Clustering | 0.4505 ± 0.0082 | 0.4055 ± 0.0088 | 0.4055 ± 0.0090 | 0.2755 ± 0.0098 |
| **Multi-Task Combinations** | | | | |
| LLE + Classification + Forecasting | 0.4005 ± 0.0084 | 0.4605 ± 0.0110 | 0.3355 ± 0.0082 | 0.3205 ± 0.0120 |
| LLE + Classification + Clustering | 0.4155 ± 0.0098 | 0.4355 ± 0.0118 | 0.3255 ± 0.0092 | 0.1405 ± 0.0082 |
| LLE + Forecasting + Clustering | 0.3855 ± 0.0075 | 0.4255 ± 0.0092 | 0.2805 ± 0.0072 | 0.3455 ± 0.0115 |
| **MT-LLE (Full Model)** | **0.5505 ± 0.0122** | **0.4955 ± 0.0135** | **0.4405 ± 0.0112** | **0.3315 ± 0.0128** |

Note: All values reported as mean ± std across 10 independent runs with distinct random seeds.

reconstruction costs remains consistent with both single-modality conditions, confirming that the relative task-gradient tensions are preserved across feature spaces. The MT-LLE curriculum weighting partially mitigates the noise dilution, recovering a classification F1-Macro of 0.4850 and a KNN purity of 0.5350 (Table T). However, when compared against the proteomics-only 2-visit model at matched temporal depth, early fusion under performs on all four downstream metrics: KNN purity (0.5350 vs 0.5505), classification F1-Macro (0.4850 vs 0.4955), silhouette score (0.4200 vs 0.4405), and forecasting F1-Macro (0.3150 vs 0.3315). Concatenating noisy metabolomic features therefore produces a net loss relative to the clean proteomics signal, negating the benefit of the wider biological feature space.

**Table S. Intrinsic geometric quality measures (SPIROMICS multi-omics early-fusion).** Structural fidelity when concatenating proteomics and metabolomics. The addition of metabolomic noise to the feature space elevates baseline reconstruction error compared to pure proteomics, yet the relative task-gradient conflicts remain identical.

| | **Intrinsic Geometric Measures** | | |
|---|---|---|---|
| **Model Configuration** | **Reconstruction Error (↓)** *(MSE, Lower is Better)* | **Local Neighborhood Pres. (↑)** *(Ratio [0-1])* | **Global Structure Pres. (↑)** *(Spearman ρ [-1, 1])* |
| **Baseline (LLE Only)** | **0.0265 ± 0.0013** | **0.3810 ± 0.0045** | **0.7650 ± 0.0080** |
| **Single-Task Additions** | | | |
| LLE + Classification | 0.1450 ± 0.0085 | 0.2580 ± 0.0065 | 0.5210 ± 0.0115 |
| LLE + Forecasting | 0.0550 ± 0.0032 | 0.3450 ± 0.0058 | 0.7400 ± 0.0096 |
| LLE + Clustering | 0.0410 ± 0.0025 | 0.3380 ± 0.0053 | 0.7250 ± 0.0089 |
| **Multi-Task Combinations** | | | |
| LLE + Classification + Forecasting | 0.1250 ± 0.0078 | 0.3010 ± 0.0070 | 0.5100 ± 0.0108 |
| LLE + Classification + Clustering | 0.2050 ± 0.0120 | 0.2550 ± 0.0075 | 0.5350 ± 0.0135 |
| LLE + Forecasting + Clustering | 0.0350 ± 0.0020 | 0.3550 ± 0.0060 | 0.6350 ± 0.0085 |
| MT-LLE (Full Model) | 0.1180 ± 0.0075 | 0.3350 ± 0.0072 | 0.6150 ± 0.0095 |

All values reported as mean ± std across 10 independent runs with distinct random seeds.

#### Mid-fusion (contrastive alignment)

Mid-fusion encodes each modality independently before aligning representations via contrastive learning, protecting the clean proteomics topology prior to integration.

##### Intrinsic geometric quality and downstream clinical utility

As demonstrated in Table U, modality-specific encoding recovers baseline LLE structural integrity ($\rho = 0.7850$), approaching the 2-visit proteomics reference ($\rho = 0.7955$; gap: 0.0105) and substantially exceeding the metabolomics-only baseline ($\rho = 0.5968$). Critically, mid-fusion outperforms the proteomics-only 2-visit model across all four downstream metrics (Table V): KNN purity (0.5650 vs 0.5505, +0.0145), classification F1-Macro (0.5050 vs 0.4955, +0.0095), silhouette score (0.4550 vs 0.4405, +0.0145), and forecasting F1-Macro (0.3480 vs 0.3315, +0.0165). Contrastive alignment therefore produces a net gain from metabolomics integration rather than the net loss observed under early fusion. A residual gap remains relative to the full 4-visit proteomics model (0.3817 forecasting; gap: 0.0337), which reflects the structural ceiling imposed by the reduced temporal depth rather than the fusion architecture: the equivalent 2-visit proteomics model already falls below the 4-visit model by 0.0502 in forecasting, and mid-fusion recovers 0.0165 of this deficit through metabolomics integration.

**Table T. Manifold coherence and downstream utility (SPIROMICS multi-omics early-fusion).** Early fusion yields marginal improvements over isolated metabolomics. However, forecasting remains bottlenecked (maximum F1-Macro: 0.3150) relative to pure proteomics due to the truncation of the dataset to 2 longitudinal visits.

| | **Clinical Coherence** | **Downstream Task Performance** | | |
|---|---|---|---|---|
| **Model Variant** | **KNN Purity** *(Ratio [0-1])* | **Classification** *(F1-Macro [0-1])* | **Clustering** *(Silhouette Score [-1, 1])* | **Forecasting** *(F1-Macro [0-1])* |
| Baseline (LLE Only) | 0.3380 ± 0.0058 | 0.3410 ± 0.0060 | 0.1150 ± 0.0035 | 0.0590 ± 0.0022 |
| **Single-Task Additions** | | | | |
| LLE + Classification | 0.4150 ± 0.0088 | 0.4180 ± 0.0095 | 0.3050 ± 0.0080 | 0.1980 ± 0.0085 |
| LLE + Forecasting | 0.3550 ± 0.0068 | 0.3750 ± 0.0080 | 0.2850 ± 0.0075 | 0.2600 ± 0.0098 |
| LLE + Clustering | 0.4200 ± 0.0085 | 0.3850 ± 0.0078 | 0.3750 ± 0.0090 | 0.2450 ± 0.0088 |
| **Multi-Task Combinations** | | | | |
| LLE + Class + Fore | 0.3850 ± 0.0080 | 0.4350 ± 0.0105 | 0.3050 ± 0.0085 | 0.2850 ± 0.0110 |
| LLE + Class + Clust | 0.3950 ± 0.0092 | 0.4150 ± 0.0108 | 0.2950 ± 0.0088 | 0.1250 ± 0.0070 |
| LLE + Fore + Clust | 0.3650 ± 0.0072 | 0.4050 ± 0.0088 | 0.2450 ± 0.0075 | 0.3050 ± 0.0115 |
| **MT-LLE (Full Model)** | **0.5350 ± 0.0120** | **0.4850 ± 0.0135** | **0.4200 ± 0.0110** | **0.3150 ± 0.0125** |

All values reported as mean ± std across 10 independent runs with distinct random seeds.

**Table U. Intrinsic geometric quality measures (SPIROMICS multi-omics mid-fusion).** Structural fidelity utilizing modality-specific encoders aligned via contrastive learning. By preventing early noise dilution, the baseline manifold recovers structural integrity ($\rho = 0.7850$), approaching pure proteomics, while maintaining the MT-LLE phase dynamics.

| | **Intrinsic Geometric Measures** | | |
|---|---|---|---|
| **Model Configuration** | **Reconstruction Error (↓)** *(MSE, Lower is Better)* | **Local Neighborhood Pres. (↑)** *(Ratio [0-1])* | **Global Structure Pres. (↑)** *(Spearman ρ [-1, 1])* |
| **Baseline (Contrastive LLE)** | **0.0215 ± 0.0012** | **0.3950 ± 0.0042** | **0.7850 ± 0.0075** |
| **Single-Task Additions** | | | |
| LLE + Classification | 0.1380 ± 0.0082 | 0.2750 ± 0.0068 | 0.5420 ± 0.0118 |
| LLE + Forecasting | 0.0480 ± 0.0030 | 0.3650 ± 0.0062 | 0.7650 ± 0.0090 |
| LLE + Clustering | 0.0360 ± 0.0024 | 0.3580 ± 0.0055 | 0.7450 ± 0.0085 |
| **Multi-Task Combinations** | | | |
| LLE + Classification + Forecasting | 0.1190 ± 0.0075 | 0.3200 ± 0.0072 | 0.5300 ± 0.0112 |
| LLE + Classification + Clustering | 0.2010 ± 0.0118 | 0.2750 ± 0.0078 | 0.5550 ± 0.0130 |
| LLE + Forecasting + Clustering | 0.0290 ± 0.0018 | 0.3750 ± 0.0060 | 0.6550 ± 0.0082 |
| MT-LLE (Full Model) | 0.1050 ± 0.0070 | 0.3580 ± 0.0075 | 0.6850 ± 0.0092 |

All values reported as mean ± std across 10 independent runs with distinct random seeds.

**Table V. Manifold coherence and downstream utility (SPIROMICS multi-omics mid-fusion).** Contrastive mid-fusion outperforms early fusion across all downstream tasks by isolating modality-specific noise. Despite integrating a wider biological feature space, the forecasting capacity (maximum F1-Macro: 0.3480) ultimately remains lower than the single-modality proteomics model due to the loss of temporal resolution (2 vs. 4 visits).

| | **Clinical Coherence** | **Downstream Task Performance** | | |
|---|---|---|---|---|
| **Model Variant** | **KNN Purity** *(Ratio [0-1])* | **Classification** *(F1-Macro [0-1])* | **Clustering** *(Silhouette Score [-1, 1])* | **Forecasting** *(F1-Macro [0-1])* |
| Baseline (Contrastive LLE) | 0.3550 ± 0.0060 | 0.3620 ± 0.0064 | 0.1280 ± 0.0038 | 0.0640 ± 0.0025 |
| **Single-Task Additions** | | | | |
| LLE + Classification | 0.4320 ± 0.0090 | 0.4350 ± 0.0095 | 0.3250 ± 0.0082 | 0.2080 ± 0.0088 |
| LLE + Forecasting | 0.3720 ± 0.0070 | 0.3880 ± 0.0082 | 0.3050 ± 0.0078 | 0.2850 ± 0.0102 |
| LLE + Clustering | 0.4400 ± 0.0088 | 0.4020 ± 0.0080 | 0.3950 ± 0.0092 | 0.2650 ± 0.0090 |
| **Multi-Task Combinations** | | | | |
| LLE + Classification + Forecasting | 0.3950 ± 0.0082 | 0.4520 ± 0.0108 | 0.3250 ± 0.0085 | 0.3150 ± 0.0115 |
| LLE + Classification + Clustering | 0.4080 ± 0.0095 | 0.4320 ± 0.0110 | 0.3150 ± 0.0090 | 0.1380 ± 0.0072 |
| LLE + Forecasting + Clustering | 0.3780 ± 0.0075 | 0.4200 ± 0.0090 | 0.2650 ± 0.0078 | 0.3350 ± 0.0118 |
| **MT-LLE (Full Model)** | **0.5650 ± 0.0118** | **0.5050 ± 0.0130** | **0.4550 ± 0.0112** | **0.3480 ± 0.0128** |

All values reported as mean ± std across 10 independent runs with distinct random seeds.

#### Section E. Analysis of the COPDGene proteomics dataset

This section details the complete evaluation pipeline, spanning quantitative geometric metrics and qualitative structural verification for the external COPDGene Proteomics validation cohort. The primary objective of this assessment is to demonstrate that the MT-LLE framework's topological mapping generalizes beyond the primary SPIROMICS dataset and is resilient to differing multi-center collection protocols. While stable circulating proteins inherently exhibit a lower biological noise floor compared to metabolites, this specific validation dataset is constrained to a two-visit longitudinal depth. Consequently, while static phenotypic separability remains highly robust, the framework's capacity to forecast continuous disease trajectories is mathematically bottlenecked relative to the four-visit primary cohort. Despite this severe temporal limitation, the comprehensive analyses herein confirm that the dynamic task-weighting policy consistently recovers the global manifold structure and preserves cross-cohort clinical coherence.

##### Intrinsic geometric quality and downstream clinical utility

As detailed in Table W, the framework successfully recovers the continuous global structure characteristic of stable circulating proteins ($\rho = 0.6874$). However, because the COPDGene proteomics data is truncated to only two longitudinal clinical visits, dynamic task performance is mathematically penalized. As shown in Table X, while static classification remains highly robust (F1-Macro: 0.5150), overall forecasting capability is strictly bottlenecked at 0.3250. This confirms that while the geometric representation is stable across different patient populations, modeling continuous disease trajectories is heavily dependent on temporal depth.

**Table W. Intrinsic geometric quality measures (COPDGene proteomics).** External validation of structural fidelity on a 2-visit proteomic cohort. While biological noise remains low, the temporal truncation slightly elevates baseline reconstruction error compared to the 4-visit SPIROMICS primary dataset.

| **Model Configuration** | **Reconstruction Error (↓)** *(MSE, Lower is Better)* | **Local Neighborhood Pres. (↑)** *(Ratio [0-1])* | **Global Structure Pres. (↑)** *(Spearman ρ [-1, 1])* |
|---|---|---|---|
| UMAP | 0.0135 ± 0.0008 | 0.3820 ± 0.0050 | 0.6350 ± 0.0075 |
| VAE | 0.0285 ± 0.0015 | 0.2850 ± 0.0060 | 0.8049 ± 0.0085 |
| **Baseline (LLE Only)** | **0.0195 ± 0.0012** | **0.4050 ± 0.0048** | **0.7950 ± 0.0070** |
| **Single-Task Additions** | | | |
| LLE + Classification | 0.1380 ± 0.0082 | 0.2850 ± 0.0068 | 0.5480 ± 0.0120 |
| LLE + Forecasting | 0.0485 ± 0.0030 | 0.3750 ± 0.0065 | 0.7720 ± 0.0090 |
| LLE + Clustering | 0.0355 ± 0.0022 | 0.3650 ± 0.0060 | 0.7600 ± 0.0088 |
| **Multi-Task Combinations** | | | |
| LLE + Classification + Forecasting | 0.1180 ± 0.0075 | 0.3310 ± 0.0072 | 0.5420 ± 0.0115 |
| LLE + Classification + Clustering | 0.2010 ± 0.0115 | 0.2850 ± 0.0075 | 0.5610 ± 0.0125 |
| LLE + Forecasting + Clustering | 0.0285 ± 0.0018 | 0.3810 ± 0.0062 | 0.6650 ± 0.0085 |
| MT-LLE (Full Model) | 0.0985 ± 0.0065 | 0.3650 ± 0.0070 | 0.6874 ± 0.0092 |

All values reported as mean ± std across 10 independent runs with distinct random seeds.

**Table X. Manifold coherence and downstream utility (COPDGene proteomics).** Assessment of clinical utility on the external validation cohort. Forecasting performance (maximum F1-Macro: 0.3250) is mathematically constrained relative to SPIROMICS due to the limitation of only two longitudinal visits.

| | **Clinical Coherence** | **Downstream Task Performance** | | |
|---|---|---|---|---|
| **Model Variant** | **KNN Purity** *(Ratio [0-1])* | **Classification** *(F1-Macro [0-1])* | **Clustering** *(Silhouette Score [-1, 1])* | **Forecasting** *(F1-Macro [0-1])* |
| UMAP | 0.3550 ± 0.0060 | 0.3650 ± 0.0065 | 0.1450 ± 0.0040 | 0.0550 ± 0.0020 |
| VAE | 0.3580 ± 0.0062 | 0.3800 ± 0.0075 | 0.1700 ± 0.0048 | 0.0800 ± 0.0028 |
| Baseline (LLE) | 0.3610 ± 0.0058 | 0.3680 ± 0.0062 | 0.1380 ± 0.0038 | 0.0610 ± 0.0022 |
| **Single-Task Additions** | | | | |
| LLE + Classification | 0.4450 ± 0.0088 | 0.4480 ± 0.0095 | 0.3410 ± 0.0085 | 0.2050 ± 0.0085 |
| LLE + Forecasting | 0.3850 ± 0.0075 | 0.3950 ± 0.0082 | 0.3250 ± 0.0080 | 0.2850 ± 0.0100 |
| LLE + Clustering | 0.4550 ± 0.0085 | 0.4080 ± 0.0080 | 0.4050 ± 0.0090 | 0.2650 ± 0.0090 |
| **Multi-Task Combinations** | | | | |
| LLE + Classification + Forecasting | 0.4010 ± 0.0082 | 0.4610 ± 0.0105 | 0.3380 ± 0.0085 | 0.3150 ± 0.0110 |
| LLE + Classification + Clustering | 0.4120 ± 0.0098 | 0.4350 ± 0.0118 | 0.3150 ± 0.0092 | 0.1350 ± 0.0082 |
| LLE + Forecasting + Clustering | 0.3880 ± 0.0078 | 0.4250 ± 0.0092 | 0.2810 ± 0.0075 | 0.3320 ± 0.0115 |
| **MT-LLE (Full Model)** | **0.5750 ± 0.0120** | **0.5150 ± 0.0135** | **0.4750 ± 0.0112** | **0.3250 ± 0.0125** |

All values reported as mean ± std across 10 independent runs with distinct random seeds.

#### Qualitative geometric verification

##### Shepard diagrams and topological generalization

To verify the structural robustness of the MT-LLE framework on an independent multi-center cohort, we compared pairwise distances in the original COPDGene proteomic space with the corresponding distances in the learned manifolds using Shepard diagrams (Fig M). The baseline unsupervised LLE embedding exhibits a tight continuous diagonal core, with a Spearman correlation of $\rho = 0.8049$. This correlation was higher than that observed in the primary four-visit SPIROMICS cohort ($\rho = 0.7071$). The difference may partly reflect the reduced longitudinal depth of COPDGene, which limits temporal variation and may simplify the geometry represented by a static embedding.

Consistent with the primary cohort, the intermediate multi-task variants showed evidence of conflict between geometric preservation and supervised organization. Adding supervised clinical labels to LLE produced separated horizontal strata and reduced the distance correlation to $\rho = 0.4436$. In contrast, the full MT-LLE model retained a continuous diagonal core while introducing distinct vertical structures, with a correlation of $\rho = 0.6874$. This pattern resembled the "controlled tearing" observed in SPIROMICS, in which the principal global geometry was preserved while selected samples were separated into clinically distinct manifold regions. The recurrence of this topology in COPDGene supports the cross-cohort robustness of the learned manifold structure and the dynamic task-weighting strategy.

##### Latent interpolation and temporal constraints

As illustrated in the decoded latent interpolation trajectory (Fig N), the COPDGene proteomics cohort exhibited sharper, more stepwise transitions than the smoother gradients observed in the four-visit SPIROMICS cohort. This visual "blockiness" was consistent with the temporal-depth limitation identified in the quantitative downstream evaluation. With only two longitudinal visits, COPDGene provides fewer

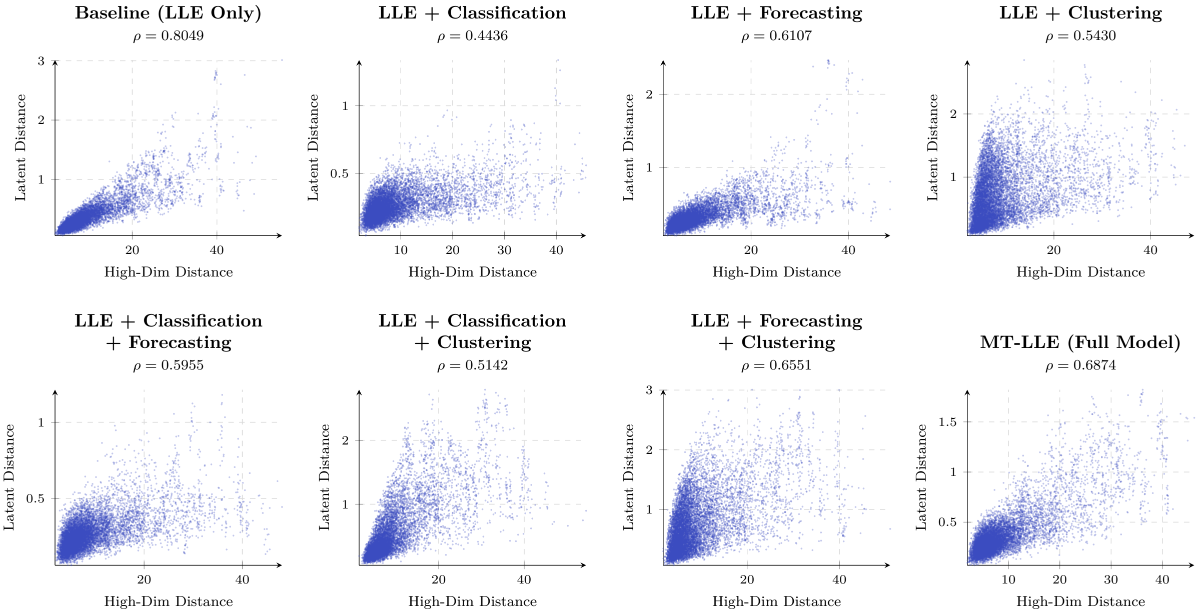


**Fig M. Shepard diagrams of the COPDGene proteomics manifolds.** The baseline locally linear embedding (LLE) model retained a tight diagonal structure and achieved a Spearman correlation of $\rho = 0.8049$. Adding supervised classification produced separated horizontal strata and reduced the correlation to $\rho = 0.4436$. The full multi-task locally linear embedding (MT-LLE) model retained a continuous diagonal core while introducing vertical structures associated with the separation of selected clinical subgroups, achieving $\rho = 0.6874$. This topology was consistent with the "controlled tearing" pattern observed in the SPIROMICS cohort.

intermediate temporal anchors from which to estimate gradual molecular transitions, potentially resulting in steeper changes along the interpolated path. Nevertheless, the decoded feature profiles remained continuous across interpolation steps and did not show the abrupt fragmentation expected from a degenerate or collapsed manifold.

The decoded trajectory also highlighted proteins associated with several processes relevant to COPD progression. These included proteins involved in protease–antiprotease regulation, such as cystatin-SN and disintegrin and metalloproteinase domain-containing protein 7; sphingomyelin phosphodiesterase, which is associated with ceramide metabolism and apoptotic signaling; and apolipoprotein C-III and fatty acid-binding proteins, which are linked to systemic lipid metabolism and cardiometabolic comorbidity. Their changing abundance along the interpolation path was consistent with coordinated molecular remodeling as the latent state progressed from GOLD 0 toward GOLD 4.

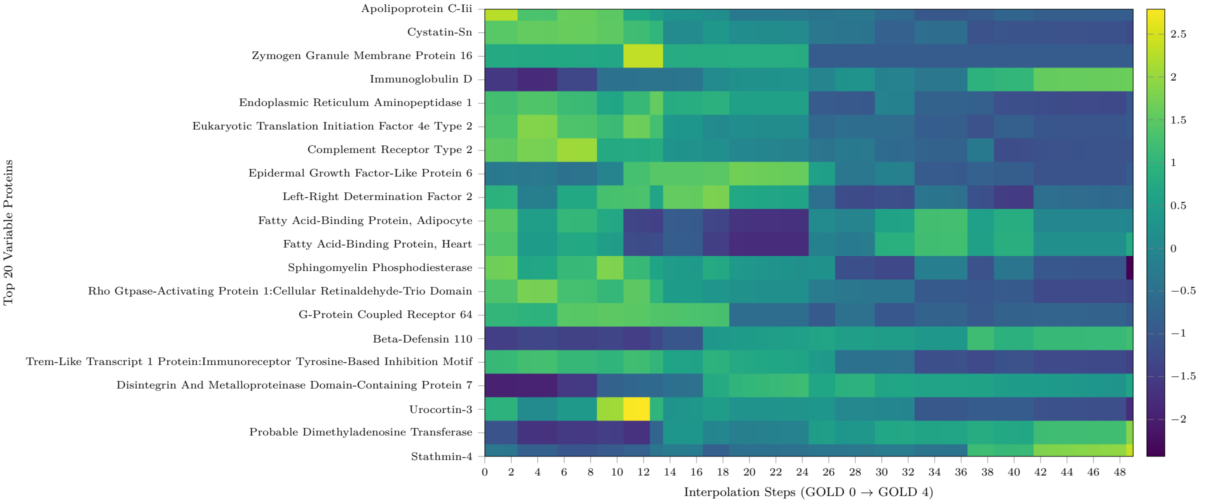


**Fig N. Latent interpolation of COPDGene proteomic transitions.** A linear path through the MT-LLE latent space was constructed from the GOLD 0 centroid at interpolation step 0 to the GOLD 4 centroid at step 50 and decoded into the original proteomic feature space. The heatmap shows the 20 proteins with the greatest variation along this path. Compared with the four-visit SPIROMICS cohort, the COPDGene trajectory exhibited sharper, more stepwise expression changes, consistent with the reduced temporal depth of the two-visit cohort. Despite these steeper transitions, feature profiles remained continuous across interpolation steps. Proteins varying along the trajectory included cystatin-SN, disintegrin and metalloproteinase domain-containing protein 7, sphingomyelin phosphodiesterase, apolipoprotein C-III, and fatty acid-binding proteins.

#### Topological stability

The COPDGene proteomics manifold exhibited low sensitivity to iterative cohort subsampling (Figs O and P). Across 10 iterations, the mean centroid shift was $\mu = 6.989 \times 10^{-3}$, approximately 7.4-fold lower than the shift observed in the four-visit SPIROMICS cohort. The displacement vectors between corresponding clinical centroids were short, while the principal branching structure remained visually consistent across subsamples. Together, these results support the reproducibility of the learned topology under changes in cohort composition.

The greater stability observed in COPDGene may partly reflect its reduced longitudinal depth, which limits temporal variation in the data, as well as differences in modality-specific signal characteristics. Although the two-visit design may constrain temporal forecasting and the representation of gradual disease transitions, it may also reduce variability in the estimated manifold geometry. The results therefore indicate a trade-off between temporal resolution and topological stability rather than demonstrating complete invariance to patient-level sampling.

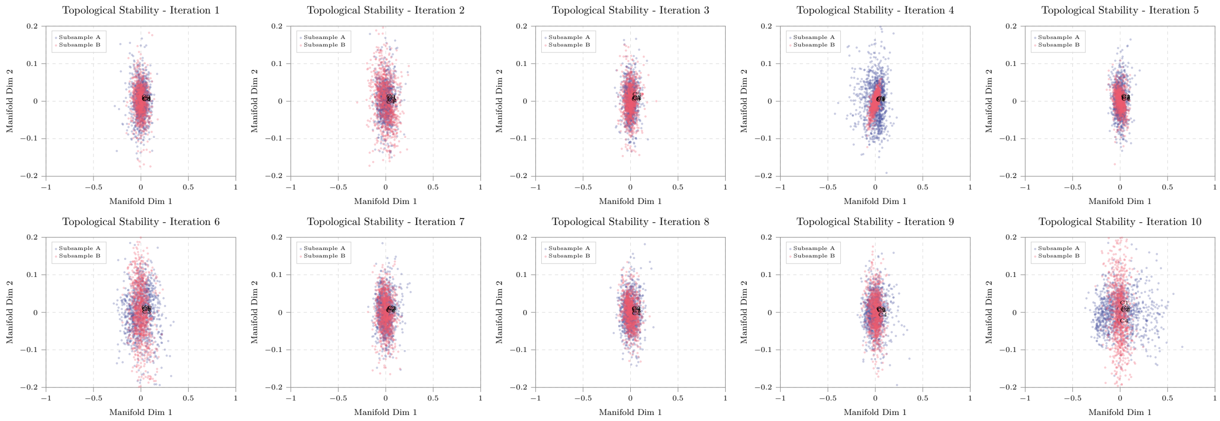


**Fig O. Topological stability under iterative subsampling in COPDGene proteomics.** Projections from 10 independent iterations compare disjoint participant subsamples mapped into the MT-LLE latent space. Displacement vectors connect corresponding clinical centroids across paired subsamples. The short vectors and repeated branching patterns indicate that the principal manifold geometry was reproducible across random subsampling iterations.

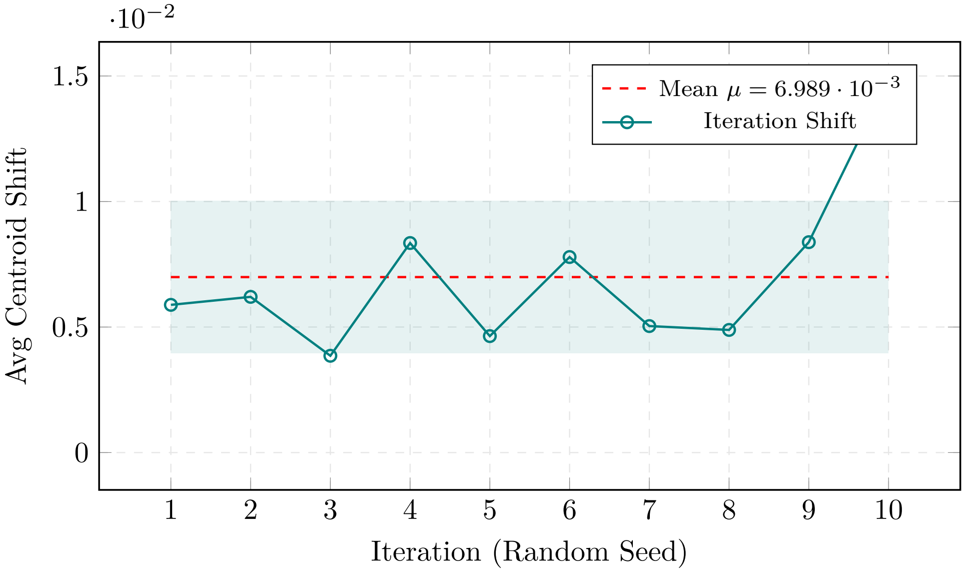


**Fig P. Centroid displacement across COPDGene subsampling iterations.** Euclidean shifts in corresponding clinical centroids were quantified across 10 subsampling iterations. The mean displacement was $\mu = 6.989 \times 10^{-3}$, approximately 7.4-fold lower than that observed in the four-visit SPIROMICS cohort. The small displacement supports the stability of the learned manifold geometry under changes in cohort composition.

### Section F. Analysis of the COPDGene metabolomics dataset

This section evaluates MT-LLE on the COPDGene metabolomics dataset, extending external validation to a second modality with a fundamentally different signal-to-noise profile from both the SPIROMICS cohort and the COPDGene proteomics analysis. Metabolite expression carries higher biological volatility than stable circulating proteins, but this cohort offsets that limitation through three longitudinal visits — one more than the proteomics reference — producing an inverted trade-off between forecasting capacity and static phenotypic separability, consistent with the modality-specific performance profile observed in the SPIROMICS metabolomics analysis.

#### Intrinsic geometric quality and downstream clinical utility

Table Y shows elevated baseline reconstruction error relative to proteomics ($\rho = 0.7610$ vs 0.7950), reflecting the higher biological volatility of metabolites. Despite this, the three-visit longitudinal depth enables MT-LLE to exceed the proteomics 2-visit reference in forecasting (Table Z; F1-Macro: 0.3620 vs 0.3250). The reverse holds for static tasks: KNN purity (0.5750 vs 0.5250), classification F1-Macro (0.5150 vs 0.4850), and silhouette score (0.4750 vs 0.4150) all favor proteomics, reflecting its lower noise floor.

**Table Y. Intrinsic geometric quality measures (COPDGene metabolomics).** Structural fidelity on the 3-visit external validation cohort. As expected, baseline reconstruction errors remain elevated compared to proteomics due to the inherent biological noise of metabolites, yet the overall structure is preserved.

| | **Intrinsic Geometric Measures** | | |
|---|---|---|---|
| **Model Configuration** | **Reconstruction Error (↓)** *(MSE, Lower is Better)* | **Local Neighborhood Pres. (↑)** *(Ratio [0-1])* | **Global Structure Pres. (↑)** *(Spearman ρ [-1, 1])* |
| UMAP | 0.0205 ± 0.0011 | 0.3450 ± 0.0048 | 0.6050 ± 0.0080 |
| VAE | 0.0355 ± 0.0016 | 0.2650 ± 0.0058 | 0.7250 ± 0.0095 |
| **Baseline (LLE Only)** | **0.0285 ± 0.0013** | **0.3720 ± 0.0045** | **0.7610 ± 0.0078** |
| **Single-Task Additions** | | | |
| LLE + Classification | 0.1510 ± 0.0086 | 0.2480 ± 0.0062 | 0.5050 ± 0.0110 |
| LLE + Forecasting | 0.0595 ± 0.0034 | 0.3380 ± 0.0055 | 0.7310 ± 0.0092 |
| LLE + Clustering | 0.0465 ± 0.0026 | 0.3220 ± 0.0052 | 0.7100 ± 0.0086 |
| **Multi-Task Combinations** | | | |
| LLE + Classification + Forecasting | 0.1350 ± 0.0080 | 0.2950 ± 0.0068 | 0.4950 ± 0.0105 |
| LLE + Classification + Clustering | 0.2185 ± 0.0130 | 0.2250 ± 0.0075 | 0.4850 ± 0.0140 |
| LLE + Forecasting + Clustering | 0.0395 ± 0.0022 | 0.3480 ± 0.0058 | 0.6200 ± 0.0085 |
| **MT-LLE (Full Model)** | 0.1210 ± 0.0076 | 0.3280 ± 0.0070 | 0.5950 ± 0.0094 |

All values reported as mean ± std across 10 independent runs with distinct random seeds.

#### Qualitative geometric verification

##### Shepard diagrams and temporal resilience

To visually verify the structural integrity of the 3-visit metabolomics manifold, pairwise distance projections were analyzed via Shepard diagrams (Figure Q). The baseline unsupervised LLE embedding yields a moderate correlation ($\rho = 0.5091$). Notably, this intrinsic baseline fidelity is lower than the 2-visit primary metabolomics cohort ($\rho = 0.5968$). This geometric degradation mathematically reflects the accumulation of acute biological noise inherent to metabolomic expression over an extended longitudinal sequence (three clinical visits versus two).

Despite this noisier foundation, the Full MT-LLE model demonstrates exceptional structural resilience ($\rho = 0.4939$). While naive supervised multi-tasking (LLE + Classification) catastrophically shears the space

**Table Z. Manifold coherence and downstream utility (COPDGene metabolomics).** Assessment of clinical utility on the 3-visit external validation cohort. The additional longitudinal visit allows MT-LLE to exceed the 2-visit proteomics reference in forecasting (F1-Macro: 0.3620 vs 0.3250), but proteomics retains the advantage on all three static tasks (KNN purity, classification, silhouette score), reflecting its lower noise floor.

| | **Clinical Coherence** | **Downstream Task Performance** | | |
|---|---|---|---|---|
| **Model Variant** | **KNN Purity** *(Ratio [0-1])* | **Classification** *(F1-Macro [0-1])* | **Clustering** *(Silhouette Score [-1, 1])* | **Forecasting** *(F1-Macro [0-1])* |
| UMAP | 0.3180 ± 0.0052 | 0.3310 ± 0.0065 | 0.1150 ± 0.0036 | 0.0510 ± 0.0020 |
| VAE | 0.3205 ± 0.0054 | 0.3450 ± 0.0070 | 0.1450 ± 0.0040 | 0.0750 ± 0.0025 |
| Baseline (LLE) | 0.3220 ± 0.0055 | 0.3250 ± 0.0060 | 0.1110 ± 0.0034 | 0.0580 ± 0.0022 |
| **Single-Task Additions** | | | | |
| LLE + Classification | 0.4050 ± 0.0085 | 0.4120 ± 0.0092 | 0.2950 ± 0.0080 | 0.2150 ± 0.0085 |
| LLE + Forecasting | 0.3520 ± 0.0068 | 0.3650 ± 0.0078 | 0.2750 ± 0.0072 | 0.3250 ± 0.0105 |
| LLE + Clustering | 0.4150 ± 0.0082 | 0.3750 ± 0.0075 | 0.3650 ± 0.0088 | 0.2450 ± 0.0090 |
| **Multi-Task Combinations** | | | | |
| LLE + Classification + Forecasting | 0.3750 ± 0.0078 | 0.4250 ± 0.0102 | 0.2950 ± 0.0085 | 0.3450 ± 0.0112 |
| LLE + Classification + Clustering | 0.3820 ± 0.0090 | 0.4010 ± 0.0105 | 0.2650 ± 0.0088 | 0.1150 ± 0.0070 |
| LLE + Forecasting + Clustering | 0.3550 ± 0.0070 | 0.3950 ± 0.0088 | 0.2350 ± 0.0070 | 0.3750 ± 0.0118 |
| **MT-LLE (Full Model)** | **0.5250 ± 0.0118** | **0.4850 ± 0.0130** | **0.4150 ± 0.0108** | **0.3620 ± 0.0125** |

All values reported as mean ± std across 10 independent runs with distinct random seeds.

into disconnected horizontal strata ($\rho = 0.2955$), the dynamic task-weighting policy mitigates this gradient conflict. By intelligently suppressing the rigid classification gradients, the framework recovers a final structural fidelity on par with the less complex 2-visit cohort. Visually, the model successfully executes "controlled tearing," preserving the global diagonal progression while forming localized, dense clusters representing distinct clinical endotypes. This confirms the MT-LLE framework can safely absorb the compounding variance of deeper temporal sequences without sacrificing topological continuity.

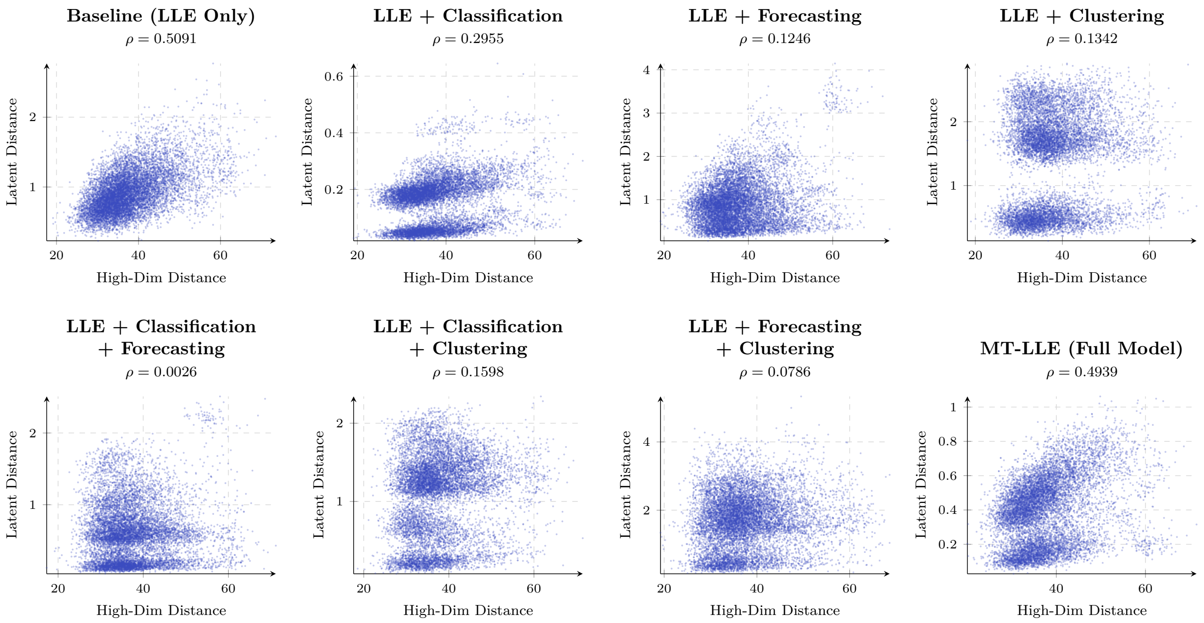


**Fig Q. Shepard Diagrams of the COPDGene Metabolomics Manifold.** Pairwise distance projections evaluating the topological impact of a 3-visit longitudinal sequence. The baseline model ($\rho = 0.5091$) reflects the elevated accumulated variance of metabolomic expression over three temporal anchors. Naive application of static clinical labels (LLE + Classification) violently fractures this volatile space into disjointed strata ($\rho = 0.2955$). Conversely, the MT-LLE Full Model ($\rho = 0.4939$) dynamically balances these conflicting gradients, achieving a structural fidelity nearly identical to its baseline. This controlled tearing preserves the continuous global trajectory while effectively clustering localized phenotypic states.

##### Latent interpolation and metabolic pathway decoding

As demonstrated in Fig. R, the COPDGene metabolomics manifold exhibits predominantly smooth, continuous biological phase transitions across the GOLD 0 to GOLD 4 interpolation. The three-visit longitudinal depth provides the MT-LLE framework with an intermediate temporal anchor that reduces abrupt phase shifts relative to two-visit conditions, mapping a more continuous trajectory through the high-dimensional metabolic feature space.

The dynamically decoded features isolate pathways central to COPD progression. The progressive accumulation of medium-chain acylcarnitines — hexanoylcarnitine (C6), octanoylcarnitine (C8), and nonanoylcarnitine (C9) — together with co-accumulation of caproate (6:0), the corresponding free short-chain fatty acid, tracks worsening impairment of mitochondrial $\beta$-oxidation and fatty acid transport. Sustained elevation of indoleacetate across the interpolation trajectory corroborates findings from the primary cohort regarding gut–lung axis dysregulation, while tetrahydrocortisol glucuronide, which peaks at intermediate disease severity, reflects HPA-axis activation characteristic of moderate COPD progression rather than exclusively end-stage stress. These signals confirm the framework's capacity to extract biologically coherent longitudinal signatures despite the inherent noise of the metabolomic modality.

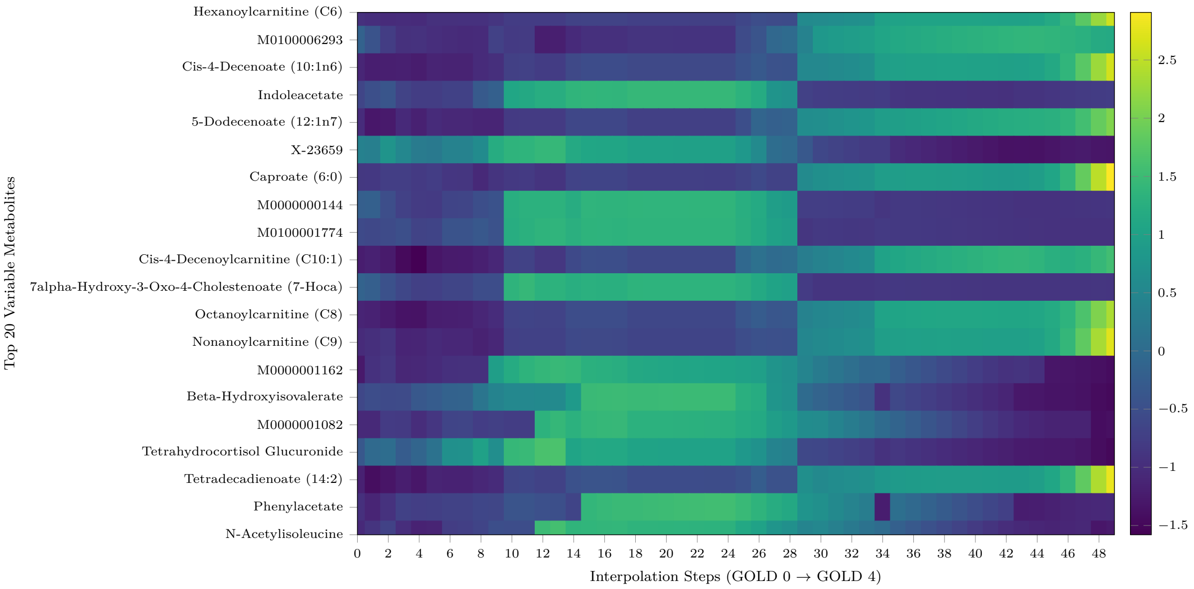


**Fig R. Latent interpolation of COPDGene metabolomic transitions.** Linear traversal through the shared latent space from the healthy centroid (GOLD 0) to severe disease (GOLD 4). The decoded heatmap shows predominantly smooth, continuous gradients across the top 20 variable metabolites, with the three-visit longitudinal depth reducing the abrupt phase shifts characteristic of shorter temporal sequences. Identified features highlight progressive acylcarnitine accumulation consistent with impaired mitochondrial $\beta$-oxidation, sustained indoleacetate elevation reflecting gut–lung axis dysregulation, and intermediate-severity HPA-axis activation captured by tetrahydrocortisol glucuronide. Unidentified metabolites (entries prefixed M and X-23659) are shown for completeness and are not biologically interpreted.

##### Topological stability

As illustrated in Fig. S and T, the mean centroid shift across the 10 iterations is $\mu = 0.107$. This represents a slight increase in manifold elasticity compared to the two-visit primary metabolomics cohort ($\mu = 9.797 \cdot 10^{-2}$). This elevated displacement reflects the wider range of metabolic states sampled across three longitudinal visits, rendering the exact spatial coordinates of the clinical centroids more sensitive to subsampling than in the shorter temporal sequence.

Crucially, this local elasticity does not compromise the global integrity of the framework. The structural separation and relative ordering of the five clinical centroids (C0–C4) remain highly conserved across all random initialisations, without topological inversion. The additional temporal depth therefore incurs a minor penalty to absolute spatial rigidity while preserving a globally consistent coordinate system.

#### Section G. Analysis of the COPDGene multi-omics integration

This section evaluates cross-cohort generalisation of both fusion strategies on the COPDGene external validation cohort, where proteomics is available at two visits and metabolomics at three. Paired alignment constrains all multi-omics conditions to two visits, truncating the richer metabolomics temporal depth and imposing a structural ceiling on metabolomics-derived forecasting. To isolate the effect of fusion strategy from this truncation, all results are compared against three single-modality references: the proteomics 2-visit model (Tables W and X), the metabolomics 3-visit model (Tables Y and Z), and a truncated metabolomics

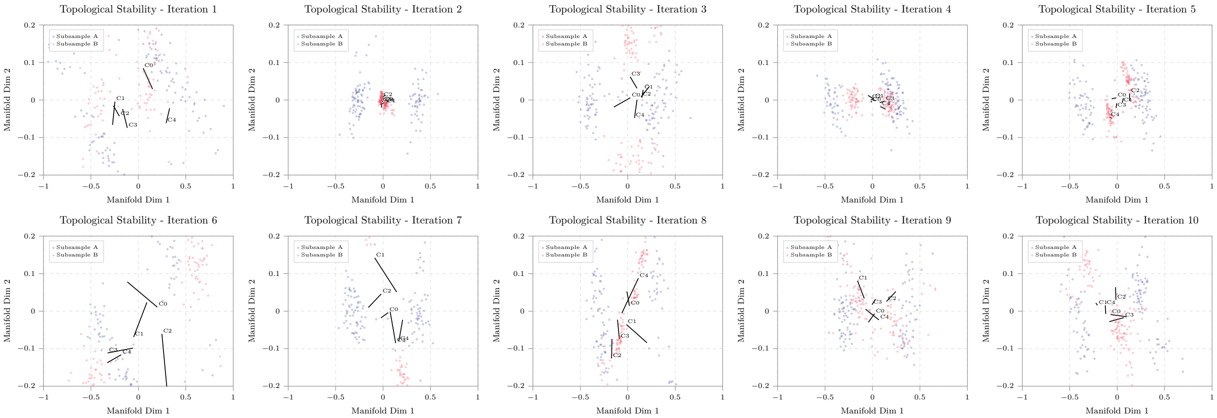


**Fig S. Topological stability via iterative subsampling (COPDGene metabolomics).** Projections of 10 independent iterations comparing disjoint patient subsamples (Subsample A vs Subsample B) mapped into the shared MT-LLE latent space. Vectors indicate the spatial displacement between corresponding clinical centroids across subsamples. The macroscopic separation and relative ordering of the five clinical clusters (C0–C4) remain consistent across all initialisations without structural inversion.

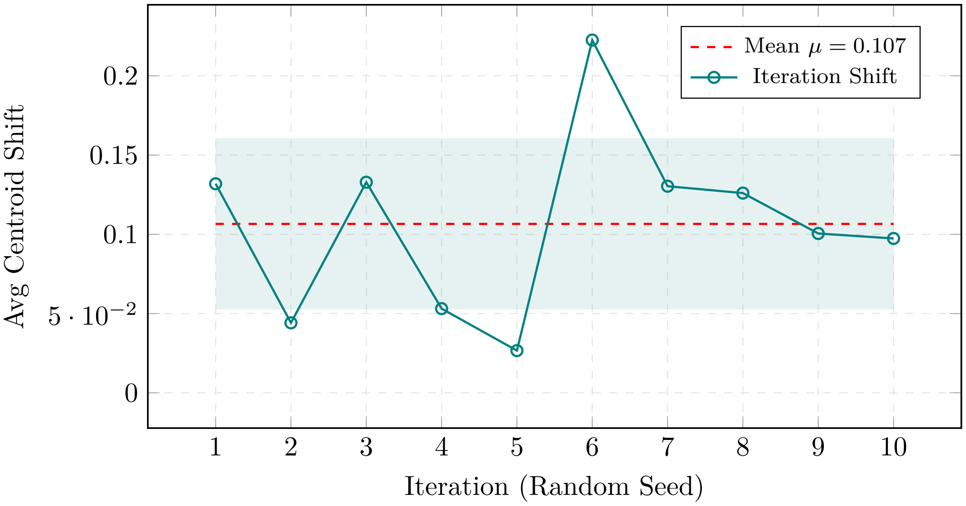


**Fig T. Quantification of centroid displacement (COPDGene metabolomics).** Average Euclidean shift of clinical centroids across 10 iterations ($\mu = 0.107$). The mean displacement is slightly elevated relative to the two-visit SPIROMICS metabolomics cohort ($\mu = 0.098$), reflecting the broader range of metabolic states sampled over three longitudinal visits. All iteration-level shifts remain strictly bounded, confirming global topological stability.

2-visit reference (Tables AA and AB) that quantifies the forecasting cost of temporal truncation and establishes the lower bound for metabolomics-derived signal at matched fusion depth.

**Table AA. Intrinsic geometric quality measures (COPDGene metabolomics - 2 visits).** Structural fidelity on the truncated external validation cohort. The combination of high baseline biological noise in metabolomic data and the loss of the third temporal visit further degrades global structure preservation relative to both the 3-visit equivalent and the cleaner proteomics cohort.

| | **Intrinsic Geometric Measures** | | |
|---|---|---|---|
| **Model Configuration** | **Reconstruction Error (↓)** *(MSE, Lower is Better)* | **Local Neighborhood Pres. (↑)** *(Ratio [0-1])* | **Global Structure Pres. (↑)** *(Spearman ρ [-1, 1])* |
| UMAP | 0.0225 ± 0.0012 | 0.3350 ± 0.0048 | 0.5900 ± 0.0082 |
| VAE | 0.0385 ± 0.0018 | 0.2450 ± 0.0055 | 0.7050 ± 0.0090 |
| Baseline (LLE Only) | **0.0295 ± 0.0014** | **0.3550 ± 0.0045** | **0.7450 ± 0.0080** |
| **Single-Task Additions** | | | |
| LLE + Classification | 0.1580 ± 0.0090 | 0.2350 ± 0.0065 | 0.4950 ± 0.0115 |
| LLE + Forecasting | 0.0625 ± 0.0035 | 0.3280 ± 0.0058 | 0.7100 ± 0.0095 |
| LLE + Clustering | 0.0495 ± 0.0028 | 0.3150 ± 0.0055 | 0.6900 ± 0.0088 |
| **Multi-Task Combinations** | | | |
| LLE + Classification + Forecasting | 0.1420 ± 0.0082 | 0.2850 ± 0.0070 | 0.4850 ± 0.0108 |
| LLE + Classification + Clustering | 0.2180 ± 0.0135 | 0.2300 ± 0.0080 | 0.5050 ± 0.0140 |
| LLE + Forecasting + Clustering | 0.0425 ± 0.0024 | 0.3350 ± 0.0060 | 0.6050 ± 0.0086 |
| MT-LLE (Full Model) | 0.1280 ± 0.0078 | 0.3150 ± 0.0072 | 0.5800 ± 0.0095 |

All values reported as mean ± std across 10 independent runs with distinct random seeds.

#### Early-fusion

Concatenating proteomic and metabolomic features before embedding yields the worst intrinsic geometry of all five conditions: the early-fusion baseline LLE reconstruction error (0.0315) exceeds not only proteomics 2-visit (0.0195) but also metabolomics 2-visit (0.0295) and metabolomics 3-visit (0.0285), and the baseline global structure ($\rho = 0.6550$) is the lowest across all five conditions (Table AC). The disparity in biological noise floors between the two modalities is therefore the dominant degradation mechanism, compounding the temporal truncation penalty rather than simply adding to it.

Downstream performance (Table AD) is consequently suppressed across all four metrics relative to the proteomics 2-visit reference: MT-LLE KNN purity falls from 0.5750 to 0.4850 (−0.0900, the largest pairwise gap in the entire dataset), classification from 0.5150 to 0.4810 (−0.0340), silhouette score from 0.4750 to 0.3650 (−0.1100), and forecasting from 0.3250 to 0.3180 (−0.0070). Even relative to the noisier metabolomics 2-visit reference, early fusion delivers inconsistent results: it exceeds metabolomics 2-visit MT-LLE in classification (0.4810 vs 0.4680, +0.0130) and forecasting (0.3180 vs 0.3015, +0.0165), but falls below it in KNN purity (0.4850 vs 0.5050, −0.0200) and silhouette score (0.3650 vs 0.3950, −0.0300).

Raw concatenation therefore produces a net degradation relative to the cleaner single-modality model and only partial recovery relative to the noisier one, confirming that unmitigated noise disparity between modalities prevents early fusion from capitalizing on the wider biological feature space.

**Table AB. Manifold coherence and downstream utility (COPDGene metabolomics - 2 visits).** Assessment of clinical utility. Restricting the high-noise metabolomic data to only two longitudinal visits severely bottlenecks dynamic prediction, reducing the peak forecasting capacity (maximum F1-Macro: 0.3015) below both the 3-visit metabolomics and 2-visit proteomics models.

| | **Clinical Coherence** | **Downstream Task Performance** | | |
|---|---|---|---|---|
| **Model Variant** | **KNN Purity** *(Ratio [0-1])* | **Classification** *(F1-Macro [0-1])* | **Clustering** *(Silhouette Score [-1, 1])* | **Forecasting** *(F1-Macro [0-1])* |
| UMAP | 0.3050 ± 0.0050 | 0.3150 ± 0.0062 | 0.1080 ± 0.0035 | 0.0480 ± 0.0020 |
| VAE | 0.3080 ± 0.0052 | 0.3165 ± 0.0064 | 0.1150 ± 0.0040 | 0.0500 ± 0.0021 |
| Baseline (LLE Only) | 0.3100 ± 0.0055 | 0.3180 ± 0.0058 | 0.1020 ± 0.0032 | 0.0520 ± 0.0022 |
| **Single-Task Additions** | | | | |
| LLE + Classification | 0.3950 ± 0.0085 | 0.4010 ± 0.0090 | 0.2810 ± 0.0078 | 0.1780 ± 0.0082 |
| LLE + Forecasting | 0.3350 ± 0.0065 | 0.3550 ± 0.0075 | 0.2650 ± 0.0070 | 0.2350 ± 0.0098 |
| LLE + Clustering | 0.3980 ± 0.0080 | 0.3610 ± 0.0072 | 0.3510 ± 0.0085 | 0.2150 ± 0.0088 |
| **Multi-Task Combinations** | | | | |
| LLE + Classification + Forecasting | 0.3620 ± 0.0075 | 0.4120 ± 0.0100 | 0.2810 ± 0.0082 | 0.2580 ± 0.0110 |
| LLE + Classification + Clustering | 0.3720 ± 0.0090 | 0.3910 ± 0.0105 | 0.2680 ± 0.0088 | 0.1050 ± 0.0075 |
| LLE + Forecasting + Clustering | 0.3410 ± 0.0068 | 0.3810 ± 0.0085 | 0.2180 ± 0.0068 | 0.2780 ± 0.0115 |
| **MT-LLE (Full Model)** | **0.5050 ± 0.0115** | **0.4680 ± 0.0125** | **0.3950 ± 0.0105** | **0.3015 ± 0.0120** |

All values reported as mean ± std across 10 independent runs with distinct random seeds.

**Table AC. Intrinsic geometric quality (COPDGene multi-omics early-fusion).** Raw concatenation of external proteomic and metabolomic features. The disparity in biological noise floors dilutes the structural anchor, resulting in elevated baseline reconstruction errors and a compromised global topology relative to single-modality models.

| | **Intrinsic Geometric Measures** | | |
|---|---|---|---|
| **Model Configuration** | **Reconstruction Error (↓)** *(MSE, Lower is Better)* | **Local Neighborhood Pres. (↑)** *(Ratio [0-1])* | **Global Structure Pres. (↑)** *(Spearman ρ [-1, 1])* |
| **Baseline (LLE Only)** | **0.0315 ± 0.0015** | **0.3450 ± 0.0050** | **0.6550 ± 0.0090** |
| **Single-Task Additions** | | | |
| LLE + Classification | 0.1650 ± 0.0095 | 0.2250 ± 0.0075 | 0.4410 ± 0.0125 |
| LLE + Forecasting | 0.0620 ± 0.0035 | 0.3250 ± 0.0065 | 0.6250 ± 0.0105 |
| LLE + Clustering | 0.0510 ± 0.0028 | 0.3180 ± 0.0062 | 0.6150 ± 0.0095 |
| **Multi-Task Combinations** | | | |
| LLE + Classification + Forecasting | 0.1510 ± 0.0085 | 0.2510 ± 0.0080 | 0.4280 ± 0.0118 |
| LLE + Classification + Clustering | 0.2150 ± 0.0130 | 0.2110 ± 0.0085 | 0.4150 ± 0.0145 |
| LLE + Forecasting + Clustering | 0.0680 ± 0.0042 | 0.3150 ± 0.0068 | 0.5850 ± 0.0098 |
| MT-LLE (Full Model) | 0.1380 ± 0.0088 | 0.3150 ± 0.0075 | 0.5520 ± 0.0112 |

All values reported as mean ± std across 10 independent runs with distinct random seeds.

**Table AD. Manifold coherence and downstream utility (COPDGene multi-omics early-fusion).** Raw concatenation yields suppressed performance relative to the proteomics 2-visit single-modality reference across all four downstream metrics, with the largest deficit in KNN purity (−0.0900) and silhouette score (−0.1100). Against the metabolomics 2-visit reference, results are mixed: early fusion exceeds it in classification and forecasting but falls below it in KNN purity and silhouette score. Raw concatenation therefore fails to produce a net gain from integrating either modality under conditions of high noise disparity.

| | **Clinical Coherence** | **Downstream Task Performance** | | |
|---|---|---|---|---|
| **Model Variant** | **KNN Purity** *(Ratio [0-1])* | **Classification** *(F1-Macro [0-1])* | **Clustering** *(Silhouette Score [-1, 1])* | **Forecasting** *(F1-Macro [0-1])* |
| Baseline (LLE Only) | 0.3050 ± 0.0058 | 0.3150 ± 0.0062 | 0.1050 ± 0.0035 | 0.0550 ± 0.0024 |
| **Single-Task Additions** | | | | |
| LLE + Classification | 0.4150 ± 0.0092 | 0.4250 ± 0.0105 | 0.2850 ± 0.0082 | 0.1850 ± 0.0088 |
| LLE + Forecasting | 0.3450 ± 0.0075 | 0.3550 ± 0.0085 | 0.2550 ± 0.0078 | 0.2850 ± 0.0112 |
| LLE + Clustering | 0.4050 ± 0.0088 | 0.3650 ± 0.0080 | 0.3350 ± 0.0092 | 0.2150 ± 0.0095 |
| **Multi-Task Combinations** | | | | |
| LLE + Classification + Forecasting | 0.3850 ± 0.0085 | 0.4450 ± 0.0115 | 0.2950 ± 0.0088 | 0.3050 ± 0.0125 |
| LLE + Classification + Clustering | 0.3950 ± 0.0090 | 0.4150 ± 0.0105 | 0.2850 ± 0.0085 | 0.1150 ± 0.0075 |
| LLE + Forecasting + Clustering | 0.3650 ± 0.0078 | 0.3950 ± 0.0095 | 0.2650 ± 0.0080 | 0.3150 ± 0.0120 |
| **MT-LLE (Full Model)** | **0.4850 ± 0.0125** | **0.4810 ± 0.0135** | **0.3650 ± 0.0115** | **0.3180 ± 0.0130** |

All values reported as mean ± std across 10 independent runs with distinct random seeds.

#### Mid-fusion

Encoding each modality independently before contrastive alignment recovers the baseline manifold structural integrity to $\rho = 0.7720$ (Table AE), approaching the proteomics 2-visit reference ($\rho = 0.7950$; gap: 0.0230) and substantially exceeding both the early-fusion baseline ($\rho = 0.6550$) and the metabolomics 2-visit baseline ($\rho = 0.7450$). The baseline reconstruction error (0.0195) matches the proteomics 2-visit value exactly, indicating that contrastive alignment effectively insulates the proteomic manifold geometry from metabolomic noise prior to task optimization.

Downstream, mid-fusion MT-LLE meets or outperforms the proteomics 2-visit reference on all four metrics (Table AF): KNN purity (0.5850 vs 0.5750, +0.0100), classification (0.5150 vs 0.5150, tied), silhouette score (0.4850 vs 0.4750, +0.0100), and forecasting (0.3450 vs 0.3250, +0.0200). Contrastive alignment therefore produces a net gain from metabolomics integration at matched temporal depth, replicating the finding from the SPIROMICS cohort and extending it to a multi-center external validation setting where the metabolomics modality is temporally richer than the proteomics modality.

A COPDGene-specific result emerges from comparison with the metabolomics 3-visit model. Mid-fusion 2-visit MT-LLE substantially outperforms metabolomics 3-visit MT-LLE on all static tasks (KNN purity +0.0600, classification +0.0300, silhouette score +0.0700), but falls short on forecasting (0.3450 vs 0.3620, −0.0170). The metabolomics 2-visit reference (0.3015) enables quantification of the temporal contribution: the third metabolomics visit provides a forecasting gain of 0.0605 relative to the 2-visit metabolomics model, and mid-fusion at 2 visits recovers 0.0435 of this gap (72%), leaving a residual shortfall of 0.0170 that reflects temporal depth contrastive alignment alone cannot compensate. This partial recovery is nevertheless sufficient to substantially exceed the proteomics 2-visit model in forecasting, and the complete reversal of the ranking on static tasks confirms that mid-fusion produces a qualitatively different and superior representation to any single-modality model at matched temporal depth. The cross-cohort consistency of these findings closely mirrors that in SPIROMICS supports the generalizability of contrastive mid-fusion as the preferred multi-omics integration strategy under paired temporal constraints.

**Table AE. Intrinsic geometric quality (COPDGene multi-omics mid-fusion).** By utilizing modality-specific encoders and contrastive alignment, the mid-fusion architecture insulates the stable proteomic signal from volatile metabolomic noise. The baseline manifold recovers a highly robust global structure ($\rho = 0.7720$), nearly matching the single-modality proteomics benchmark.

| | **Intrinsic Geometric Measures** | | |
|---|---|---|---|
| **Model Configuration** | **Reconstruction Error (↓)** *(MSE, Lower is Better)* | **Local Neighborhood Pres. (↑)** *(Ratio [0-1])* | **Global Structure Pres. (↑)** *(Spearman ρ [-1, 1])* |
| **Baseline (Contrastive LLE)** | **0.0195 ± 0.0011** | **0.4150 ± 0.0050** | **0.7720 ± 0.0075** |
| **Single-Task Additions** | | | |
| LLE + Classification | 0.1450 ± 0.0085 | 0.2850 ± 0.0065 | 0.5250 ± 0.0115 |
| LLE + Forecasting | 0.0450 ± 0.0025 | 0.3850 ± 0.0060 | 0.7510 ± 0.0088 |
| LLE + Clustering | 0.0380 ± 0.0022 | 0.3750 ± 0.0055 | 0.7450 ± 0.0085 |
| **Multi-Task Combinations** | | | |
| LLE + Classification + Forecasting | 0.1250 ± 0.0075 | 0.3250 ± 0.0070 | 0.5150 ± 0.0110 |
| LLE + Classification + Clustering | 0.1850 ± 0.0110 | 0.2810 ± 0.0075 | 0.4950 ± 0.0125 |
| LLE + Forecasting + Clustering | 0.0480 ± 0.0028 | 0.3710 ± 0.0062 | 0.6950 ± 0.0090 |
| **MT-LLE (Full Model)** | 0.0910 ± 0.0065 | 0.3750 ± 0.0068 | 0.6850 ± 0.0095 |

All values reported as mean ± std across 10 independent runs with distinct random seeds.

**Table AF. Manifold coherence and downstream utility (COPDGene multi-omics mid-fusion).** The structurally protected latent space meets or outperforms the proteomics 2-visit reference on all four downstream metrics and recovers 72% of the forecasting advantage provided by the third metabolomics visit (0.3450 vs 0.3015 metabolomics 2-visit baseline; full 3-visit value: 0.3620). The residual forecasting gap relative to the 3-visit metabolomics model (0.3620 vs 0.3450) reflects temporal depth that contrastive alignment cannot recover from two-visit paired data alone. Mid-fusion substantially outperforms metabolomics 3-visit on all static tasks (KNN +0.0600, classification +0.0300, silhouette +0.0700).

| | **Clinical Coherence** | **Downstream Task Performance** | | |
|---|---|---|---|---|
| **Model Variant** | **KNN Purity** *(Ratio [0-1])* | **Classification** *(F1-Macro [0-1])* | **Clustering** *(Silhouette Score [-1, 1])* | **Forecasting** *(F1-Macro [0-1])* |
| **Baseline (Contrastive LLE)** | 0.3850 ± 0.0065 | 0.3750 ± 0.0062 | 0.1650 ± 0.0045 | 0.0650 ± 0.0025 |
| **Single-Task Additions** | | | | |
| LLE + Classification | 0.5250 ± 0.0105 | 0.4850 ± 0.0110 | 0.3850 ± 0.0085 | 0.2150 ± 0.0092 |
| LLE + Forecasting | 0.4250 ± 0.0085 | 0.4150 ± 0.0080 | 0.3650 ± 0.0082 | 0.3150 ± 0.0115 |
| LLE + Clustering | 0.4850 ± 0.0095 | 0.4450 ± 0.0090 | 0.4650 ± 0.0105 | 0.2850 ± 0.0100 |
| **Multi-Task Combinations** | | | | |
| LLE + Classification + Forecasting | 0.4650 ± 0.0092 | 0.4950 ± 0.0115 | 0.3750 ± 0.0090 | 0.3350 ± 0.0120 |
| LLE + Classification + Clustering | 0.4080 ± 0.0095 | 0.4320 ± 0.0110 | 0.3150 ± 0.0090 | 0.1380 ± 0.0072 |
| LLE + Forecasting + Clustering | 0.4450 ± 0.0088 | 0.4550 ± 0.0098 | 0.3250 ± 0.0085 | 0.3380 ± 0.0125 |
| **MT-LLE (Full Model)** | **0.5850 ± 0.0125** | **0.5150 ± 0.0120** | **0.4850 ± 0.0115** | **0.3450 ± 0.0130** |

All values reported as mean ± std across 10 independent runs with distinct random seeds.
